# Immune cell type divergence in a basal chordate

**DOI:** 10.1101/2025.05.20.655184

**Authors:** Tal D. Scully, C. J. Pickett, Nicolas A. Gort-Freitas, Bradley Davidson, Allon M. Klein

**Author notes:** Corresponding author: Allon Klein.

## Abstract

Evolutionary adaptations often occur at the level of cell types and cellular function. Innate immune cells are a promising system for studying cell type evolution, as they are widespread across metazoans, have several conserved functions, and are under selective pressure from pathogens. However, molecular characterizations of invertebrate immune cells are limited, and it remains unclear whether invertebrate immune cell types are homologous to those in vertebrates. Here we use single-cell RNA sequencing, *in situ* hybridization, and live reporters to define the identity of blood cell states from a basal chordate, *Ciona robusta*. We find evidence that *C. robusta* circulating blood contains a differentiation hierarchy that gives rise to at least eight major morphotypes, constituting approximately half of mature blood cell states. The mature cell states include phagocytes, as well as cells variously expressing vanadium-binding proteins, carbonic anhydrases, pattern recognition receptors, cytokines, and complement factors. Despite the expression of homologs to vertebrate immune components, extensive divergence between tunicate and vertebrate immune cells obscures cell state homology. Altogether, this work modernizes blood cell classifications in *C. robusta* and extends the known repertoire of immune cells within chordates.

## Introduction

Evolutionary adaptations occur at different scales, from genes to body plans. Intermediate to these is the diversification of gene expression and cell types^1–3^. The emergence of novel cell types has facilitated major changes in organismal lifestyle and physiology, as observed with the emergence of vertebrate adaptive immune lymphoid cells and oxygen-transporting red blood cells^4^. New cell types can arise in conjunction with the emergence of novel proteins^2,5^, like red blood cells and hemoglobin^6^, or from changes in the expression of existing proteins^3,7^.

A tissue of practical interest for studying cellular diversification is the blood and its constituent immune cells. Immune cells are broadly present across metazoans^8,9^, with widely conserved cellular functions that include pathogen recognition, phagocytosis, and cytotoxicity^10^. Immune systems affect ecosystem dynamics^11^ and can serve as a source of antimicrobials^12^. They are also under selective pressure from rapidly evolving pathogens^13^, potentially making immune systems a laboratory for rapid evolution of cellular sensing and memory^14,15^.

Our understanding of immune cell types is largely drawn from studies of vertebrates^16^. Less is known about invertebrate immune cells, and much of what we do know is focused on features shared with vertebrates—including homologous pattern recognition receptors (PRRs)^15,17,18^, cytokines^19,20^, and complement factors^21–23;^ as well as shared phagocytic^24–26^ and cytotoxic^9,27,28^ cellular functions. Some immune proteins are notably diversified in invertebrate species, particularly PRRs^10,17^. At the level of cell states, it is unclear whether invertebrate and vertebrate immune cells are homologous. Certain immune cells in *Drosophila* and other invertebrates have been identified as macrophages based on shared phagocytic function^29–31^, but in vertebrates, macrophages are multifunctional, also playing a role in antigen presentation^32^. For most invertebrates, we lack the molecular information to fully assess cell state homology. Immune cells in most species are still largely classified by morphology^30,33,34^, which may conceal heterogeneous cell states. Our understanding of these cells would benefit from detailed molecular characterization.

We focus on the blood cells of tunicates, which shared a common ancestor with vertebrates ∼550 million years ago^35^. Tunicates sit at a key juncture in immune cell evolution because they are the most closely related living animals to vertebrates, and they lack adaptive immunity.

Tunicate immune cells mainly reside in the blood, which is pumped by a heart through a semi-open circulatory system^36,37^. Immune cells are morphologically diverse and are capable of phagocytosis, cytotoxic cell killing, foreign body encapsulation, allorecognition, and regeneration^38–41^. Like most invertebrates to date, their blood cells have been defined based on morphology, with limited molecular characterization^34,38^.

Here, we have generated a combined transcriptomic and morphological atlas of circulating blood cells from the well-studied tunicate species, *Ciona robusta* (**Fig. 1a**). We find a differentiation hierarchy in *C. robusta* blood, and confirm the identity of several progenitor and differentiated cell states *in vivo* by deployment of live cell reporters. We show that these blood cells are substantially more diverse than previous morphological classifications suggest. We begin to characterize several mature cell states, including cells expressing homologs of vertebrate immune effector genes. However, we find that *C. robusta* blood cells have diverged considerably from vertebrate blood cells, offering no clear evidence of the nature of ancestral immune cell types common to vertebrates and tunicates. Altogether, these observations extend a description of chordate immune cells beyond vertebrates and make clear the evolutionary plasticity of immune cell identity and gene expression.

**Figure 1:**
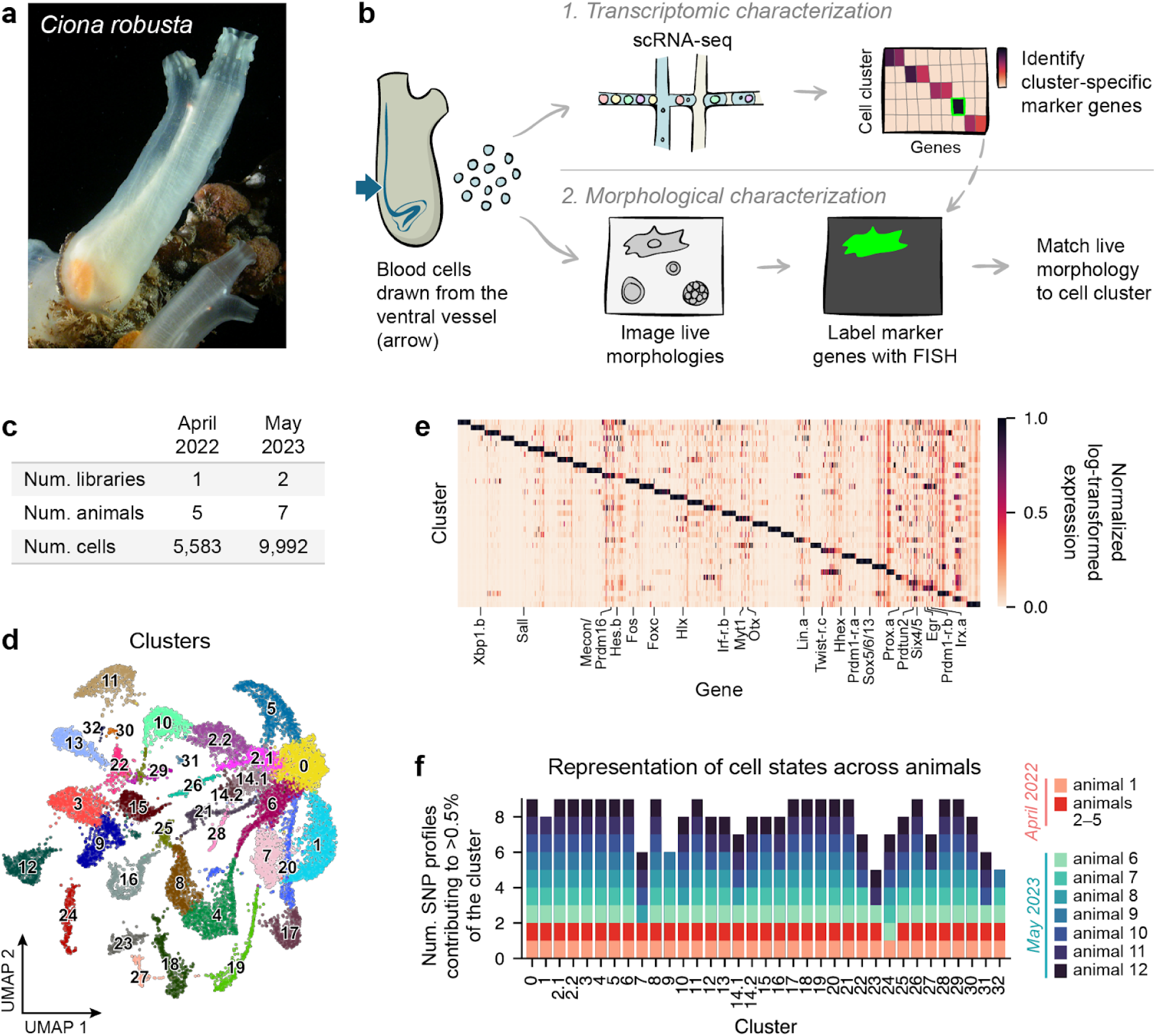
scRNA-seq reveals a complex set of transcriptional states in *C. robusta* blood. **(a)** Photo of adult *C. robusta* (formerly *C. intestinalis* type A^76^) from Queen Anne’s Battery Marina, Plymouth, UK. Image was kindly shared by John Bishop. **(b)** Approach for characterizing blood cells transcriptionally, then linking morphological information to transcriptional clusters using *in situ* hybridization. **(c)** Table showing, for each experimental day, the number of scRNA-seq libraries, the number of animals contributing blood, and the number of cells detected after removing low-quality cell barcodes. **(d)** UMAP of scRNA-seq data with cells colored and labeled by Leiden cluster. **(e)** Heatmap showing the top 20 enriched and cluster-specific genes for each Leiden cluster. Genes were chosen by identifying differentially expressed genes using the Wilcoxon rank-sum test, requiring a log2 fold change above 1 and a false discovery rate below 0.05. To enforce cluster specificity, we additionally required each gene to have a difference of log-transformed expression above 0.5 between its most highly expressed and second most highly expressed cluster. The 20 genes with the top highest log2 fold change are plotted. **(f)** For each Leiden cluster, the number of detected SNP profiles (with each profile corresponding to 1 or a few animals). A SNP profile is detected if it contributes to >0.5% of a cluster.

## Results

### A single-cell transcriptomic atlas of *C. robusta* blood

We set out to document the cells present in *C. robusta* blood through transcriptional profiling and imaging (**Fig. 1b**). This informally establishes a census of “cell types”, but definitions of a “cell type” are evasive and a matter of some debate^42,43^. To avoid ambiguity, we here use the term “morphotype” when referring to cells by their morphology and “cell state” when referring to gene expression state. We use “cell type” only when discussing conceptual theories related to cellular diversification, or when referring to cells in well-studied organisms which are widely referred to as distinct cell types (e.g. T and B cells).

Previously characterized *C. robusta* blood cell morphotypes are diverse^34,39,44,45^ and include small round cells that resemble lymphocytes, hyaline ameboid-like cells, a range of granule-containing cells, and cells with features that differ strongly from vertebrate cell morphologies—such as morula cells (MC) and compartment cells with several large vacuoles, as well as signet ring cells (SRC) and unilocular refractile granulocytes (URG) that each have a single large vacuole. However, classifications of tunicate blood cells vary, with studies defining six^39^, seven^36,45^, or nine^34,44,46^ morphotypes. Cellular nomenclature is not consistent, with over 30 different designations used across studies^34,39^.

To obtain a harmonized definition of cell states, we generated a transcriptomic atlas of *C. robusta* blood cells. We collected circulating blood cells from 12 adult animals over two separate days (**Fig. 1c**) and profiled them by single-cell RNA sequencing (scRNA-seq). Collection of this data required optimization to accommodate the high salinity of *C. robusta* blood^47^. We found that blood cells die when exposed to standard buffers for scRNA-seq, but cell viability can be rescued by a buffer containing mannitol, as we have reported in a separate technical paper^47^. In this way, we sampled 15,575 cells post-filter, with a median of 6,138 transcripts detected per cell.

We investigated the range of cell states in *C. robusta* blood by carrying out library batch correction, clustering, and UMAP visualization of the scRNA-seq data. The resulting embedding reveals that *C. robusta* blood contains a highly complex mixture of transcriptional states (**Fig. 1d,e**). We fractionated cell states to a resolution of 35 transcriptional clusters by Leiden clustering and then manually partitioning two heterogeneous clusters. All cell clusters are represented in data from both experimental days (**Fig. 1f**).

Since blood was sampled from wild-caught animals, there might be animal-specific differences in cell state resulting from differences in genotype, environment, symbionts, or infection. To evaluate whether the observed cell states are representative of all animals sampled, we took advantage of natural single nucleotide polymorphisms (SNPs) found within profiled mRNAs to assign cells to individual animals^48,49^. We distinguished nine SNP profiles: eight representing one animal each, and a ninth mixed SNP profile representing four animals with fewer cells sampled (**Supplementary Fig. 1a–c**). This analysis showed that 26 of the 35 clusters are represented by at least 8/9 SNP profiles, and four more are in 7/9 (**Fig. 1f**). The remaining clusters are represented by 5/9 or 6/9 SNP profiles and may represent environmentally-induced cell states. This SNP analysis also revealed that, for two pairs of clusters, animal-specific differences caused transcriptionally similar cell states to cluster separately (clusters 1/7 and 0/14.1; **Supplementary Fig. 1d–k**); in both cases, we combined the clusters, resulting in 33 total cell states. Thus, the number of common, transcriptionally distinct cell states in *C. robusta* blood is between 26 and 33—a substantially larger number than the 6–9 morphotypes in current classifications^34,36,39,44,45^.

### Linking transcriptional state to cell morphology

In vertebrates, the first scRNA-seq atlases of blood yielded transcriptional states that could immediately be associated with canonical cell types, owing to decades of work in purifying cell populations and identifying genes specific to each^50^. In tunicates, there is some limited prior information associating blood morphotypes with gene expression from *in situ* hybridization and immunostaining studies (see **Supplementary Table 1**). However, we found that these previously identified markers are insufficient to annotate clusters: several markers are not cluster specific (e.g. *CrGal-a*, *CrGal-b*), other markers which should label multiple morphotypes are expressed by only one cluster (e.g. *CiEM-a*), and many clusters do not express any previously proposed markers (**Supplementary Fig. 2a**). We therefore used microscopy to associate cell morphologies with expression of new sets of markers, which we derived from the scRNA-seq data (**Fig. 1b**).

To characterize cell morphology free from fixation artifacts (**Supplementary Fig. 2b**), we used differential interference contrast (DIC) imaging of live blood cells *ex vivo*. We then associated observed morphotypes with transcriptional cell states by fixing the cells in place with paraformaldehyde and labeling select genes using hybridization chain reaction fluorescent *in situ* hybridization (HCR FISH)^51^. We selected 1–2 genes to mark each cluster, along with a ubiquitously expressed reference gene (**Supplementary Fig. 2c,d**). We chose clusters that had distinct markers, including those that were not ubiquitous across animals (as documented in **Fig. 1f**). Cells were manually matched across multiple rounds of imaging, allowing association of live- and fixed-cell morphologies with gene expression (**Fig 2a**, **Supplementary Fig. 2e**).

**Figure 2:**
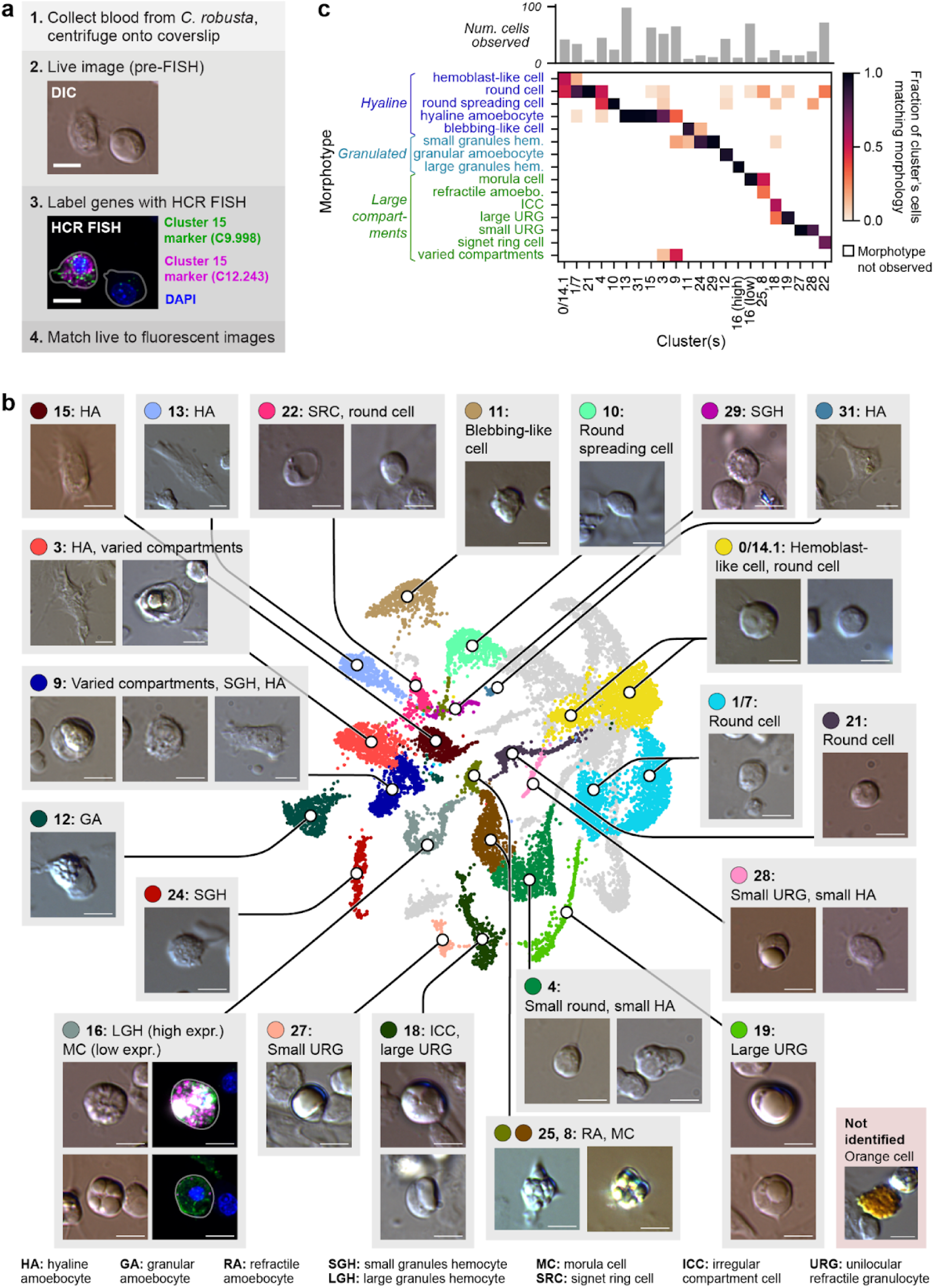
Linking transcriptional clusters to morphotype demonstrates limitations of previous morphological categories **(a)** Approach for identifying live morphologies of marker-positive cells. Example images are shown for Leiden cluster 15. Scale bars = 5 µm. **(b)** UMAP with Leiden clusters annotated by representative images and morphological descriptions. Fluorescence images are shown for cluster 16 to show high vs. low marker gene expression; see Supplementary Fig. 3 for each cluster’s fluorescence images. Scale bars = 5 µm. **(c)** Bottom: Heatmap showing the frequency of observed morphologies for each Leiden cluster labeled. Top: Bar graph indicating the total number of marker-positive cells counted for each Leiden cluster. 16-high and 16-low refer to cells expressing high or low levels of cluster 16 marker genes, respectively. Abbreviations: HA, hyaline amoebocyte; GA, granular amoebocyte; RA, refractile amoebocyte; SRC, signet ring cell; SGH, small granules hemocyte; LGH, large granules hemocyte; MC, moral cell; ICC, irregular compartment cell; URG, unilocular refractile granulocyte; Hem., hemocyte; Amoebo., amoebocyte.

Using this approach, we annotated 24 of the 33 clusters in the scRNA-seq data (**Fig. 2b**, **Supplementary Fig. 3**). We assigned a transcriptional identity to all previously documented tunicate blood morphotypes, with the exception of rare orange cells, which were observed but not labeled by any marker genes tested (**Fig. 2b**). As a result of these integrated measurements, we propose an updated joint cell state and morphotype classification of *C. robusta* blood, summarized in **Table 1** (see also **Supplementary Fig. 4a,b**, **Supplementary Table 2**). We propose names for each cell state, incorporating both classical morphotype names and functional annotations that will be justified in subsequent results sections. Where possible, we have aligned nomenclature with prior definitions^34,36,44,45^.

**Table 1:**
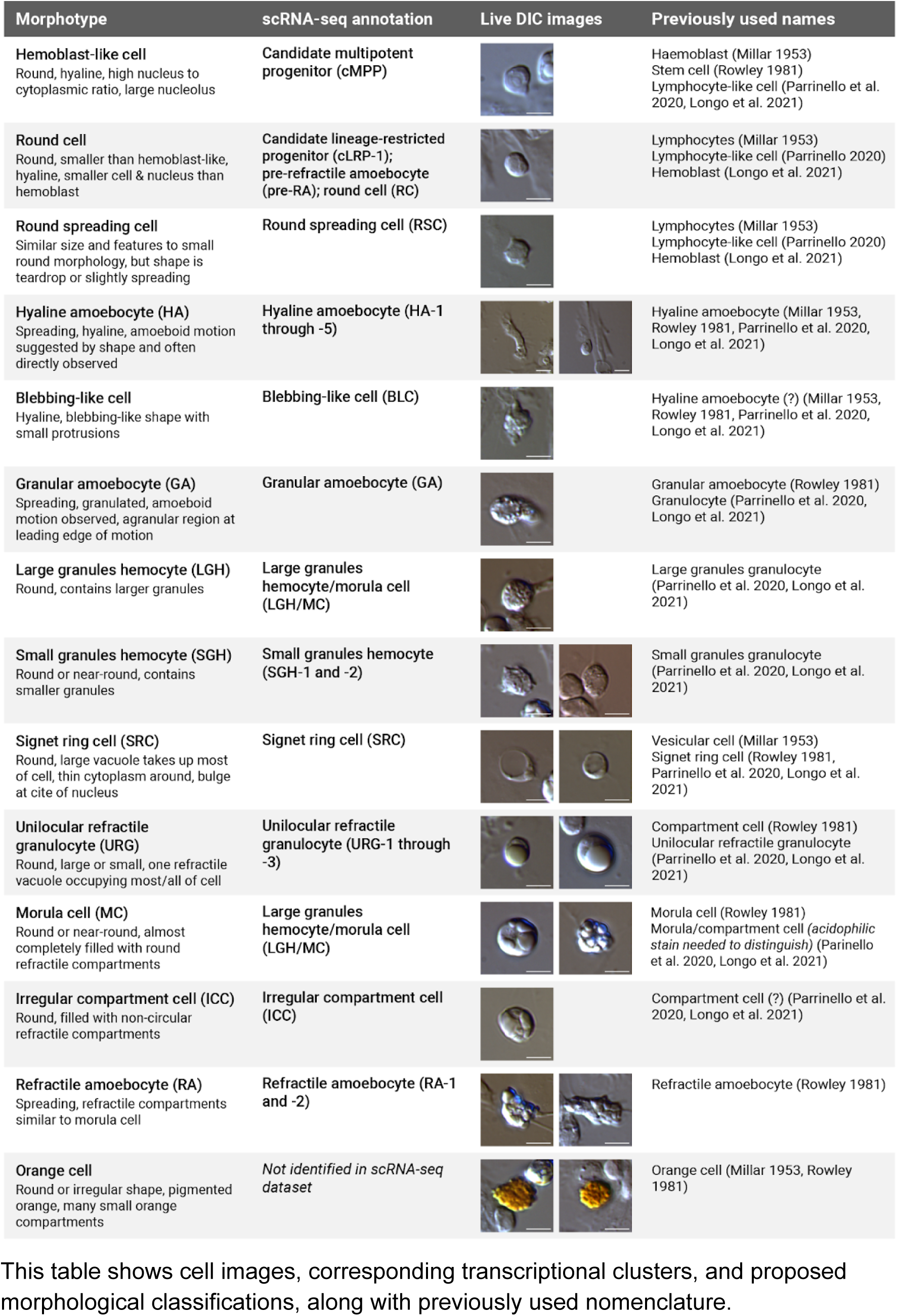
Proposed revision to *C. robusta* blood cell categorizations This table shows cell images, corresponding transcriptional clusters, and proposed morphological classifications, along with previously used nomenclature.

| Morphotype | scRNA-seq annotation | Live DIC images | Previously used names |
| --- | --- | --- | --- |
| **Hemoblast-like cell**<br>Round, hyaline, high nucleus to cytoplasmic ratio, large nucleolus | Candidate multipotent progenitor (cMPP) |  | Haemoblast (Millar 1953)<br>Stem cell (Rowley 1981)<br>Lymphocyte-like cell (Parrinello et al. 2020, Longo et al. 2021) |
| **Round cell**<br>Round, smaller than hemoblast-like, hyaline, smaller cell & nucleus than hemoblast | Candidate lineage-restricted progenitor (cLRP-1); pre-refractile amoebocyte (pre-RA); round cell (RC) |  | Lymphocytes (Millar 1953)<br>Lymphocyte-like cell (Parrinello 2020)<br>Hemoblast (Longo et al. 2021) |
| **Round spreading cell**<br>Similar size and features to small round morphology, but shape is teardrop or slightly spreading | Round spreading cell (RSC) |  | Lymphocytes (Millar 1953)<br>Lymphocyte-like cell (Parrinello 2020)<br>Hemoblast (Longo et al. 2021) |
| **Hyaline amoebocyte (HA)**<br>Spreading, hyaline, amoeboid motion suggested by shape and often directly observed | Hyaline amoebocyte (HA-1 through -5) |  | Hyaline amoebocyte (Millar 1953, Rowley 1981, Parrinello et al. 2020, Longo et al. 2021) |
| **Blebbing-like cell**<br>Hyaline, blebbing-like shape with small protrusions | Blebbing-like cell (BLC) |  | Hyaline amoebocyte (?) (Millar 1953, Rowley 1981, Parrinello et al. 2020, Longo et al. 2021) |
| **Granular amoebocyte (GA)**<br>Spreading, granulated, amoeboid motion observed, agranular region at leading edge of motion | Granular amoebocyte (GA) |  | Granular amoebocyte (Rowley 1981)<br>Granulocyte (Parrinello et al. 2020, Longo et al. 2021) |
| **Large granules hemocyte (LGH)**<br>Round, contains larger granules | Large granules hemocyte/morula cell (LGH/MC) |  | Large granules granulocyte (Parrinello et al. 2020, Longo et al. 2021) |
| **Small granules hemocyte (SGH)**<br>Round or near-round, contains smaller granules | Small granules hemocyte (SGH-1 and -2) |  | Small granules granulocyte (Parrinello et al. 2020, Longo et al. 2021) |
| **Signet ring cell (SRC)**<br>Round, large vacuole takes up most of cell, thin cytoplasm around, bulge at cite of nucleus | Signet ring cell (SRC) |  | Vesicular cell (Millar 1953)<br>Signet ring cell (Rowley 1981, Parrinello et al. 2020, Longo et al. 2021) |
| **Unilocular refractile granulocyte (URG)**<br>Round, large or small, one refractile vacuole occupying most/all of cell | Unilocular refractile granulocyte (URG-1 through -3) |  | Compartment cell (Rowley 1981)<br>Unilocular refractile granulocyte (Parrinello et al. 2020, Longo et al. 2021) |
| **Morula cell (MC)**<br>Round or near-round, almost completely filled with round refractile compartments | Large granules hemocyte/morula cell (LGH/MC) |  | Morula cell (Rowley 1981)<br>Morula/compartment cell (*acidophilic stain needed to distinguish*) (Parrinello et al. 2020, Longo et al. 2021) |
| **Irregular compartment cell (ICC)**<br>Round, filled with non-circular refractile compartments | Irregular compartment cell (ICC) |  | Compartment cell (?) (Parrinello et al. 2020, Longo et al. 2021) |
| **Refractile amoebocyte (RA)**<br>Spreading, refractile compartments similar to morula cell | Refractile amoebocyte (RA-1 and -2) |  | Refractile amoebocyte (Rowley 1981) |
| **Orange cell**<br>Round or irregular shape, pigmented orange, many small orange compartments | Not identified in scRNA-seq dataset |  | Orange cell (Millar 1953, Rowley 1981) |

**Table 1** and **Fig. 2** show several notable relationships between transcriptional state and morphotype. Several established morphotypes consist of more than one transcriptional cluster (**Fig. 2c**), with four clusters mapping to URGs (18, 19, 27, and 28), five to hyaline amoebocytes (HA) (3, 9, 13, 15, and 31), and five to small round cells (0/14.1, 1/7, 21, 4, 22). Despite their morphological similarity, the clusters vary in expression of hundreds of genes and express unique sets of transcription factors (**Supplementary Fig. 4c–e**). Conversely, ten transcriptional states each associate with more than one morphology (clusters 0/14, 3, 4, 8/25, 9, 16, 18, 19, 22, and 28; **Fig. 2b,c**). This morphological heterogeneity might align with transcriptional heterogeneity within clusters that capture a continuum of states. Within cluster 16, for example, the genes *KY21.Chr1.1992* and *KY21.Chr8.1350* have a gradient of expression (**Supplementary Fig. 3-12a,b**), and HCR FISH showed high expression in large granules hemocytes (LGH) and low expression in MCs (**Fig. 2b** bottom right, **Fig. 2c** “16 (high)” vs. “16 (low)”, **Supplementary Fig. 3-12c–f**). In another case, cluster 22 maps to both the small round morphology and SRCs in a range of sizes. This morphological heterogeneity is consistent with small round cells giving rise to small SRCs, which then grow to large SRCs, as was suggested some seventy years ago^36^. In total, the map of cell states to morphotypes shows that morphological definitions fail to reflect the full diversity of *C. robusta* immune cell states, and that some morphologically distinct cells are transcriptionally similar.

These morphological annotations come with caveats. First, live morphologies can depend on sample preparation; we found, for instance, that HA-3 morphologies differ across buffer conditions (**Supplementary Fig. 2f**). Additionally, some studies have used histological staining to distinguish morphotypes^45^, but our annotations lack this information since these stains are not yet compatible with HCR FISH. Finally, our annotation is not comprehensive: HCR FISH of markers for a few transcriptional clusters failed to label any observed cells (markers of 17, 23, 30, and 14.2; see **Supplementary Fig. 2c**); one of these clusters (23) was only detected in 5/9 SNP profiles (**Fig. 1f**), and might represent an environmentally induced state. We also observed that orange cells’ pigmentation blocked some fluorescence wavelengths (**Supplementary Fig. 2g**), which may have prevented their annotation with HCR FISH. Nevertheless, our results offer a morphotype annotation of most cell clusters.

### Observing *in vivo* cell behavior

To observe cell morphologies and behaviors in live animals, and to enable future *in vivo* analysis of cell states, we designed fluorescent reporters for six transcriptional cell states: GA, HA-1, HA-2, HA-3, URG-1, and cLRP-1. We isolated 0.5–3kb segments of 5’ intergenic DNA located upstream of transcriptional start sites of genes specifically enriched in each cell state (**Supplementary Fig. 5a**). We cloned these DNA elements upstream of green fluorescent protein (GFP), introduced them into *C. robusta* embryos by electroporation, then grew the animals until the juvenile stage, at which point the circulatory system has developed^52^ (**Fig. 3a**).

**Figure 3:**
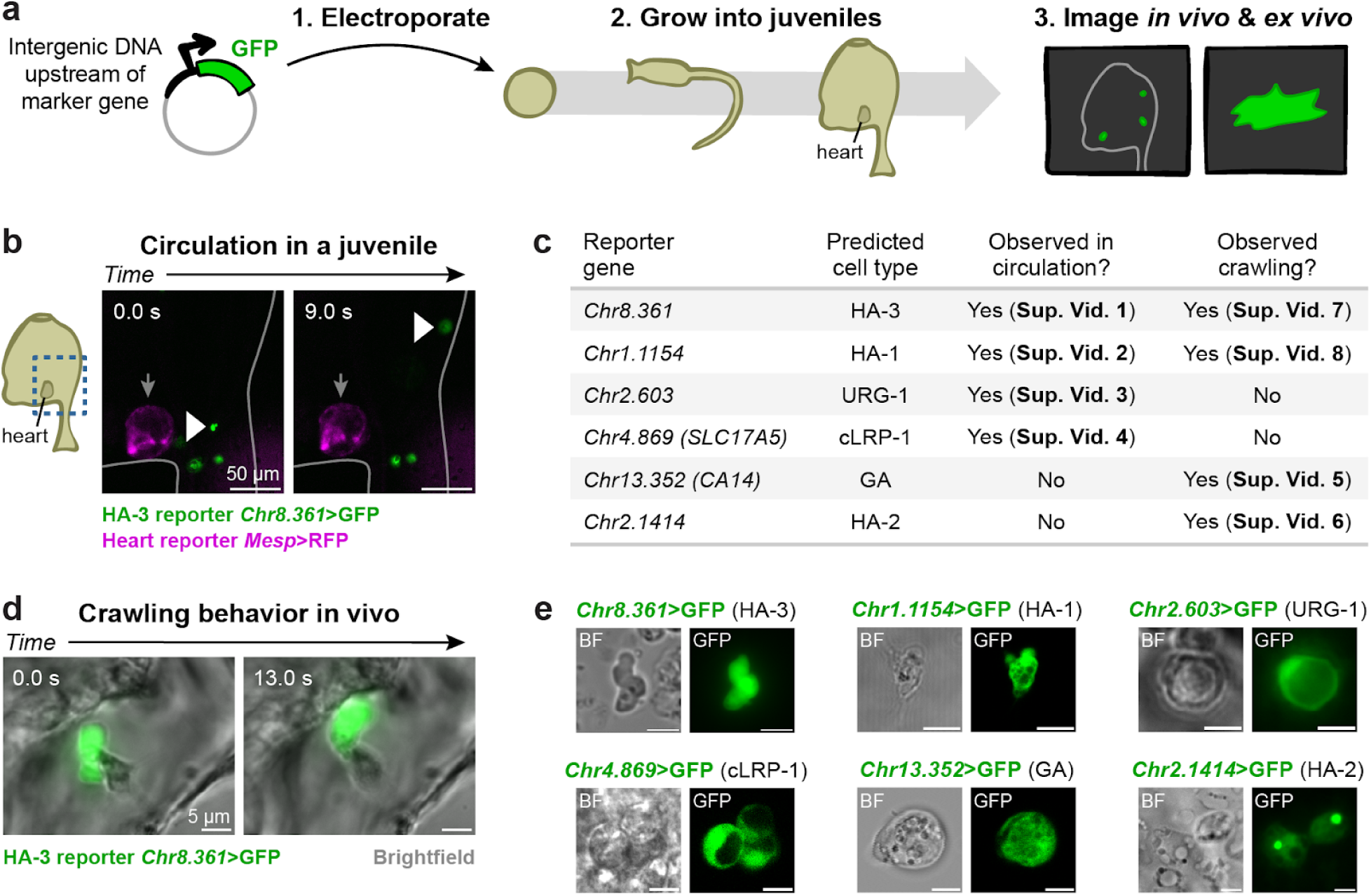
Live reporters label transcriptional clusters for *in vivo* observation **(a)** Approach for labeling live blood cells with fluorescent reporters. **(b)** Frames from **Supplementary Video 1** showing an example of a GFP+ cell (presumed to be HA-3) circulating in a juvenile. White arrowheads point to a circulating GFP+ cell. Grey arrows indicate the heart. Scale bars = 50 µm. **(c)** Table listing reporters, showing the cell state labeled, marker gene used for reporter design, and whether circulation and/or crawling behavior was observed *in vivo*. Supplementary Videos showing circulating/crawling behavior are indicated for each reporter. **(d)** Frames from **Supplementary Video 6** showing an example of a GFP+ cell (presumed to be HA-3) crawling in a juvenile. Scale bars = 5 µm. **(e)** Representative images of single-cell morphologies observed. HA-3, GA, and HA-1 morphologies were observed *in vitro*. URG-1, cLRP-1, and HA-2 morphologies were observed *in vivo*. See Supplementary Fig. 5b–g for more example images and separated channels. Scale bars = 5 µm.

For each reporter, we first checked whether GFP+ cells in juvenile animals included cells in circulation. Four of the transgenes labeled cells in circulation, predicted to be HA-3, HA-1, URG-1, and cLRP-1 (**Fig. 3b,c**, **Supplementary Videos 1–4**). Cells labeled by the remaining two transgenes, predicted to be amoebocytes GA and HA-2, were observed crawling within the juvenile body (**Fig. 3c**, **Supplementary Videos 5,6**). Cells expressing HA-3 and HA-1 reporters also exhibited this motility and appeared to alternate between circulation and crawling behavior (**Fig. 3c,d**, **Supplementary Video 7,8**).

We then confirmed the cells’ morphologies *in vivo* in the juveniles and *ex vivo* after cell isolation. GFP+ cells for all six of the labeled states included the expected morphologies (**Fig. 3e**, **Supplementary Fig. 5b–g**). Together, these results support the classification of observed cell states, demonstrate *in vivo* circulation or ameboid-like motility of some cells, and identify reporter genes enabling live analysis of several blood cell states in tunicates.

### Characterizing identity and hierarchy in *C. robusta* blood

With access to the transcriptional and morphological atlas of blood cells, several questions can be explored relating to these cells’ functions. We provide evidence that the observed cell states include a hematopoietic hierarchy. We also identify phagocytes, as well as cells expressing vanadium-binding proteins, enzymes associated with gas transport, and genes related to immune function.

## 1. A hematopoietic hierarchy in circulation

Unlike vertebrates, *C. robusta* and other tunicates are thought to have an abundant population of circulating hematopoietic progenitor cells, which are proliferative and have a hemoblast-like morphology^36,39,45^. We indeed find evidence of a hematopoietic hierarchy in circulation, with hemoblast-like and small round cells being progenitors. Hemoblast-like and small round cells (clusters 0/14.1, 1/7, and 21; see **Fig. 2b**) are part of a continuum of states expressing multiple markers of proliferation, including orthologs of *MKI67,* cyclin B1 (*CCNB1*), and topoisomerase 2a (*TOP2A*) (**Fig. 4a**, **Supplementary Fig. 6a,b**). Using RNA velocity analysis of spliced vs. unspliced transcripts^53^, we observed that nearly all cells appear to differentiate away from the cluster 0/14.1 hemoblast-like cells (**Fig. 4b**). We therefore labeled these cells as candidate multi-potent progenitors (cMPP) and labeled other proliferative clusters as candidate lineage-restricted progenitors (cLRP) (**Table 1**, **Supplementary Table 2**). cMPPs could be self-renewing or could themselves be replenished from a hematopoietic niche not sampled in this study.

**Figure 4:**
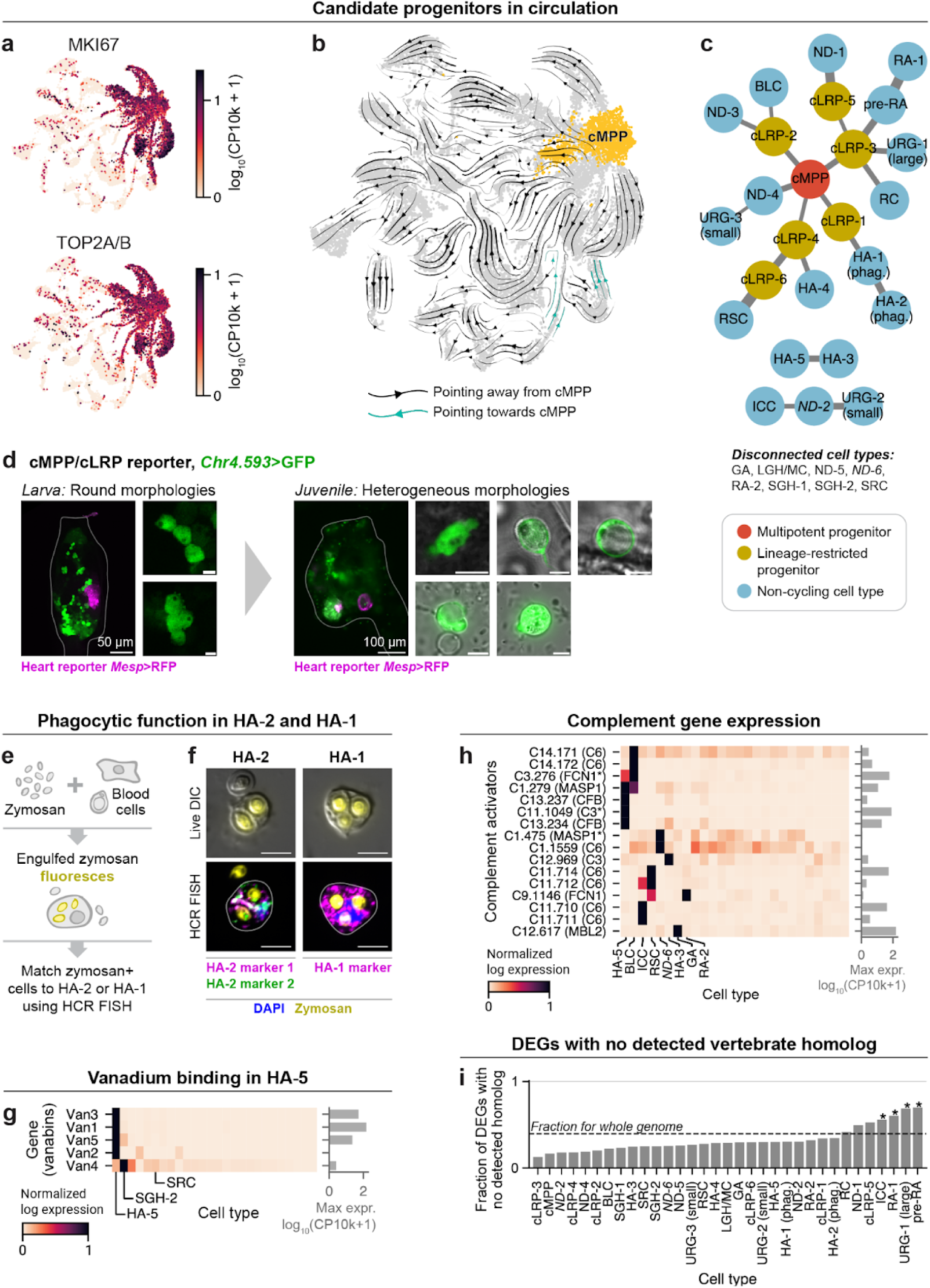
*C. robusta* blood contains a likely hematopoietic hierarchy and mature immune cell states **(a)** Expression of KY21.Chr2.1280, orthologous to *MKI67*, a marker of proliferation. **(b)** RNA Velocity of the *C. robusta* blood dataset. Arrows pointing away from the cMPP cluster are black; arrows pointing towards cMPP are in light blue. **(c)** Coarse grain tree of cell states showing the inferred hematopoietic hierarchy. **(d)** Images showing that GFP expressed under a marker of cMPP and cLRP clusters labels round cells in swimming larvae (left) and a morphologically diverse population after metamorphosis (right). See Supplementary Fig. 5i for separated channels of the juvenile single-cell images. **(e)** Approach for testing whether HA-2 and HA-1 are capable of phagocytosis. **(f)** Images showing that cells positive for markers of HA-2 and HA-1 can phagocytose zymosan. **(g)** Vanadium-binding proteins are expressed highly in HA-5 and at lower levels in SGH-2. **(h)** *C. robusta* homologs of several complement activator genes are expressed across several blood cell states. **(i)** The fraction of differentially expressed genes (DEGs) for each cluster which have no detected vertebrate homolog. DEGs are determined via the Wilcoxon rank-sum test (FDR<0.05, log2 fold change > 2). Genes are considered to have no detectable vertebrate homolog if a gene has both (1) no vertebrate homolog detected by OrthoFinder, and (2) no detected BLAST similarity to any human gene (with an e-value<1e-6). Scale bars are 5 µm unless otherwise noted. In this and all subsequent figures, cell states which both are present in fewer than 7/9 SNP profiles (see Supplementary Fig. 4b) and have not been verified by HCR FISH are labeled with italic text.

To infer *C. robusta*’s hematopoietic hierarchy, we used a k-nearest neighbor graph of single cell transcriptomes to construct a coarse-grained tree connecting cMPPs to differentiated cell states^54^. The resulting tree (**Fig. 4c**) connects 20 of the 33 annotated clusters in five major lineages, with seven connected clusters representing cycling cMPP and cLRP cell states. Five clusters not linked to the tree form two separate continua: HA-3 and HA-5; and separately irregular compartment cells (ICC), URG-2, and a cluster whose morphology was not determined (ND-2) (**Fig. 4c**, bottom). The remaining nine clusters—including SRCs, GAs, and LGH/MCs—are disconnected from the rest. These cells may represent long-lived cell states that lack constitutive precursors, or they might be generated by distinct lineage-restricted progenitors in a niche.

To collect further evidence that the cMPP/cLRP cells are progenitors, we generated a live-cell reporter, *Chr4.593>GFP*, from a gene broadly expressed across these clusters (*KY21.Chr4.593*, homolog to human fibrillarin genes *FBL* and *FBLL1*; **Supplementary Fig. 5a**). We used the reporter to track the earliest appearance of these cells during animal development, as well as their capacity to give rise to differentiated cells. In several tunicate species, blood cells arise from undifferentiated mesodermal progenitors that form pouch-like clusters in tailbud-stage embryos^55^, and they are thought to do the same in *C. robusta*^52,56^ We indeed observed *Chr4.593>GFP*+ cells in these mesodermal pouches at the tailbud stage, while reporters for differentiated cells HA-3 and GA labeled no cells at this stage (**Supplementary Fig. 5h**). From this point onwards, *Chr4.593>GFP+* cells were continuously present: in swimming larva, the cells dispersed, with a subset appearing to migrate towards the anterior of the trunk (**Supplementary Fig. 5h**); one day after the start of metamorphosis, some labeled cells were concentrated at the base of the trunk, and others in an anterior organ called the stolon; a few hours later, some *Chr4.593>GFP+* cells were highly motile and appeared to extravasate through the epidermis into the tunic, as has been previously described^57^ (**Supplementary Video 9**); and after metamorphosis, at the early juvenile stage, some *Chr4.593>GFP+* cells circulated in the bloodstream, while others remained static in the tunic, between the ciliary gill slits, and in the dorsal region of the juvenile body (**Supplementary Video 10**). At this stage, in juveniles, *Chr4.593>GFP+* cells showed a range of morphologies, including round cells, vacuolated cells, and crawling amoebocytes (**Fig. 4d**, **Supplementary Fig. 5i**). We do not currently know whether all *Chr4.593>GFP+* cells are transcriptionally similar to adult cMPP/cLRP cells.

Nevertheless, these results are consistent with cMPP/cLRP-like cells developing from embryonic mesodermal progenitors and being capable of giving rise to a diverse population of cells.

## 2. Hyaline amoebocytes HA-2 and HA-1 are phagocytes

There is broad conservation of metazoan phagocytosis and associated transcriptional regulators^26^, and past work identified hyaline amoebocytes and granulated cells as phagocytes in *C. robusta*^34,44^. We show here that HA-2 and its hypothesized precursor state, HA-1 (see **Fig. 4c**) are phagocytes. The HA-2 cell cluster shows evidence of being phagocytic based on (i) its expression of several markers of phagocytosis, including the widely conserved phagocyte-associated transcription factor *Cebpa*^26^ (**Supplementary Fig. 7a,b**); and (ii) images from HCR FISH experiments of what appears to be engulfed cells (**Supplementary Fig. 3-6c,d**). To test whether HA-1 and HA-2 cells are capable of phagocytosis, we incubated blood cells with fluorescent zymosan, then labeled marker genes of the states using HCR FISH (**Fig. 4e**). Phagocytic cells indeed expressed HA-1 and HA-2 marker genes, linking these transcriptional states to this function (**Fig. 4f**, **Supplementary Fig. 7c–i**). These clusters accounted for 85% of observed zymosan-positive cells (240/282 cells), so HA-1 and HA-2 account for most but not all phagocytes in circulation (**Supplementary Fig. 7j**).

## 3. Hyaline amoebocytes HA-5 are likely vanadocytes

Some species of tunicates contain vanadocytes, blood cells which accumulate high levels of vanadium, though these cells’ function is unclear^58^. Vanadocytes have not been documented in *C. robusta* blood^58^, and have been proposed to have an SRC morphology in other species^59^. However, in the scRNA-seq data, HA-5 cells express high levels of genes encoding vanadium binding proteins^60^ (150 counts per 10,000 [CP10k] of *Van1*, 51 CP10k of *Van3*, and 21 CP10k of *Van5*), while SRC did not express these genes (**Fig. 4g**). This expression pattern suggests that HA-5 cells may be vanadocytes. SGH-2 also expresses vanadium-binding proteins, but at much lower levels (1.5 CP10k of *Van4*). HA-5 cells show weak yellow pigmentation (**Fig. 2b**, **Supplementary Fig. 3-21c**), matching the +5 oxidation state of vanadium, while the +3 and +4 states (blue and green) have been identified in vanadocytes of other tunicate species^59^. HA-5 may thus represent a distinct vanadocyte population, but we still know little about the identity and function of these cells across tunicates^59^.

## 4. GA, HA-3, and other cells express enzymes associated with gas exchange

A common function of blood is to facilitate gas transport. *C. robusta* have no red blood cells, and although globin genes have been identified^61^, their blood does not store or accumulate oxygen^58^. Consistent with this, we find that globin gene expression is extremely low in the blood (**Supplementary Fig. 6c**). The highest expression of any globin is *Hb2* (0.09 CP10k in RA-2 and 0.07 CP10k in SRC), expressed at a level several orders of magnitude lower than that found in vertebrate red blood cells^50^. However, red blood cells also play a role in carbon dioxide transport, expressing carbonic anhydrases (CAs) which catalyze conversion of carbon dioxide into soluble carbonic acid^62^. We observed that GA, HA-3, URG-2, blebbing-like cells (BLC), and other *C. robusta* blood cells express high levels of homologs to CA genes (e.g. 42 CP10k of CA *KY21.Chr13.352* in GA cells) (**Supplementary Fig. 6d**). One or more of these cells might play a role in gas transport and respiration.

## 5. Expression of genes with homology to vertebrate immune factors

We found complex expression patterns in *C. robusta* blood of genes with predicted homology to vertebrate immune proteins, including complement factors, cytokines, cytokine receptors, and PRRs (**Supplementary Fig. 6e–g**). The specific expression of these genes in different blood cell subsets indicates they are indeed likely to be immune proteins. Several of these immune gene families appear to have expanded in *C. robusta.* The complement factor *C6*, a single gene in vertebrates, is represented in *C. robusta* by 13 copies across 5 chromosomal clusters (**Supplementary Table 3**). Of these copies, seven are expressed in the blood, specifically in BLCs, ICCs, round spreading cells (RSCs), and ND-5 (**Fig. 4h**). This dramatic expansion might be associated with the acquisition of novel protein functions.

In the case of *C6* and several other complement factors, expression in *C. robusta* blood cells does not align with characterized expression patterns in vertebrates. In humans, complement factor *C6*, complement factor B (*CFB*), and mannose-binding lectin (*MBL2*) are expressed and secreted by hepatocytes in the liver, rather than by blood cells^63,64^. In *C. robusta*, in addition to *C6* expression in blood cells, *CFB* is expressed in HA-5, the putative vanadocyte cluster, while *MBL2* is expressed in GA (**Fig. 4h**). The expression of these factors in tunicate mesoderm (blood) vs. vertebrate endoderm (liver) suggests considerable plasticity in the cellular deployment of these genes.

## 6. Expression of genes with no detected vertebrate homologs

So far, we have hypothesized cell function based on expression of genes with predicted homology to known vertebrate genes. However, a large fraction of *C. robusta* genes have no known homologs in vertebrates. In the *C. robusta* genome, 40% of genes have no vertebrate homolog detected by either orthology inference (OrthoFinder^65^) or BLAST (**Supplementary Table 4**). Notably, in three *C. robusta* blood cell states, the fraction of enriched differentially expressed genes (DEGs) that lack detectable similarity to vertebrate genes is considerably higher (FDR<0.01, Fisher’s Exact Test) (**Fig. 4i**, genes listed in **Supplementary Table 5**). These cells are ICCs (56% non-vertebrate genes), URG-1s (69%), and refractile amoebocytes (70% in pre-RA, 60% in RA-1). For these cells, and possibly for others, knowledge from vertebrate immunity may have limited utility in understanding their function.

### Evidence of major divergence between protochordate and chordate blood cells

Genomic and functional analyses in tunicates and other invertebrates reveal many immune-specific genes and functions shared with vertebrates^15,17–27^. However, the degree to which expression of these shared genes is distributed similarly across conserved immune cell types remains unclear. We set out to systematically evaluate the degree of conservation or divergence in immune cell states between *C. robusta* and vertebrates as seen through whole transcriptome analysis.

We chose to focus on humans as the vertebrate species for comparison^64^ and evaluated cell state homology using the Self-Assembling Manifold mapping (SAMap) algorithm^66^. SAMap uses a many-to-many gene homology mapping to score similarity in transcriptional states in a manner robust to vertebrate- and tunicate-specific gene duplications^67^ (**Fig. 5a**). As a benchmark, we separately compared human blood and bone marrow to differentiating blood cells in zebrafish^68^ (diverged from mammals 420 million years ago^35^). This benchmark established a SAMap similarity score that distinguishes one-to-one orthologous states (scores of 0.3 or above, **Supplementary Fig. 8a**).

**Figure 5:**
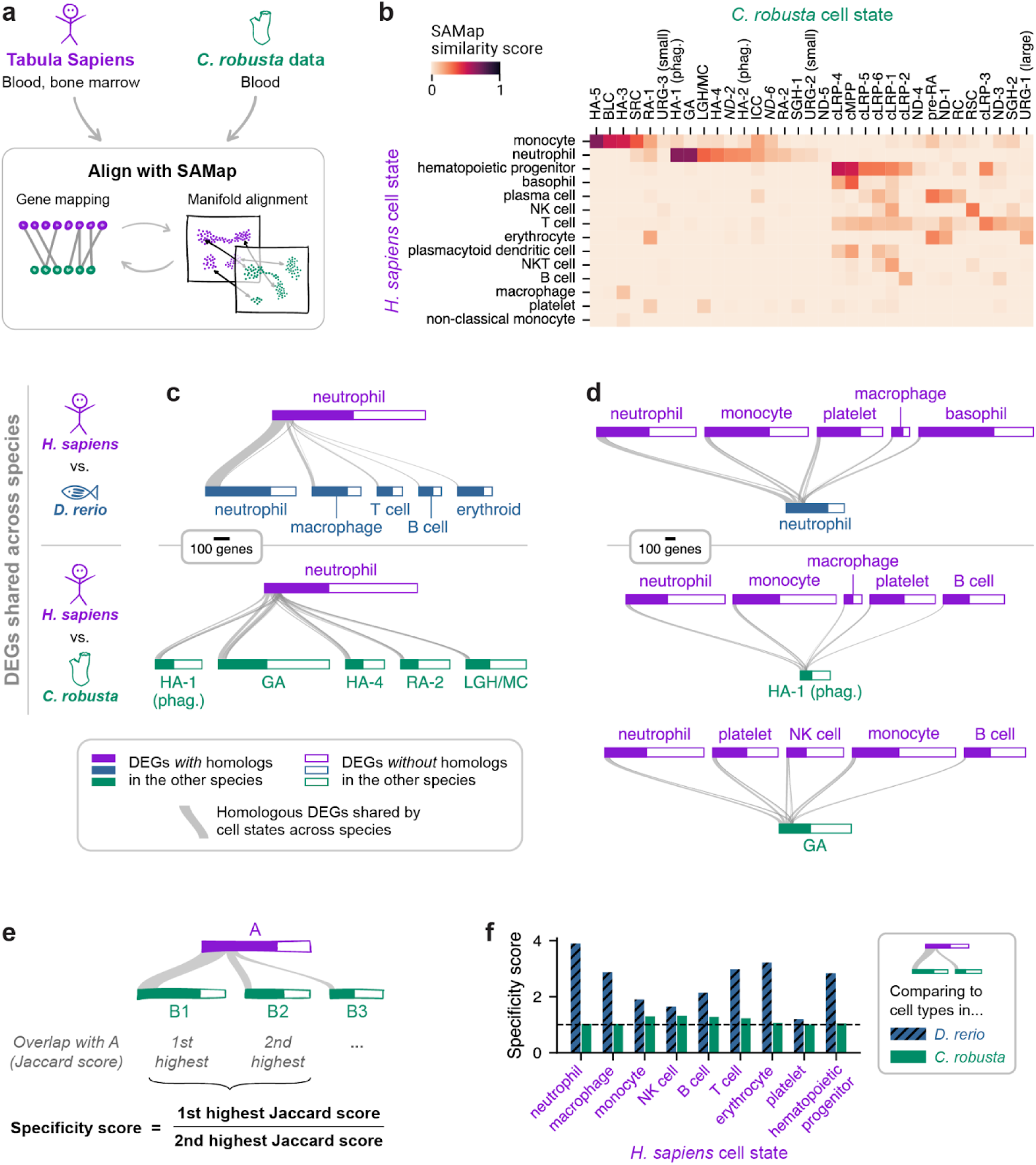
Comparison to human blood shows dramatic divergence of immune cell states between tunicates and vertebrates **(a)** *C. robusta* blood was compared to human blood and bone marrow data from Tabula Sapiens using SAMap. **(b)** Similarity matrix generated by SAMap, showing the similarity score between each human and *C. robusta* cell state. **(c, d)** Generalized Sankey (GS) plots illustrating the number of enriched DEGs shared between cell states across species. Horizontal bar widths represent the number of enriched DEGs in each cell state; solid-filled areas within these bars show the number of those DEGs with a homolog in the other species, as determined by OrthoFinder^65^; and the widths of connections indicate how many DEGs in two connected cell states are homologous. Panel **(c)** shows human neutrophils vs. the most similar 5 zebrafish (top) or *C. robusta* (bottom) cell states. Panel **(d)** shows the most similar cell states in zebrafish (top) or *C. robusta* (bottom) to human neutrophils vs. the most similar 5 human cell states. **(e)** Calculation of specificity score, defined for a cell state in species 1 as the ratio between the highest and second highest Jaccard similarity scores with cell states in species 2. The specificity score for cell state A in species 1 is high (>1) if A matches closely to a single cell state in species 2, and low (∼1) if A matches closely to more than one cell state in species 2. **(f)** Specificity scores for human cell states when compared against *C. robusta* or zebrafish. Scores near 1 for the comparison to *C. robusta* show that human cell states match closely to multiple *C. robusta* cell states.

Using this benchmark, the SAMap alignment of human and *C. robusta* blood maps human hematopoietic progenitors and basophils to *C. robusta* cMPP and cLRP clusters; monocytes to HA-5, BLC, HA-3 and SRC; and neutrophils to HA-1 (phagocytes), GA, and LGH/MC (**Fig. 5b**). The remaining 10 human and 19 *C. robusta* cell states do not map to any cell state in the other species with similarity >0.3 (**Fig. 5b**).

SAMap similarity scores are suggestive of cell state homology, but we worried that the algorithm might over-emphasize weak similarities driven by a small number of genes when comparing cells across large evolutionary distances. To critically examine cases of possible cell state homology, we developed a gestalt approach using generalized Sankey (GS) plots to visualize shared enriched genes between species’ cell types (**Fig. 5c,d**, **Supplementary Fig. 8b**, see **Fig. 5** legend for GS plot description). In addition, the proportion of shared genes between human cell types and *C. robusta* cell states was quantified using a Jaccard index (**Supplementary Fig. 8c–e**).

We focus on the relationship between human neutrophils and *C. robusta* cell states, as these cells were among the most similar based on SAMap. As a control, we first confirmed that zebrafish neutrophils are the most similar to human neutrophils by a large margin (**Fig. 5c** top), with a Jaccard similarity score 3.9 times higher than the next most similar zebrafish cell state; we call this ratio a specificity score (**Fig. 5e**). By contrast, human neutrophils mapped evenly to multiple *C. robusta* cell states (**Fig. 5c** bottom), with a specificity score of 1.03 (**Fig. 5f**). A converse analysis likewise shows that HA-1 (phagocyte) and GA cells, which were identified as most similar to human neutrophils, are not significantly more similar to neutrophils than to other human cell types (**Fig. 5d**), with specificity scores of 1.5 and 1.3, respectively. These results suggest there is no clear homologous cell state in *C. robusta* to human neutrophils. Instead, the genes defining neutrophil identity in humans appear mixed across *C. robusta* cell states, and *C. robusta* cell state identities appear mixed across human cells. We see similar results for comparing human monocytes to *C. robusta* blood cells (**Fig. 5f**, **Supplementary Fig. 8f,g**). Overall, there appears to be a lack of clear blood cell homology to vertebrates.

### Evidence of divergence in hematopoietic regulation

The divergence we observe between vertebrate and tunicate blood cell states also extends to the transcription factors that specify cell types. In vertebrate hematopoiesis, fate specification is mediated by the combinatorial expression of several transcription factors (TFs)^69,70^ listed in **Fig. 6a**. Many of these TFs have orthologs in *C. robusta* that are variably expressed across immune cells (**Fig. 6b**). However, the patterns of TF expression are divergent between human and *C. robusta*, as supported by two pieces of evidence. First, we analyzed differentially expressed TFs at putative branch points in the hypothesized *C. robusta* hierarchy (**Fig. 6c**). Although these TFs include homologs of known fate regulators in mammalian hematopoiesis (*Spi1/B/C*, *Nfkb*, *Cebpz*, *Egr*, and *Stat.b*), they also include TFs not expressed in mammalian hematopoietic cells^50,64^—such as *Sox14/15/21* (upregulated in HA-4) and *Hey* (upregulated in ND-4). Second, TFs homologous to those involved in vertebrate fate specification show entirely distinct patterns of co-expression in *C. robusta* (**Fig. 6b**). In mammals, *SPI1* is coexpressed with *IRF8* in monocytes^70^; in *C. robusta*, co-expression with *IRF8* is absent. Similarly, three TFs typically coexpressed in neutrophils—*CEBPA*, *GFI1*, and *LEF1*—are expressed in three separate cell states in *C. robusta*. Examining all co-expression pairings of these canonical hematopoietic TFs (listed in **Fig. 6a**) across cell states leads to a similar conclusion that TF co-expression patterns have diverged between tunicates and vertebrates (**Fig. 6d** left). As a control, the same analysis carried out between humans and zebrafish shows better conservation of co-expression (**Fig. 6d** right).

**Figure 6:**
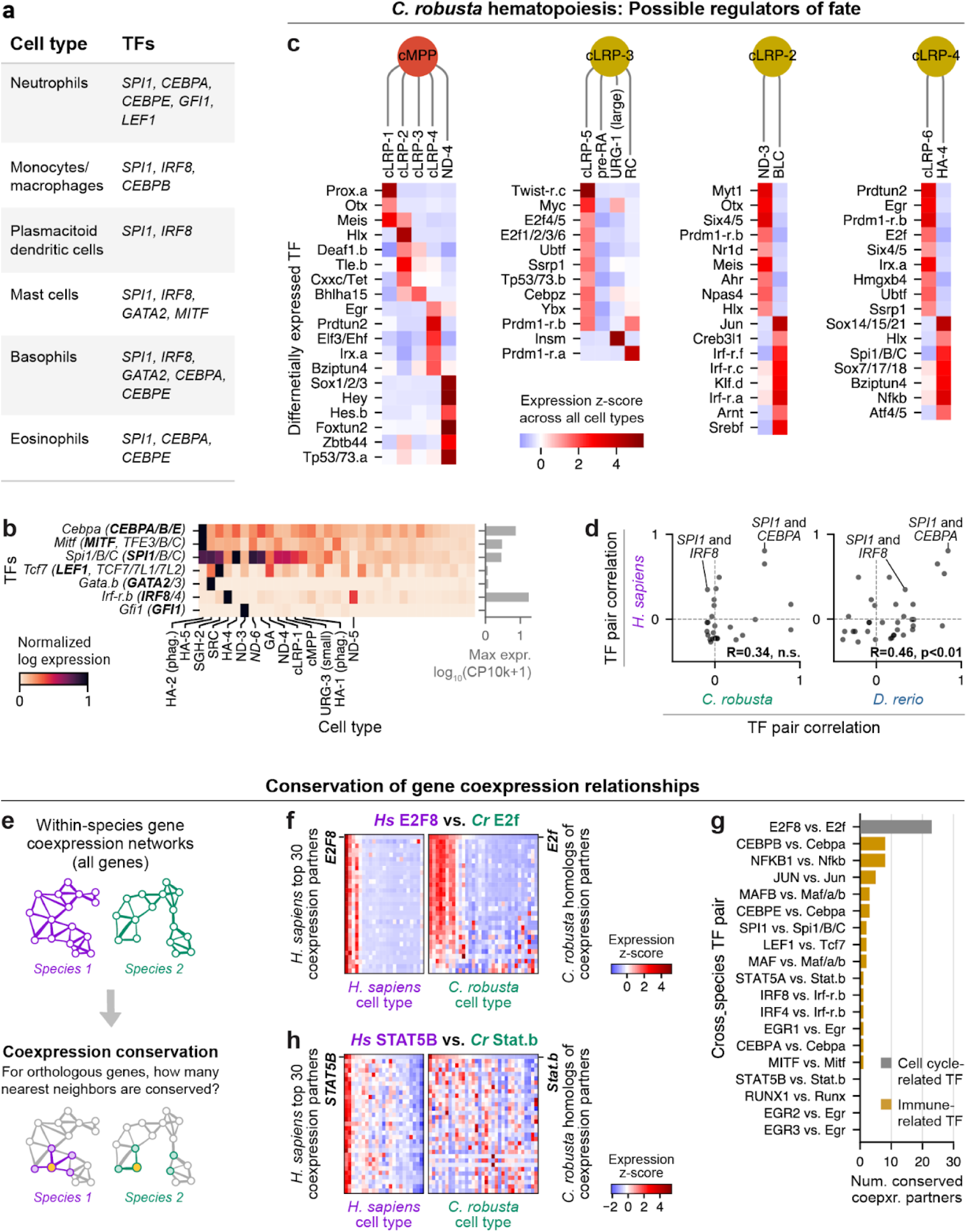
Transcription factor expression has diverged between *C. robusta* and vertebrates **(a)** TF combinations important for fate specification in mammalian hematopoiesis^70^. **(b)** Heatmap showing expression in *C. robusta* cell states of TFs homologous to those that specify fate in mammalian hematopoiesis. Human homologs of each TF are listed in parentheses for each gene, with TFs listed in panel **(a)** bolded. **(c)** Heatmaps showing differentially expressed TFs at each branch point in the *C. robusta* hypothesized hematopoietic hierarchy in Fig. 4c. Gene KY21 IDs are listed in **Supplementary Table 6**. **(d)** Scatter plot showing the expression correlation of TF pairs from panel **(a)** for human vs. (left) *C. robusta* or (right) zebrafish. Correlation of each TF pair within each species is calculated across mean log-transformed expression in all cell states. In cases of one-to-many TF homology, multiple points are plotted; e.g. there are three points for *C. robusta Cebpa* vs. each of human *CEBPA*, *CEBPB*, and *CEBPE*. The overall correlation, R, for each scatter plot is listed, along with the statistical significance. **(e)** Cartoon illustrating the approach to assessing conservation of gene co-expression across species, based on methods presented by Crow et al.^71^ **(f, h)** Heatmaps showing examples of a TF with high co-expression correlation, **(f)** human *E2F8* and *C. robusta E2f* and **(h)** human *STAT5B* and *C. robusta Stat.b*. Left: top 30 human co-expression partners for *E2F8*. Right: homologs of those co-expression partners in *C. robusta*. Human *E2F8* and *C. robusta E2f* expression is the first row of each heatmap; co-expression partners are then ordered by decreasing correlation between the *C. robusta* homologs and *E2f*. **(g)** Number of co-expression partners shared across species for *E2F8/E2f* (grey) and several immune-related TFs (gold).

We next expanded this analysis to systematically probe the conservation of regulatory relationships between TFs and their possible targets using the proxy of gene co-expression. Adapting a method established by Crow et al.^71^, we quantified the degree to which each TF’s co-expression relationships to other genes are maintained between species (**Fig. 6e**). The TFs with the most highly conserved co-expression relationships include those which regulate the cell cycle, such as human *E2F8* and *C. robusta* ortholog *E2f*. Out of the top 30 genes coexpressed with *E2F8* in the human dataset, 24 have *C. robusta* homologs which remain correlated with *E2f* (**Fig. 6f**). By contrast, TFs associated with mammalian hematopoiesis have much lower levels of co-expression conservation (**Fig. 6g**). In a particularly divergent case, *C. robusta Stat.b* appears to be anti-correlated with several homologs of human *STAT5B* co-expression partners (**Fig. 6h**). Repeating this analysis between humans and zebrafish shows significantly better conservation of co-expression (**Extended Data. Fig. 9a**). These results hold when conservation is quantified with a second statistical metric introduced by Crow et al.^71^ (**Supplementary Fig. 9b,c**).

Altogether, these results suggest that, in addition to divergence of mature cell states, the regulatory relationships governing hematopoiesis have diverged significantly between vertebrates and tunicates.

## Discussion

In this study, we establish a joint transcriptional and morphological atlas of *C. robusta* blood cells. We introduce live reporters that allow observing several cell states *in vivo*, and we present evidence that some differentiated cells arise from a hematopoietic hierarchy in circulation. Our atlas demonstrates that *C. robusta* blood cells occupy a set of transcriptional states that are far more diverse than suggested by previously identified morphological archetypes. We then show that blood cells have diverged significantly since the last common ancestor of vertebrates and tunicates. This divergence is reflected in poor homology to human cell states, distinct transcription factor co-expression, and cell state–specific expression of genes that have no detected vertebrate homologs.

From what we know in vertebrates, it was to be expected that the number of transcriptional states would exceed the number of morphotypes. In mammals, the discovery of B, T, and dendritic cells lagged decades after initial morphological immune cell classifications, and it took decades more work to identify subsets of T cells, dendritic cells, monocytes, and neutrophils^72,73^. Cell state diversity reflects the existence of multiple developmental lineages, as well as differences in cell age, responses to inflammatory signaling, and infection.

Nonetheless, the diversity of cell states documented here extends well beyond that in vertebrates. This diversity suggests that *C. robusta* may support qualitatively different strategies for sensing, responding to, and adapting to pathogens. Such strategies are not likely to be discovered by focusing only on features common to vertebrates. We speculate that similarly complex, still unobserved immune cell repertoires are common across invertebrates. Defining the roles of these cells may be fruitful: historically, major breakthroughs in immunity—including the discoveries of phagocytosis, antimicrobial peptides, and toll-like receptors—have emerged from studies of invertebrates^25,74^.

Still, our study has at least two limitations in scope. First, we do little to clarify the function of identified cell states, or the role of most genes specifically expressed in them. Several functional assays exist for tunicate blood cells^24,38,41,45^, and these can now be revisited while tracking multiple transgenically labeled cell subsets in live animals. Sessile tunicates, including the juveniles of *C. robusta*, are stationary and transparent, making them practical models for live immune assays. The second limitation lies in the difficulty of determining gene and cell state homologies. State-of-the-art methods for identifying gene homology are imperfect^71^, in our case failing to identify known homologs of toll-like receptors in *C. robusta*^75^. Methods such as SAMap seek to identify homologous cell states with flexible gene homology, but with distant species the results are hard to interpret, as seen from the GS plots introduced here. Cell state homology might be better determined by clustering cell states from many species. Doing so would require blood cells sampled from multiple phyla. Such data can now readily be collected and can be used to test whether certain blood cell states in vertebrates and tunicates derive from shared ancestral cell types.

Finally, we reflect on what can be learnt about immune cell evolution. New cell types are thought to arise by a ‘sister cell’ mechanism: duplication of a developmental lineage, followed by divergence in the set of genes turned on in each lineage^1–3^. Here we find that the divergence between *C. robusta* and humans is too large to identify sister cell relationships, if they exist: genes specific to human immune cell states appear split across multiple *C. robusta* cell states, and vice versa *C. robusta* cell state identity is split across human cells. We also fail to find examples of conserved co-expression between transcription factors implicated in lineage determination and their putative targets. It is still possible that vertebrate and tunicate blood cells descended from common ancestral cell types, or, alternatively, these cells might have evolved from distinct ancestral populations. In either case, the divergence between vertebrate and tunicate immune cells suggests a pliability to which genes and functions can be expressed in the same cell. Altogether, this atlas of *C. robusta* blood provides a step towards documenting the diversity of immune systems and towards understanding how that diversity arises during evolution.

## Declarations

### Data Availability

The datasets generated and analyzed in the current study are available in the Gene Expression Omnibus (GEO) repository: the full processed dataset and May 2023 libraries are available from Series <u>GSE296253</u>, and the April 2022 library is available from Sample GSM8869531 of Series <u>GSE292926</u>. Imaging data is available upon request. Code for data analysis is available on Github at https://github.com/AllonKleinLab/paper-data/tree/master/Scully_Ciona_blood_2025.

An interactive version of the scRNA-seq dataset is available at https://kleintools.hms.harvard.edu/paper_websites/scully_ciona_robusta_blood/.

## Acknowledgments

We thank Lillian Horin and Tim Mitchison for providing zymosan for the phagocytosis assay. We are grateful to Laura Bagamery for processing and annotating the zebrafish hematopoiesis dataset. We thank Andrew Murphy for his expertise in building aquariums and supporting animals, and John Bishop for kindly sharing an image of adult *C. robusta* animals shown in Fig. 1a. We acknowledge the Single Cell Core at Harvard Medical School for performing the scRNA-seq sample preparation, the Core for Imaging Technology & Education at Harvard Medical School for help with light microscopy, and the The Bauer Core Facility at Harvard University for sequencing services. We thank all core facility staff for their technical assistance. The work was supported in part by an Edward Mallinckrodt Jr Scholar Award and NIH grant R01GM153805 to AMK, and NSF grant 2052517 to BD.

## Authors’ contributions

TDS and AMK conceptualized the work. AMK and BD supervised the work. CJP carried out live reporter design and experiments, and NGF carried out RNA Velocity analysis. TDS carried out all other experiments and data analysis including scRNA-Seq, RNA FISH, phagocytosis assays, cross-species comparison, gene orthology analysis, development of GS Plots. TDS, BD, and AMK wrote the manuscript. All authors read and approved the final manuscript.

## METHODS

Blood Collection from *C. robusta* Adults

### Animal Husbandry

*Ciona robusta* adults were collected from the wild in Carlsbad, California, USA by M-REP. They arrived in the lab <24 hours after collection, where they were kept at 17–18°C in 35–40 ppt artificial seawater (Instant Ocean SS15-10) and fed 6–7 days a week with 15 mL of Phyto-Feast (Reef Nutrition). Blood was collected for experiments 1–14 days after arrival in the lab, during which time animals remained healthy by visual inspection. Animals for scRNA-seq experiments were used 1 day (April 2022 collection) and 7 days (May 2023 collection) after arrival. Prior to blood collection, animal size was recorded, see **Supplementary Table 7** for animal size measurements.

### Blood Cell Collection

To reduce movement during dissection, animals were relaxed by incubating 3–5 adults for 3–4 hours at 4 °C in 400 mL artificial seawater from the aquarium system with 5 g of 99% L-menthol crystals (Thermo Scientific A10474). Volume and weight measurements were approximate. To collect blood, animals were dissected to expose a large blood vessel adjacent to the heart and running along the endostyle (“ventral vessel” in Millar 1953^36^). Blood was drawn from the vessel with a zero dead space tuberculin syringe with a 25G needle (Exelint 26046), then transferred to a microcentrifuge tube on ice. For scRNA-seq experiments, this tube was first coated with bovine serum albumin (BSA) by washing with 0.5 mL of 10% (v./v.) BSA (Sigma 9048-46-8) in Dulbecco’s phosphate-buffered saline (DPBS, Gibco 14200075).

## Profiling Blood Cells with scRNA-seq

### Single-Cell Barcoding, Library Preparation, and Sequencing

Methods for scRNA-seq collection for animals A-E (**Supplementary Table 7**), harvested in April 2022, are described in a separate technical paper^47^ (the “PBS-M” sample). For the May 2023 scRNA-seq collection, samples were processed according to the same approach with just one change noted below: first, blood was harvested from the 7 animals F-L (**Supplementary Table 7**). Equal blood volumes were then pooled for animals F,G,H,I, and then again for animals F,J,K,L, such animal F contributed blood to both pools. Blood was then filtered through a 40 µm strainer (pluriSelect 43-10040-60) and transferred to a microcentrifuge tube, which had been coated with bovine serum albumin (BSA) by washing with 0.5 mL of 10% (v./v.) BSA (Sigma 9048-46-8) in Dulbecco’s phosphate-buffered saline (DPBS, Gibco 14200075). Blood cells were pelleted by spinning at 800 g for 10 minutes at 4 °C in a swinging bucket centrifuge (Eppendorf Centrifuge 5810R). The supernatant was removed, and cells were resuspended in 0.7 M D-mannitol (Sigma Aldrich M4125) in 1X PBS (gibco 70011-044) (PBS-M). This PBS-M buffer was optimized for scRNA-seq of cells adapted to high salinity environments, as documented in the separate technical paper^47^. Cells were then filtered through a 40 µm strainer once more to remove large clumps. The two filtration steps represent the only protocol changes between this May 2023 collection and the previously reported April 2022 collection.

Cell encapsulation, reverse transcription (RT), cDNA amplification, and library preparation was done with Chromium Next GEM Single Cell 3′ v3.1 Kits (10X Genomics 1000123, 1000127, and 1000190). We followed the manufacturer’s guidelines except for the preparation of the RT mix prior to cell encapsulation. The manufacturer’s protocol directs users to add nuclease-free water, followed by cell suspension; instead of adding nuclease-free water, we added an equivalent volume of PBS-M. scRNA-seq was provided by the Single Cell Core at Harvard Medical School, Boston, MA.

Both libraries were sequenced on an Illumina NovaSeq 6000, and 10X Genomics Cell Ranger 6.1.2 was used to generate a count matrix from FASTQ files. Reads were aligned to a reference genome which combined the KY21 gene models^77^ for the HT2019 assembly^78^ (downloaded from Ghost Database^79^) with mitochondrial genes from the Ensemble KH genome assembly^80^. To make it compatible with Cell Ranger, the KY21 gene model GFF3 file was converted to a GTF file using GffRead^81^. Genome files compatible with Cell Ranger for the combined genome are available at https://github.com/AllonKleinLab/paper-data/tree/master/Scully_mannitol_2025/C_robusta_genome_HT2019_KY21_with_Ensembl_mito

All further analyses were done on libraries from both experimental days: one library from April 2022 (from the technical paper^47^) and two from May 2023 (just described).

### Initial Cell Filtration

Cell barcodes corresponding to viable cells were filtered based on (i) the number of mRNA molecules detected with unique molecular identifiers (UMI), (ii) the fraction of mitochondrial reads, and (iii) the likelihood of being a doublet. For (i), cells were kept that had more than 2,000 UMIs. This threshold was set based on clear separation from background as seen by plotting a log-scale histogram of UMIs per cell barcode for each library. For (ii), a mitochondrial fraction below 10% was required. For (iii), we used the SCRUBLET package^82^ to identify likely doublets, which were removed.

### Processing and Dimensionality Reduction

Preprocessing and dimensionality reduction were performed with Scanpy^83^ version 1.8.1, using default parameters except where otherwise noted. Counts were normalized to the median counts per cell across all libraries (scanpy.pp.normalize_total). Highly variable genes were identified as described in Klein et al.^84^ (code available at https://github.com/AllonKleinLab/paper-data/blob/master/Scully_Ciona_blood_2025/helper_functions/highly_variable_genes.py). Counts were log transformed (scanpy.pp.log1p, base=10), z-scores were calculated (scanpy.pp.scale), then principal component (PC) analysis was performed (scanpy.tl.pca, use_highly_variable=True). Batch integration was performed using BBKNN^85^ to generate a single-cell network, treating the three libraries as the batches to be integrated (scanpy.external.pp.bbknn, batch_key=‘library’, neighbors_within_batch=3). This network was used for Leiden clustering of cells (scanpy.tl.leiden) and UMAP visualization (scanpy.tl.umap).

### SNP Genotyping and Demultiplexing

We assigned cells in each library to individual wild-caught animals using naturally present SNPs. Variant SNPs were identified using Cellsnp-lite^48^ (mode 2, i.e. without an a priori list of SNPs; cellsnp 0.3.2 and cellsnp-lite 1.2.3). Cell barcodes were then assigned to animals using Vireo^49^ (vireosnp 0.5.6). Initial inspection revealed problems with Vireo assignments for the April 2022 library, so we developed a custom pipeline. The inspection and new pipeline are described below.

### Quality checks for Vireo’s assignments

Initial inspection of Vireo assignments showed that some transcriptional clusters were called by Vireo as having a large proportion of between-genotype doublets. These clusters were abundant and showed unique gene expression, which is not consistent with them being doublets. Specifically, Vireo labeled over 50% of cells in four clusters as doublets in the April 2022 library (Supplementary Fig. 10a). Further inspection showed that Vireo made use of SNPs associated with genes expressed specifically in these clusters, which suggested that Vireo was assigning clusters within one animal to different SNP profiles. This problem may occur because some animals in the April 2022 collection contributed <2% of the initial blood sample by volume. For the two scRNA-seq libraries generated from the May 2023 collection, we did not observe similar problems and thus directly used the assignments provided by Vireo without using the custom pipeline described below.

### Custom SNP demultiplexing

To address the problem for the April 2022 library, we implemented a custom pre-processing step, which avoids focusing on SNPs that are cell type specific and instead uses SNPs broadly detected across cell states. To this end, we selected a subset of cells whose transcriptional clusters lack highly unique SNPs (cell barcodes listed in **Supplementary Table 8**). We carried out principal component analysis on the variable SNPs for this subset of cells, and then projected the SNP profiles for all cells onto the first principal components (PCs). The PC space of variable SNPs was generated by log-transforming the SNP count matrix for this cell subset (using scanpy.pp.log1p), then selecting highly variable SNPs (using scanpy.pp.highly_variable_genes, min_mean=0.0125, max_mean=3, min_disp=0.5), and running principal component analysis (scanpy.pp.pca, with default settings and use_highly_variable=True). PC1 separated the animal with the most cells from the rest (Supplementary Fig. 10b). We thus defined new animal assignments to be “animal 1” for cells with a PC1 value < 0.3, “animals 2–5” for cells with a PC1 value > 0.7, and “undetermined” for all other cells (Supplementary Fig. 10c). We did not fractionate cells from animals 2-5 as the cell count for these animals was low.

### Identification of cells from an animal contributing to more than one library

As noted above, one animal (animal F) contributed cells to both libraries in the May 2023 collection (see section ‘Single-Cell Barcoding, Library Preparation, and Sequencing’). To associate the cells from this animal in both libraries, Vireo-annotated animals were matched across libraries using vireoSNP.utils.vcf_utils.match_VCF_samples. A single SNP profile showed a close match between the libraries (Supplementary Fig. 10d), and was taken to be the shared animal.

### Identification and Removal of HA-3/SRC Doublets

We identified a subset of cell barcodes in the scRNA-seq dataset as doublets or multiplets of cells from the HA-3 and SRC clusters. Below, we describe (1) defining and removal of likely doublets, and (2) further the evidence that this subset represents doublets or multiplets.

## 1. Defining the cell subset coexpressing HA-3 and SRC marker genes

We found that Leiden cluster 19 (pre-doublet removal) comprises two populations—while all cluster 19 cells express markers of HA-3, a subset additionally expresses markers of SRC (corresponding to Leiden cluster 2 pre-doublet removal). We defined new cluster boundaries to reflect the presence of these two populations. For the cells in clusters 19 and 2, we categorized cells into 3 subsets: one expressing only HA-3 markers, one expressing only SRC markers, and one with co-expression of HA-3 and SRC markers.

Specifically, we focused on HA-3 marker KY21.Chr9.653 and SRC marker KY21.Chr11.830, which were used in HCR FISH experiments. We determined which cells coexpress these genes at the level of fine-grain clusters using the following approach. For Leiden clusters 2 and 19, we generated a new kNN graph (scanpy.pp.neighbors, n_neighbors=10), then assigned cells to fine-grain clusters with Leiden clustering at a high resolution (scanpy.tl.leiden, resolution=3). We classified each fine-grain cluster as “Chr9.653+” if they express Chr9.653 but not Chr11.830, as “Chr11.830+” if they express Chr11.830 but not Chr9.653, or as “double-positive” if they express both genes. Specifically, we considered a gene to be expressed in fine-grain cluster *F* if its mean log-transformed expression in cluster *F* is more than double the mean across cells in the full dataset which are not in Leiden clusters 2 or 19. The resulting redefinition of cell cluster boundaries are shown in Supplementary Fig. 11b (barcodes listed in **Supplementary Table 9**).

Cell barcodes of the “double-positive” population were removed, and we then repeated the post-filtration processing steps as described in “Processing and Dimensionality Reduction”. Fig. 1 shows regenerated Leiden clusters after doublet removal.

## 2. Evidence that cells coexpressing HA-3 and SRC marker genes are doublets

We expand here with a supplemental note giving three pieces of evidence suggesting that the “double-positive” cells are doublets or multiplets. (i) The genes enriched in these cells are all shared with Chr9.653+ (HA-3) or Chr11.830+ (SRC) cells (Supplementary Fig. 11e). (ii) The double-positive cells have a 1.95-times and 4.39-times higher mean number of transcripts detected per cell as compared to Chr9.653+ and Chr11.830+ cells, respectively (Supplementary Fig. 11f). (iii) HCR FISH staining revealed no cells that co-express Chr9.653 and Chr11.830, despite these genes being both detected in the double-positive population (p<0.05) (Supplementary Fig. 11g–h). The double-positive cells include cell barcodes from both experimental days (Supplementary Fig. 11c), so we speculate that it reflects a reproducible physical association of these two cell types.

### Manual Sub-clustering

After manual inspection of the Leiden clusters, we additionally sub-clustered Leiden clusters 2 and 14, due to within-cluster heterogeneity. In both cases, we split each of 2 and 14 into two by Leiden clustering with a resolution of 0.1; we called the bigger subclusters 2.1 and 14.1 and the smaller subclusters 2.2 and 14.2, respectively.

## Morphological Annotation

### Identification of Marker Genes for HCR FISH and Live Reporters

#### Filtering genes for HCR FISH

For choosing HCR FISH marker genes, we required that transcripts be long enough to accommodate 10 probes hybridizing to it. To allow 10 probe pairs of 53 nucleotides each, with a spacing of at least 10 nucleotides between each pair, transcripts must be at least 610 nucleotides long. We only considered marker genes from the KY21 gene models^77^ whose shortest annotated transcript was at least 610 nucleotides in length.

#### Filtering genes for live reporters

For choosing live reporter marker genes, we wanted to avoid genes which might be expressed in the developmental precursors of blood cells; if GFP protein remained in these cells’ descendents, GFP+ blood cells could include cells not currently expressing the marker gene. Where possible, we selected genes with low or no expression in scRNA-seq data of *C. robusta* developmental mesenchyme^86^. Specifically, when possible, we chose genes with log-transformed expression log_10_(CP10k +1) below 0.1 in all mesenchyme clusters.

#### Selecting marker genes

Marker genes for each cluster were selected to (i) be expressed highly enough to be detectable with HCR FISH or live reporters; (ii) show a large dynamic range with near-binary in expression, such that cells can be easily categorized as expressing or not expressing the gene; and (iii) be specific to each cluster.

To achieve (i), we ran preliminary experiments labeling genes with a range of expression levels to identify the detection limit. In the following, log-transformed expression refers specifically to log_10_((*counts per M*) + 1), where *M*=6,138 is the median unnormalized counts per cell across the dataset, evaluated before doublet cluster removal. In preliminary HCR FISH experiments, we found genes with a mean log-transformed expression below 1 were not detectable, but genes with expression above 2 were easily detected: KY21.Chr3.306 (mean log expression of 0.70 in GA) was not detected; Chr8.361 (1.6 in HA-3) was weakly detected; and all of Chr13.352 (2.8 in GA), Chr9.653 (3.5 in HA-3), and Chr12.617 (4.2 in GA) were easily detected. Based on these experiments, we required that candidate marker genes for each cluster of interest have a mean log-transformed expression above 1 in that cluster.

For (ii), we defined a score that we reasoned would identify genes that appear binary by HCR FISH as follows. Let *r* _*u*_: = *min* {*x* ɛ*X*: *x* > *u* } − *max* {*x* ɛ*X*: *x* < *u* }, where *X* = {*x*, *x* _2_,…, *x* _N_} and *x* _k_ is the log-transformed mean expression of the gene in cluster *k*. The value of *r* _*u* =1_ gives the log-ratio in expression between clusters above and below the detection limit (*u*=1). With this score, marker genes for each cluster were selected by first filtering genes to consider only those with mean log expression >1 in at least one cluster and <0.35 in at least one other, then further filtering genes with mean log expression above 1 in the cluster, and finally selecting the top several genes with highest *r* _*u* =1_ values. Finally, we carried out a manual inspection of expression of these genes on the UMAP embeddings, to select 1–2 genes with highly specific expression for each cluster.

In some cases, no genes passing these filters uniquely labeled a cluster. For cluster pairs 1/7 and 8/26, which are transcriptionally similar, we chose marker genes which labeled both clusters together (Supplementary Fig. 3-2b**, 3-5b**). For cluster pair 0/14, also transcriptionally similar, no filtered genes labeled only 0/14 and no other clusters; we chose two genes which are only coexpressed in 0/14 (Supplementary Fig. 3-1b). The selected marker genes are listed in Supplementary Fig. 2c (HCR FISH) and Supplementary Fig. 5a (live reporters).

#### Selection of Ubiquitous Control Gene

We selected a ubiquitously expressed gene to be used as a positive control in HCR FISH experiments. We required that such a gene have a mean *log* _10_ (*CP* 10*k* + 1) above 1 in at least 30 out of 33 Leiden clusters. Four genes pass this threshold, from which we chose KY21.Chr11.687 (homologous to *ACTB/G1*) as a control.

#### Probe Design for HCR FISH

Once genes were selected for HCR FISH experiments, HCR amplifier sequences^51,87^ (B1, B2, or B3) were chosen for each gene. These sequences determine which fluorescent hairpins hybridize to a gene’s probe set. The ubiquitous control gene KY21.Chr11.687 was given amplifier B2, and all cluster marker genes were given either B1 or B3 as reflected in **Supplementary Table 10**. Probe sequences (split initiator probe pairs) were designed for each gene and HCR amplifier sequence as described by Choi et al.^51^. Between 10 and 20 probe pairs were evenly spaced across each gene’s longest transcript sequence (excluding introns, including untranslated regions), with a minimum of 10 nucleotides between probe pairs. Sequences for each gene can be found in **Supplementary Table 10**. Probes were ordered from Integrated DNA Technologies.

#### Cell Preparation for HCR FISH with Live Imaging

In preparation for cell imaging, 8-well polymer coverslip pretreated with poly-L-lysine (PLL) (ibidi 80824) were coated with concanavalin A as follows: 150 µL of 1 mg/mL concanavalin A (ConA) (Sigma C2010-1G) in nuclease free water (Invitrogen 10977-023) was added to each well to coat the bottom for at least 90 minutes or overnight; the solution was then removed, and wells were left to dry for at least 3 hours or overnight. The coverslips were then used within <1 day.

Blood cells were then collected as described above (“Blood cell collection”), with 4–5 animals contributing cells for each experiment. After collection, blood samples from all animals were kept on ice, then pooled and filtered through a 40 µm strainer (pluriSelect 43-1004-60). The cell concentration was estimated using the BioRad TC20 Automated Cell Counter, then the suspension was diluted with Ca^2+^- and Mg^2+^-free artificial seawater (CMF-ASW; 0.5 M NaCl, 9 mM KCl, 5 mM HEPES, pH 7.4) to a concentration of 2e6 cells/mL. DAPI (Sigma Aldrich D9542) was added to a concentration of 3 µM for identification of dead cells during subsequent imaging. Next, 150 µL of cell suspension was added to each well of the pre-treated 8-well polymer PLL+ConA coated coverslips. The coverslips were then taped to the bottom of a swinging bucket centrifuge and spun at 400g for 10 minutes at 4 °C to precipitate cells.

The coverslip wells containing live cells were then imaged on a widefield Nikon Ti2 inverted microscope with a Lumencor Sola light engine, the Perfect Focus System, and a Nikon Plan Apo VC 100x Oil 1.40 NA objective. Morphological information was collected using DIC imaging, with color information collected by illuminating cells with red, green, and blue light in series.

DAPI signal was collected using excitation filter Chroma ET395/25x, dichroic mirror Chroma T425lpxr, and emission filter Chroma ET460/50m. Images were collected using a Hamamatsu Flash4.0 LT camera (6.5 µm^2^ photodiode) with NIS-Elements image acquisition software.

Imaging was carried out at room temperature, and a 9 by 9 grid of images was captured for each sample using a Nikon motorized xy-stage. Immediately after each well was imaged, cells were fixed by gently adding 150 µL of 8% paraformaldehyde (PFA, Electron Microscopy Sciences 15710) in CMF-ASW (final concentration 4% PFA in 300 µL). After a 30–45 min incubation at room temperature, PFA was washed twice with 150 µL of 1X PBS (gibco 70011-044). Fixed and washed cells were imaged once more across the same fields of view. Across all samples on all days, 60–75 minutes passed between the first animal’s blood collection and cell suspension centrifugation onto the coverslip, and 45–90 minutes passed between centrifugation and the start of PFA fixation.

For most samples, we proceeded immediately to HCR FISH (see next section) on the same day. For labeling markers of clusters 0/14, 8/25, 10, 11, 12, 13, 22, and 24, we dehydrated cells with methanol and stored them at −20 °C before beginning HCR FISH. We found that methanol dehydration did not affect the quality of *in situ* hybridization in these cells, so this was used as a convenience to split experiments across multiple days when needed. Cells were dehydrated by washing three times for 5 minutes in 150 µL of 100% methanol chilled to −20 °C (OmniSolv MX0484-1). Dehydrated cells were stored overnight or up to 5 days at −20 °C.

#### HCR FISH

We followed an HCR FISH protocol adapted from previously published protocols^51,88^. Unless otherwise stated, a volume of 150 µL was used for all washes and incubations. For dehydrated samples, cells were rehydrated with a series of washes: 5 minutes with 75% methanol in PBST (1X PBS with 0.1% Tween-20 (Millipore Sigma P9416-50ML)), 5 minutes with 50% methanol in PBST, 5 minutes with 25% methanol in PBST, and then 5 5-minute washes in PBST. All samples were permeabilized with a 5–7 minute wash in 1X PBS with 0.1% Triton X-100 (VWR VW3929-2), followed by a wash in PBST.

Next, cDNA probes were hybridized to target mRNA in each sample. Before hybridization, samples were incubated in probe hybridization buffer (Molecular Instruments) for 30 minutes at 37 °C. Then hybridization began by replacing the buffer with probe solution (4 nM concentration of each gene’s probe set in probe hybridization buffer) which had been preheated to 37 °C. Samples were incubated with probes for 20-25 hours at 37 °C. Probes were then removed by washing 4 times for 15 minutes at 37 °C with probe wash buffer (Molecular Instruments), then washing 2 times for 5 minutes at room temperature with 5X SSCT buffer (5X SSC buffer (Invitrogen AM9763) with 0.1% Tween-20).

Fluorescence signal was then added and amplified using metastable fluorescent hairpins^51^. These fluorescent hairpins were first snap-cooled by heating to 95 °C for 5 minutes, then cooling to room temperature for 30 min. A hairpin solution was prepared by adding snap-cooled hairpins to amplification buffer (Molecular Instruments) at a ratio of 1:50 for each hairpin. See **Supplementary Table 11** for the hairpins used for each transcriptional cluster. Amplification was done at room temperature: samples were first incubated in 5X SSCT buffer for 30 min, then incubated in the hairpin solution for 14.5–17.5 hours. Excess hairpins were removed by washing samples 5 times for 5 minutes in 5X SSCT buffer at room temperature. Finally, cells were labeled with DAPI by washing with PBST, incubating with 3 µM DAPI in PBST for 20–35 minutes, then washing twice more for 3 minutes with PBST. Samples were stored at 4 °C.

HCR FISH and DAPI fluorescence was imaged using a Yokogawa W1 spinning disk confocal on an Nikon Ti inverted microscope with the Perfect Focus System and a Nikon Plan Apo Lambda D 100x Oil 1.45 NA objective. Fluorescence was excited and collected with the lasers, dichroic mirrors, and emission filters listed in **Supplementary Table 12**. Images were acquired using a Hamamatsu ORCA-Fusion BT CMOS camera (6.5 µm^2^ photodiode) with NIS-Elements image acquisition software. The same fields of view were imaged as for the live imaging. To achieve this, at the start of the experiment, one well on each 8-well coverslip was left empty, and an “X” was etched into the center using a diamond pen. Coordinates for each field of view were saved relative to the center of this X during live imaging, then used to find the same fields of view during HCR FISH imaging using a Prior Proscan III motorized stage. At each field of view, 9 z-slices were collected with a step size of 1 µm. DIC images were additionally captured at each field of view to aid in matching cells across rounds of imaging. Fluorescence z-stacks are displayed as maximum-intensity z-projections, and contrast limits were adjusted (identically for all cells within a sample) using a custom image analysis pipeline built in Python using napari^89^ (available at https://github.com/AllonKleinLab/paper-data/tree/master/Scully_Ciona_blood_2025/HCR_FISH/napari_image_analysis_pipeline). Using this pipeline, cells were manually matched between the live images and HCR FISH fluorescence images to link marker-positive cells to live morphologies (Supplementary Fig. 2e).

We found that some cells tended to non-specifically accumulate HCR FISH signal (example in Supplementary Fig. 2e, grey arrow). We showed that this fluorescence was non-specific by using a probe set of zebrafish col5a1, which still labeled some cells (Supplementary Fig. 12). This non-specific fluorescence was distinguishable from FISH-labeled mRNA in that it (1) was much brighter than HCR FISH signal, (2) tended not to have puncta but rather to have a large uniform region of fluorescence, and (3) was fully overlapping in all channels. Fluorescence matching these features was ignored when identifying marker-positive cells.

## Live *in vivo* Imaging

### Molecular Cloning

To construct reporter transgenes, HMW *C. robusta* genomic DNA was PCR-amplified with oligos listed in Supplementary Table 13. Putative enhancer fragments were restriction-digested and subcloned into *Mesp>GFP* (described by Davidson et al.^90^) after restriction digest removal of the *Mesp* enhancer fragment. All transgenes were sequenced prior to use. *Mesp>RFP* was described previously by Davidson et al.^91^.

### Electroporation

Electroporation procedures followed the method described by Zeller et al.^92^. Transgene DNAs in water were combined and diluted in 0.77M D-mannitol not exceeding a final concentration of 125ug/mL. Embryos and juveniles were subsequently cultured at 18 °C in 0.22 μm-filtered artificial seawater (FASW) supplemented with penicillin (10 U/mL) and streptomycin (10 μg/mL).

### Imaging

For *in vivo* imaging, transgenic *C. robusta* juveniles were paralyzed by incubation in menthol-treated FASW. Samples were imaged in menthol-treated FASW in glass-bottomed dishes. For *ex vivo* imaging, transgenic juvenile *C. robusta* in FASW were roughly dissociated using a plastic tube pestle. Samples were mounted on slides. Images were captured on a Zeiss LSM 980 via confocal imaging or fluorescent widefield with an Axiocam 305 camera.

## Characterization of Cell Function

### Identification of *C. robusta* Vertebrate Homologs with OrthoFinder

We used OrthoFinder^65^ (version 2.5.5) to identify likely vertebrate homologs of *C. robusta* genes. OrthoFinder was run on the genomes with species, source repositories and versions listed in **Supplementary Table 14**, with default parameters. If genes from two species are in the same orthogroup as defined by OrthoFinder, they are considered homologous.

### Gene Ontology Enrichment Analysis

For **Extended Data** Figs. 6b and 7b, Gene Ontology (GO) enrichment analysis was carried out to identify GO terms which were significantly enriched in each cell type’s list of differentially expressed genes (DEGs). DEGs for each cell type were determined using the Wilcoxon rank-sum test (scanpy.tl.rank_genes_groups, method=’wilcoxon’, requiring log2 fold change > 1 and FDR<0.01). Lists of *C. robusta* genes associated with each GO term were generated based on their homology to human genes associated with that GO term, as described below. Fisher’s exact test was then used to identify GO terms which were significantly enriched in each cell type’s DEG list. Code for this enrichment analysis is available at https://github.com/AllonKleinLab/paper-data/tree/master/Scully_Ciona_blood_2025/scRNA-seq_analysis/go_enrichment_analysis.

For each GO term, we defined the *C. robusta* gene list as the set of *C. robusta* genes which have at least one human homolog (defined by OrthoFinder) in that GO term’s human gene list. We avoided GO terms represented by small numbers of genes by only testing enrichment of GO terms with both of the following: (i) at least 10 *C. robusta* genes in the final gene list, and (ii) at least 10 distinct human genes with a homolog in *C. robusta* in the original human list. For performing Fisher’s exact test, we considered only *C. robusta* genes which were detected in more than 20 cells in the scRNA-seq data. Genes expressed in fewer cells were removed from cell DEG and GO term lists. Fisher’s exact test was then run using scipy.stats.fisher_exact (alternative=‘greater’). Multiple hypothesis correction was performed using the Benjamini-Hochberg procedure, requiring a false discovery rate (FDR) below 0.05 (statsmodels.stats.multitest.fdrcorrection, alpha=0.05).

### RNA Velocity Analysis

RNA velocity^53^ was used to generate hypotheses of differentiation trajectories. Velocyto^53^ (version 0.17.17) was used to quantify spliced and unspliced expression levels for each gene. We then used scVelo^93^ (version 0.3.0) to compute a splice-aware neighbors graph of all cells and estimate velocities for all genes. These were used to compute a velocity graph whereby the transition probability between pairs of cells was estimated. This graph was overlaid on the pre-existing UMAP coordinates of the *C. robusta* barcoded cells (Fig. 4b). Arrows pointing away from the cMPP cluster were manually selected and differently colored after vector image generation.

### Differentiation Hierarchy Analysis

To build a tree connecting cell types in a differentiation hierarchy (Fig. 4c), we adapted a method in Wagner et al. 2018^54^. The tree is constructed as a maximum spanning tree (MST, implemented with networkx.maximum_spanning_tree^94^) on an undirected, weighted cell type graph. The graph is constructed with edge-weights biased towards connections between cycling cell clusters (assumed to be progenitors) and non-cycling clusters (assumed to be differentiated non-progenitors). The bias is achieved by combining two graphs: a graph connecting all clusters to all clusters, and a bipartite graph connecting progenitors to non-progenitors. This biased approach enables the use of different parameters (thresholds on edge weights) for progenitor and non-progenitor clusters, as we observed that there is otherwise a dominance of connections within progenitor clusters. The method is detailed below, with Python code implementing the graph construction available at https://github.com/AllonKleinLab/paper-data/tree/master/Scully_Ciona_blood_2025/scRNA-seq_analysis/differentiation_hierarchy_analysis. The final tree was visualized in Fig 4c using Graphviz’s neato spring model layout^95^, with some node positions manually adjusted to improve readability.

For building the differentiation hierarchy, we used cell type labels as defined in **Supplementary Table 2**. We built the maximum spanning tree, *MST* (*G*), of graph *G* = (*V*, *E*, *W*), where vertices *V* are the set of cell clusters; edges *E* are the set of all possible edges, making a fully connected graph; and weights *W* reflect the connections between cell clusters. For simplicity, we omit *E* from further definitions. This graph *G* = (*V*, *W*) was constructed from two other graphs: (1) an *G* _(0)_ = (*V*, *W* _(pd)_), and (2) a bipartite graph *G* _pd_ = (*V*, *W* _p,d_) of progenitor-to-differentiated cell cluster connections. Within *G*, the weight between cell clusters *i* and *j* was defined as *w_ij_ = max(w^(0)^_ij_, w^(pd)^_ij_)*.

We now define the construction of (0) and (pd) below. For each, we used the batch-corrected k-nearest neighbors (*k*NN) graph on the scRNA-seq data generated by BBKNN (see section “Processing and Dimensionality Reduction”). (1) To build (0), we established an undirected coarse-grained graph between annotated cell clusters as follows: Within the cell *k*NN graph, let *n*(*i*, *j*) be the number of edges connecting individual cells between cell cluster *i* and cell cluster *j*, where *i* ≠ *j*. Let *N*(*i*) = ∑ *n*(*i*, *k*) be the total number of outgoing edges from cell cluster *i*. We define a coarse-grained graph as having one node per cell cluster, with edge weights, *w_i,j_*, between cell cluster *i* and *j* given by:

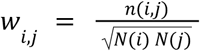

We then excluded edge weights which are less than 30% of the maximum edge weight to obtain final edge weights: *w_i,j_*_p,d_ = {*w_i,j_* for *w_i,j_*_p,d_ > α max (*w_i,j_*_p,d_); 0 otherwise}, where α = 0. 3. (2) We built *G*, a bipartite graph connecting progenitor to non-progenitor cell clusters, as follows: We labeled each cell cluster as a progenitor or non-progenitor cluster based on expression of cell cycle markers. Specifically, cell clusters were labeled as progenitors if their median expression of the *MKI67* ortholog, KY21.Chr2.1280, was above zero; all other cell clusters were labeled as non-progenitors. For building this bipartite graph ^(pd)^, we only considered edges in the cell *k*NN graph which connected a progenitor to a non-progenitor. The process for determining edge weights otherwise followed the process for building *G*. Note that since only a subset of edges from the kNN graph are used for building *G*, the values of *n*(*i*, *j*) and *N*(*i*) are different from those used for building ^(0)^. The same value α = 0. 3 was used.

### Zymosan Phagocytosis Assay

To test phagocytic function of clusters HA-1 and HA-2, we incubated cells with zymosan, a yeast cell wall extract which is readily engulfed by phagocytes^96^. The cells were then fixed and stained by HCR FISH with markers of HA-1/2. pH-sensitive (pHRhodo) dye–labeled zymosan was provided as a kind gift from Lillian Horin. It was prepared by washing unlabeled zymosan (Invivogen tlrl-zy) 3 times with 0.1 M sodium bicarbonate (Sigma Aldrich S6014-1KG), then incubating the pellet with 200 µM pHrodo amine-reactive dye (Invitrogen P36011) in PBS with 0.02% sodium azide for 30 minutes, rocking at room temperature. The zymosan was then washed 5 more times with PBS with 0.02% sodium azide, then stored for 3 years at 30 mg/mL in PBS with 0.02% sodium azide at 4 °C. It was diluted in CMF-ASW to a concentration of 1 mg/mL shortly before being added to cells.

Next, blood was collected as described above (“Blood Cell Collection”), then mixed with diluted zymosan in a ratio of 9:1 blood:zymosan by volume. The cells were then incubated for 2 hours at 18 °C in a rocking microcentrifuge tube, followed by filtration through an 40 µm strainer and centrifugation onto a PLL+ConA coated coverslip for live imaging as described above (“Cell Preparation for HCR FISH with Live Imaging”). We confirmed that intracellular zymosan, but not extracellular zymosan, was fluorescent (Supplementary Fig. 7c). We then proceeded to label HA-1 and HA-2 marker genes as described in “HCR FISH” (**Supplementary Table 11**).

### Identification of *C. robusta* Likely Human Homologs with BLAST

For gene expression analysis presented in Fig. 4h and Supplementary Fig. 6e–g, we plot the expression patterns of immune-related genes across *Ciona robusta* cell types. To identify homologous *C. robusta* genes corresponding to human genes of interest (detailed in **Supplementary Table 15**), we employed a BLAST-based approach rather than relying solely on OrthoFinder results. This methodological choice was necessitated by OrthoFinder’s failure to identify several known immune-related homologs in *C. robusta*, including previously documented Toll-like receptor (TLR) homologs^75^. The BLAST-based approach enabled a more comprehensive identification of functional homologs between human and *C. robusta* genomes for subsequent expression analysis.

Our BLAST-based homology detection pipeline accounted for vertebrate- and tunicate-specific gene duplication events by allowing one-to-many gene matching. Specifically, we identified each *C. robusta* gene’s best human gene hit based on BLASTp (aligning amino acid sequences, see **Supplementary Table 14** for proteome versions; BLAST+^97^ version 2.12.0). To ensure specificity and prevent spurious matches, we filtered the results to exclude cases where the identified human hit had substantially closer sequence similarity to a different *C. robusta* gene, specifically: for each *C. robusta* gene *g*^*Cr*^ with best human hit *g*^Hs^, the gene pair (*g*^*Cr*^, *g*^Hs^) was excluded if the bit score between ^*Cr*^ and ^Hs^ was less than half the bit score between ^Hs^ and its best *C. robusta* hit.

## Comparison to Human Hematopoiesis

### Comparison of Cell Types using SAMap

We ran pairwise SAMap twice: once to compare human (*Homo sapiens*) to zebrafish (*Danio rerio*), and once to compare human to *C. robusta*. For the human dataset we used the Tabula Sapiens blood and bone marrow data^64^ (TS_Blood.h5ad.zip and TS_Bone_Marrow.h5ad.zip from https://figshare.com/articles/dataset/Tabula_Sapiens_release_1_0/14267219). To improve the interpretability of results, we combined similar human cell type annotations and performed most analyses using this coarse-grain cell clustering (**Supplementary Table 16**). For the zebrafish dataset, we used data of zebrafish kidney marrow from Wang et al.^68^ (Genome Sequence Archive CRA017585). The cells for this dataset were manually re-annotated by inspecting gene expression, and new annotations for each cell barcode are listed in **Supplementary Table 17**. Some cell barcodes were identified as likely kidney progenitors and were excluded from analysis.

SAMap was run in Python3 (version 1.0.15) with samap.run (pairwise=True). Similarity matrices were generated with samap.analysis.get_mapping_scores (n_top=0). To display similarity matrices, one species’ cell types were ordered from highest to lowest similarity score. For the second species, we used the following ordering procedure: For each cell type *i* in the ordered list of the first species, we identified all cell types from the second species that had their maximum similarity with cell type *i*. These matching cell types were then added to the ordered list of the second species, arranged from highest to lowest maximum similarity score.

### Jaccard Similarity Score

We define a cell type similarity score based on the Jaccard index to quantify the fraction of cell type DEGs which overlap between two species. Let *A* be the set of DEGS in cell type 1 of species 1, excluding genes with no homolog in species 2. Let *B* be the set of DEGs in cell type 2 of species 2, excluding genes with no homolog in species 1. Given the many-to-many gene homology, we calculate the intersection of *A* and *B* in two ways: We define {*A* ∩ *B*} as the set of species 1 genes in *A* with at least one homologous gene in *B*, and {*A* ∩ *B*} as the set of species 2 genes in *B* with at least one homologous gene in *A*. We then calculate two Jaccard indexes, 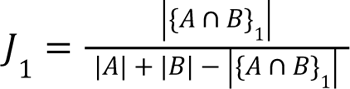 and 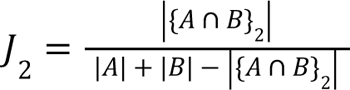
. The Jaccard similarity score is defined as the mean of these two values: J = ½ (*J*_1_ + *J*_2_).

We identified DEGs for each cell type in the human, zebrafish, and *C. robusta* datasets using the following approach. For the human and zebrafish data, we normalized the count matrix to 1e4 counts per cell (scanpy.pp.normalize_total, target_sum=10^4) and log-transformed the data (scanpy.pp.log1p, base=10). For all three species datasets, we identified enriched DEGs using the Wilcoxon rank-sum test (scanpy.tl.rank_genes_groups, method=’wilcoxon’), keeping genes with a log2 fold change > 2 and a false discovery rate < 0.05.

## Generalized Sankey (GS) Plots

### Definition of GS plots

GS plots are a generalization of Sankey plots, which visualize flows between nodes of a directed graph. A canonical Sankey plot visualizes incompressible flows, and the objects flowing between nodes are not individually identifiable. GS plots make two generalizations: they allow flows to be compressible (expanding or contracting), and they trace flow of identifiable elements between nodes. In a limit of incompressible flow and arbitrary element identity, a GS plot is a Sankey plot.

In Sankey and GS plots, graph nodes are represented by rectangles, and graph edges are represented by ribbon-like connectors^98^. Canonical Sankey plots visualize flow magnitude by the widths of connectors. Flows do not overlap, and the sum of connector widths into and out of a node are constant to reflect incombressible flow. For the general case represented by GS plots (Supplementary Fig. 8b), the underlying flows can overlap when the same elements (in our case, genes) from one node (in our case, a cell type) flow to elements in more than one node.

Second, compressible flows are represented by connector widths changing between start and end. Third, we consider the case in which nodes contain additional elements not present in the flows; graphically, this is represented by allowing node rectangle widths to exceed the total width of incoming flows.

In the following, we define GS plots for the case of a directed graph with two layers (two species, in our case), rooted in a single node (one cell type, in our case) in one layer, which flows to one or more nodes in a second layer (multiple cell types). The code for making these plots is available on Github (https://github.com/AllonKleinLab/paper-data/tree/master/Scully_Ciona_blood_2025/species_comparisons/gs_plots). These GS plots span three levels of ontology: two layers, which contain one or more nodes, which contain multiple elements. Elements may have many-to-many relationships across layers. The GS plots compare a focus node (FN) from one layer (Layer A) to a set of nodes from another layer (Layer B). Each node contains a number of elements and is represented by a rectangle whose width corresponds to the number of those contained elements.

GS plots use ribbon-like flows whose widths represent the number of shared elements between the FN in Layer A and each node in Layer B. For the flows between two nodes, the width where a flow meets the top (FN) rectangle represents the number of the FN’s elements which are related to at least one element in the connected bottom node; the width at the bottom represents the number of the bottom node’s elements which are related to at least one element in the FN. Since we allow many-to-many element relationships, the flows’ widths at the top and bottom may not be the same. Flows are plotted to overlap in the FN when the same elements are present in multiple categories of Layer B. The flows are specifically rendered as the area between two Bézier curves.

### Application of GS plots to cross-species comparison (Fig. 5c,d)

To use GS plots in cross-species comparisons (Fig. 5c**,d**), each layer represents a species, nodes represent cell types, and elements represent enriched DEGs. DEGs were determined as described for the Jaccard similarity score calculation (see “Jaccard Similarity Score”).

Many-to-many homologous relationships between genes were determined by OrthoFinder (see “Identification of *C. robusta* Vertebrate Homologs with OrthoFinder”). For each cell type, we additionally represented genes with no homology (i.e. node elements which can never be part of flows) through rectangle coloration: filled-in regions represent the number of genes which have at least one homolog in the other species’ genome, and empty regions represent genes with no detected homolog.

When visualizing relationships between a focus cell type (FCT) and cell types from another species, we selected the five cell types with the highest Jaccard similarity scores, excluding cell types with fewer than 10 shared genes. Cell types were arranged from left to right in order of decreasing Jaccard similarity score.

### Lineage-Enriched Transcription Factor Analysis

We first generated a list of *C. robusta* TFs by combining a curated list of *C. robusta* TFs from the Ghost Database^79^ with 70 additional genes which OrthoFinder mapped to known human TFs^99^. **Supplementary Table 6** contains the full list of 381 *C. robusta* TFs, along with names used in this paper. Next, using the coarse-grain differentiation hierarchy tree (Fig. 4c), we found TFs which might be involved in fate decisions by identifying differentially expressed TFs at each branching point (Fig. 6c). There are three branch points in the tree (i.e. nodes with degree > 2): cMPP, cLRP-2, and cLRP-4. At each of these branch points, we identified differentially expressed TFs across the subset of cell clusters which are child nodes of the branching point (e.g. for the branch point cLRP-3, we compared cLRP-5, pre-RA, RC, and URG-1). Specifically, we used the Wilcoxon rank-sum test (scanpy.rank_genes_groups, method=‘wilcoxon’, corr_method=‘benjamini-hochberg’) to compare TF expression in each cell cluster to other cell clusters in the subset. TFs with a log fold change > 2 and a false discovery rate < 0.01 were labeled as enriched in the given cell cluster and are plotted in Fig. 6c.

### Co-expression Conservation

Conservation of gene co-expression relationships was calculated following methods established in Crow et al.^71^ with one minor modification. For completeness we describe the approach here. We first built a gene co-expression network within each species (*C. robusta*, human, and zebrafish) as follows: the scRNA-seq data was clustered at a high resolution (scanpy.tl.leiden, resolution=20), and then for all pairs of genes we calculated the Spearman correlation of mean log-transformed expression across clusters. These correlation coefficients were ranked (descending) across all gene pairs to generate a weighted graph of gene co-expression relationships. We next quantified gene co-expression conservation based on how the networks centered around each gene compares across species using the “AUROC score” described in Crow et al.^71^, allowing *N*-to*-M* homology. Gene homology was determined using OrthoFinder. This approach recapitulates the method in Crow et al.^71^, with only a difference in the original high-resolution clustering step.

As a complementary approach, we calculated a second conservation score, which shows similar results (Fig. 6g, Supplementary Fig. 9). The second score, labeled as the “number of shared co-expression partners”, represents the number of co-expression partners shared across species, and was calculated as follows. We compared a human gene of interest (*g*_ℎ_) to one homologous gene (*g*_*x*_) in a non-human species (*C. robusta* or zebrafish). Taking the top 30 human co-expression partners of *g*_ℎ_ (the 30 genes with the largest edge weights connected to *g*_ℎ_), we identified the non-human species’ genes which are homologous to at least one human co-expression partner. There were more or fewer than 30 homologous genes in cases of *N*-to-*M* homology. We then defined the score as the number of these co-expression partner homologs which are highly correlated with *g*_*x*_, specifically the number of these genes whose Pearson correlation with *g*_*x*_ is above 0.5.

For cases where the genes of interest *g*_ℎ_ or *g*_*x*_ have more than one homolog in the other species, we calculated separate co-expression conservation scores for all homologous gene pairs. For example, human *IRF4* and *IRF8* are both homologous to *C. robusta Irf-r.b*, so we calculated two conservation scores—one each for *IRF4* vs. *Irf-r.b* and *IRF8* vs. *Irf-r.b* (Fig. 6g).

## SUPPLEMENTARY INFORMATION

### Supplementary Figures

**Supplementary Fig. 1:**
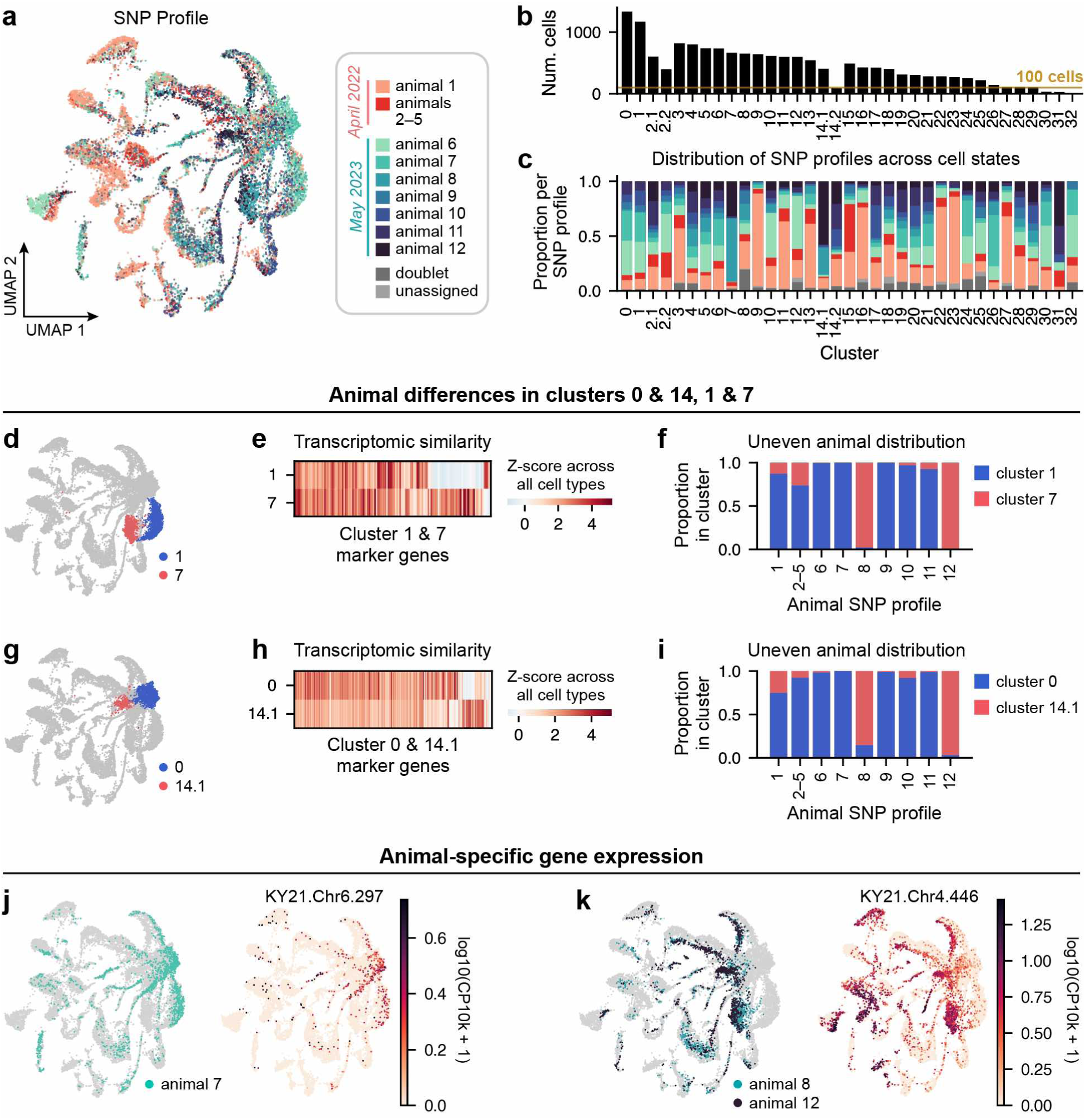
Gene expression differences across animals **(a)** UMAP with cells colored by SNP profile. **(b)** The number of cells per Leiden cluster. Of the five clusters detected in fewer than 7/9 SNP profiles (Fig. 1f), two—clusters 31 and 32—have fewer than 100 cells detected (yellow line). We would not expect to observe these rare cell states in animals with fewer cells sampled. **(c)** The fraction of each SNP profile contributing to each Leiden cluster. **(d–i)** Clusters **(d–f)** 1 and 7, as well as **(g–i)** 0 and 14.1 are shown in **(d)** and **(g)**. **(e, h)** Heatmaps of enriched genes (Wilcoxon rank-sum test, log fold-change > 2, p-value<0.05) show that clusters in each pair are transcriptionally similar. **(f, i)** Clusters 7 and 14.1 are almost entirely composed of cells from animals 8 and 12, while clusters 1 and 0 contain cells from the other SNP profiles. **(j, k)** At least some of the transcriptional differences between cluster pairs 1 vs. 7 and 0 vs. 14.1 are driven by genes which are broadly upregulated in specific animals. For example, **(j)** KY21.Chr6.297 is broadly upregulated in animal 7, and is expressed in clusters 0 and 1 but not 14.1 and 7. **(k)** KY21.Chr4.446 is broadly upregulated in animals 8 and 12, and is expressed in clusters 14.1 and 7 but not 0 and 1.

**Supplementary Fig. 2:**
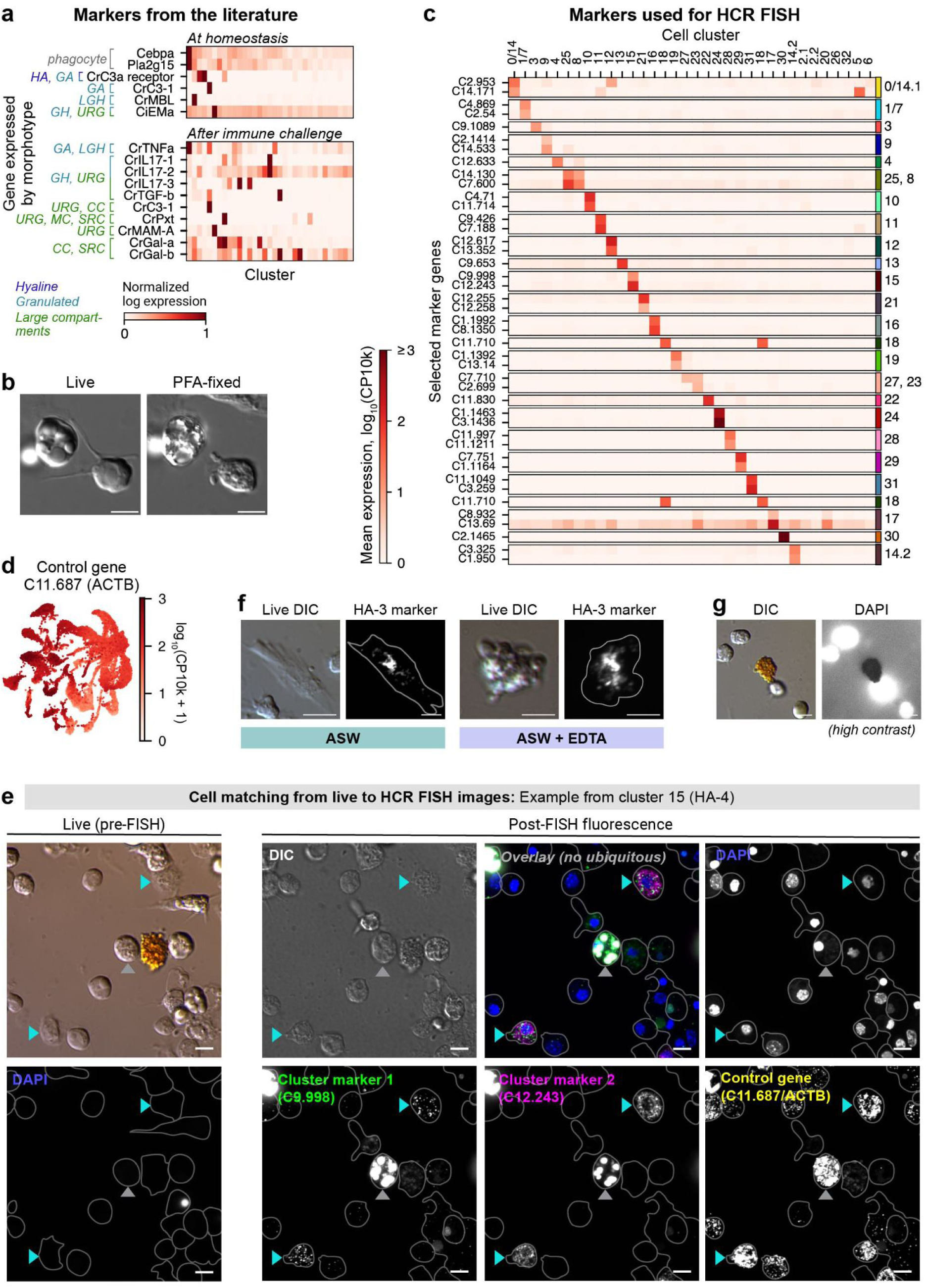
**(a)** Previously identified morphotype marker genes. Markers are separated based on whether they are expressed at homeostasis vs. after injection with lipopolysaccharide. See **Supplementary Table 1** for gene IDs and citations. **(b)** Image of cells immediately before and after fixation with PFA, showing fixation-induced morphological changes. **(c)** Cluster-specific marker genes chosen for HCR FISH experiments. See also **Supplementary Table 10**. **(d)** Ubiquitously expressed control gene chosen for HCR FISH experiments. **(e)** Example field of view from cluster 15 (HA-4) showing how cells were matched between live and post-FISH rounds of imaging. Teal arrows point to double-marker-positive cells. Grey arrows point to a cell with non-specific fluorescence (see Supplementary Fig. 12). **(f)** Cells positive for an HA-3 marker in two buffer conditions (ASW vs. ASW with 10 mM EDTA (ethylenediaminetetraacetic acid, Avantor EM-4050)) have dramatically different live morphologies. **(g)** Live orange cells block fluorescence in the DAPI channels. Image shows live cells labeled by DAPI. Contrast limits have been adjusted to show that background fluorescence is blocked by the orange cell. All scale bars are 5 µm.

**Supplementary Fig. 3:**
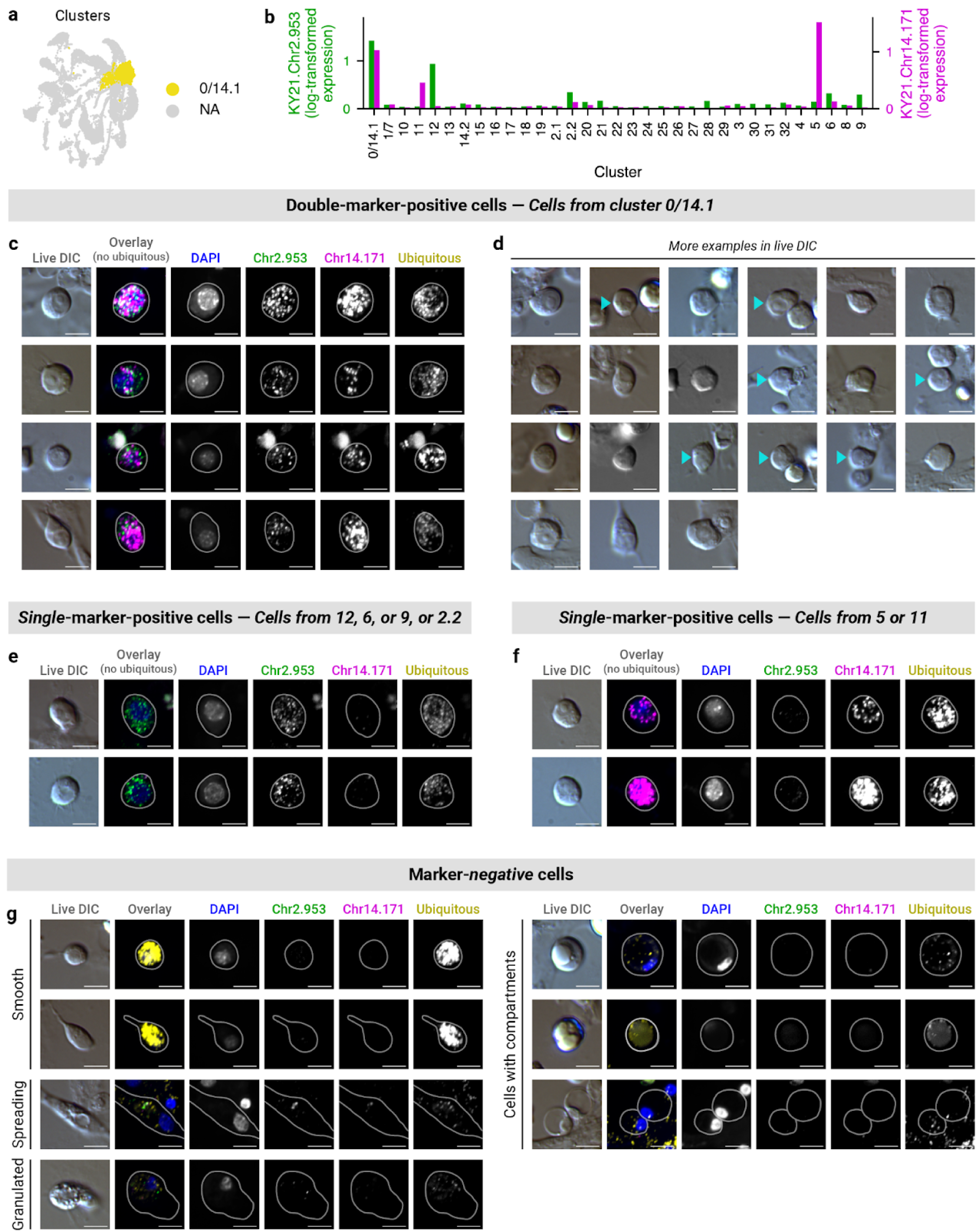
HCR FISH results by cell cluster. *3-1*: Cluster 0/14.1 (cMPP) **(a)** UMAP showing cluster 0/14.1. **(b)** Bar plot showing marker gene expression (log_10_(CP10k + 1)), only coexpressed in 0/14.1. **(c, d)** Example double-marker-positive cell images, shown **(c)** with or **(d)** without fluorescence. Teal arrows indicate double-marker-positive cells. **(e, f)** Example single-marker positive cell images. **(g)** Example marker-negative cell images.

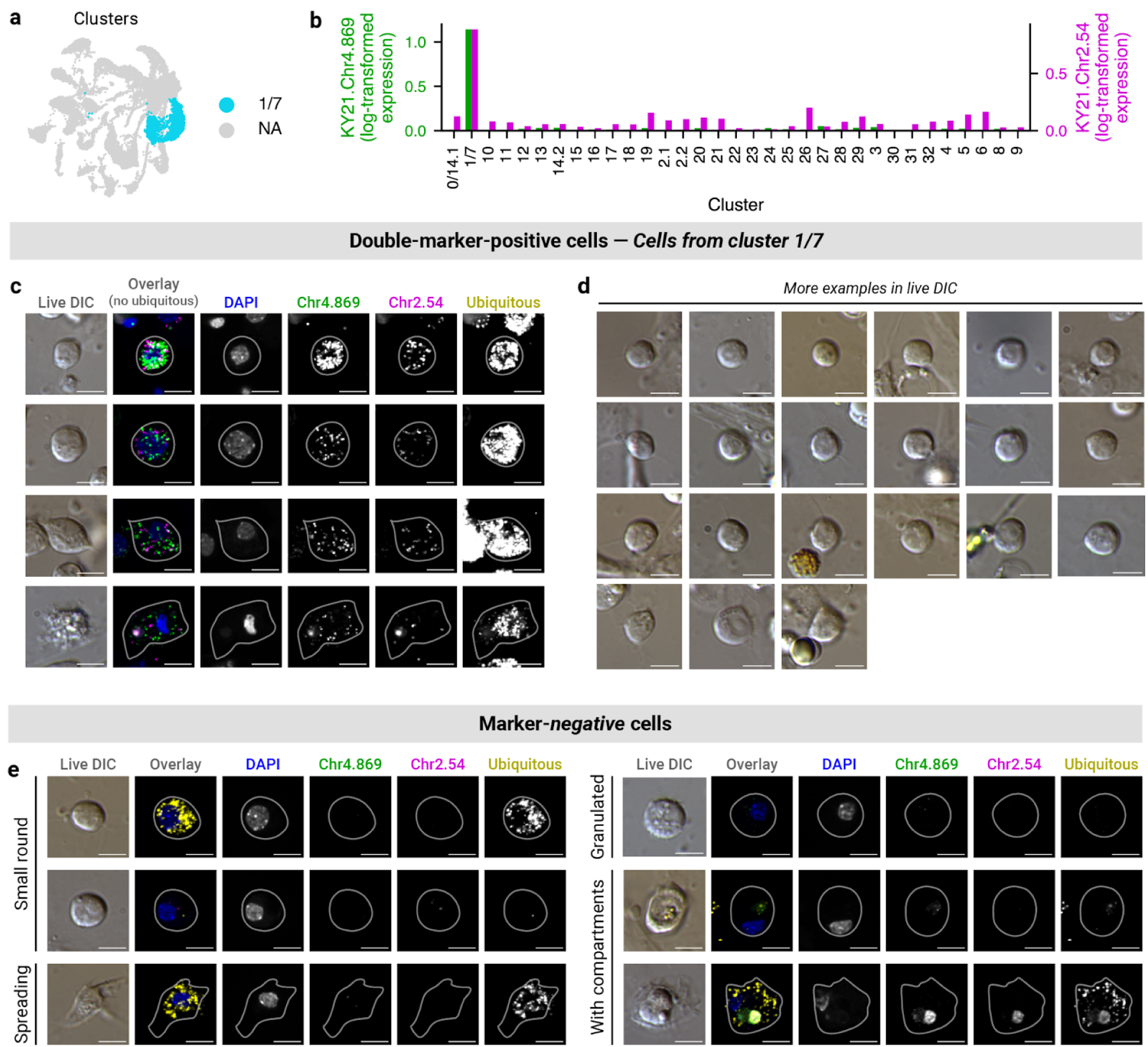
*3-2*: Cluster 1/7 (cLRP-1) **(a)** UMAP showing cluster 1/7. **(b)** Bar plot showing marker gene expression (log_10_(CP10k + 1)). **(c, d)** Example marker-positive cell images, shown **(c)** with or **(d)** without fluorescence. **(e)** Example marker-negative cell images.

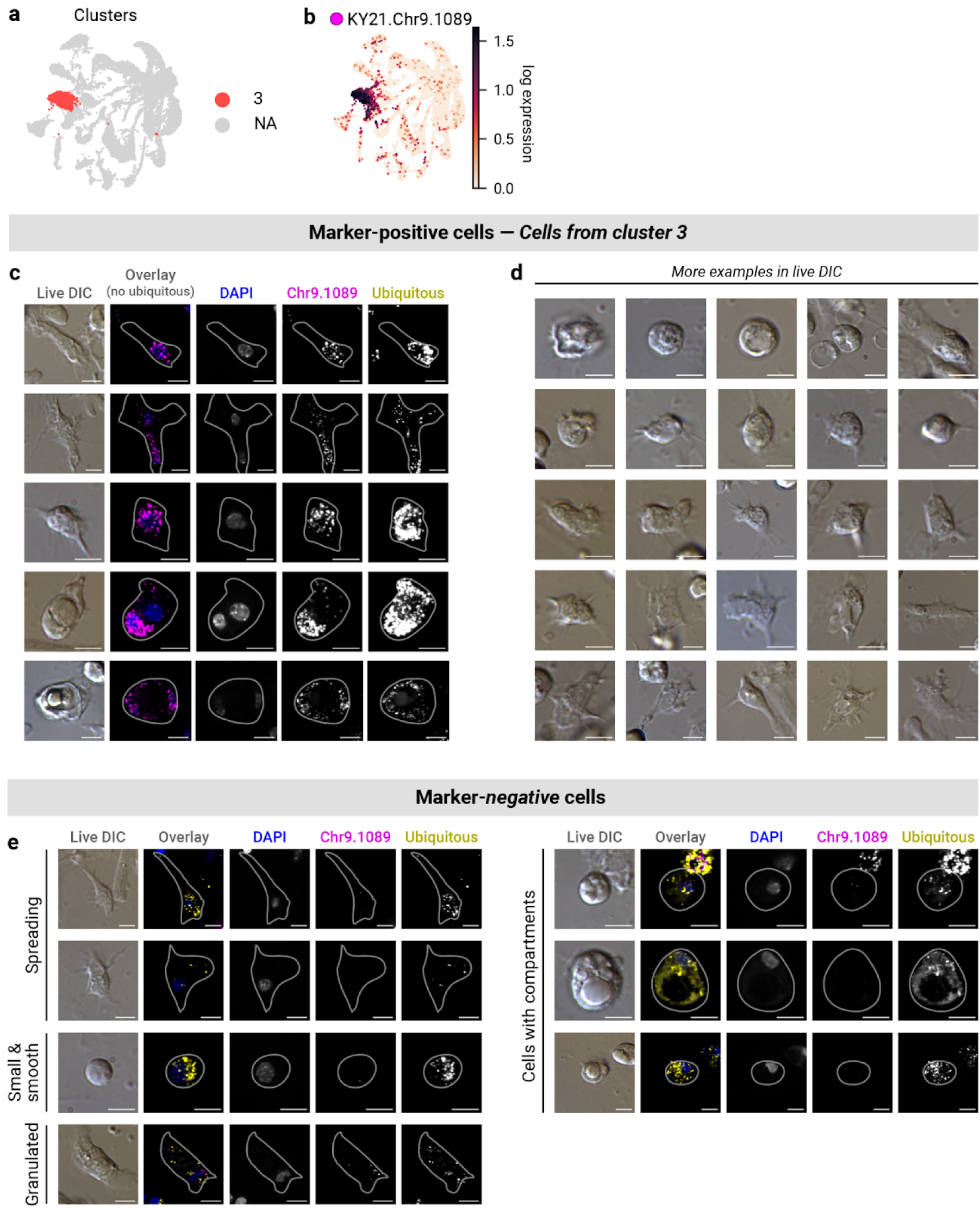
*3-3*: Cluster 3 (HA-1 (phag.)) **(a, b)** UMAPs showing **(a)** cluster 3 and **(b)** marker gene expression (log_10_(CP10k + 1)). **(c, d)** Example marker-positive cell images, shown **(c)** with or **(d)** without fluorescence. **(e)** Example marker-negative cell images.

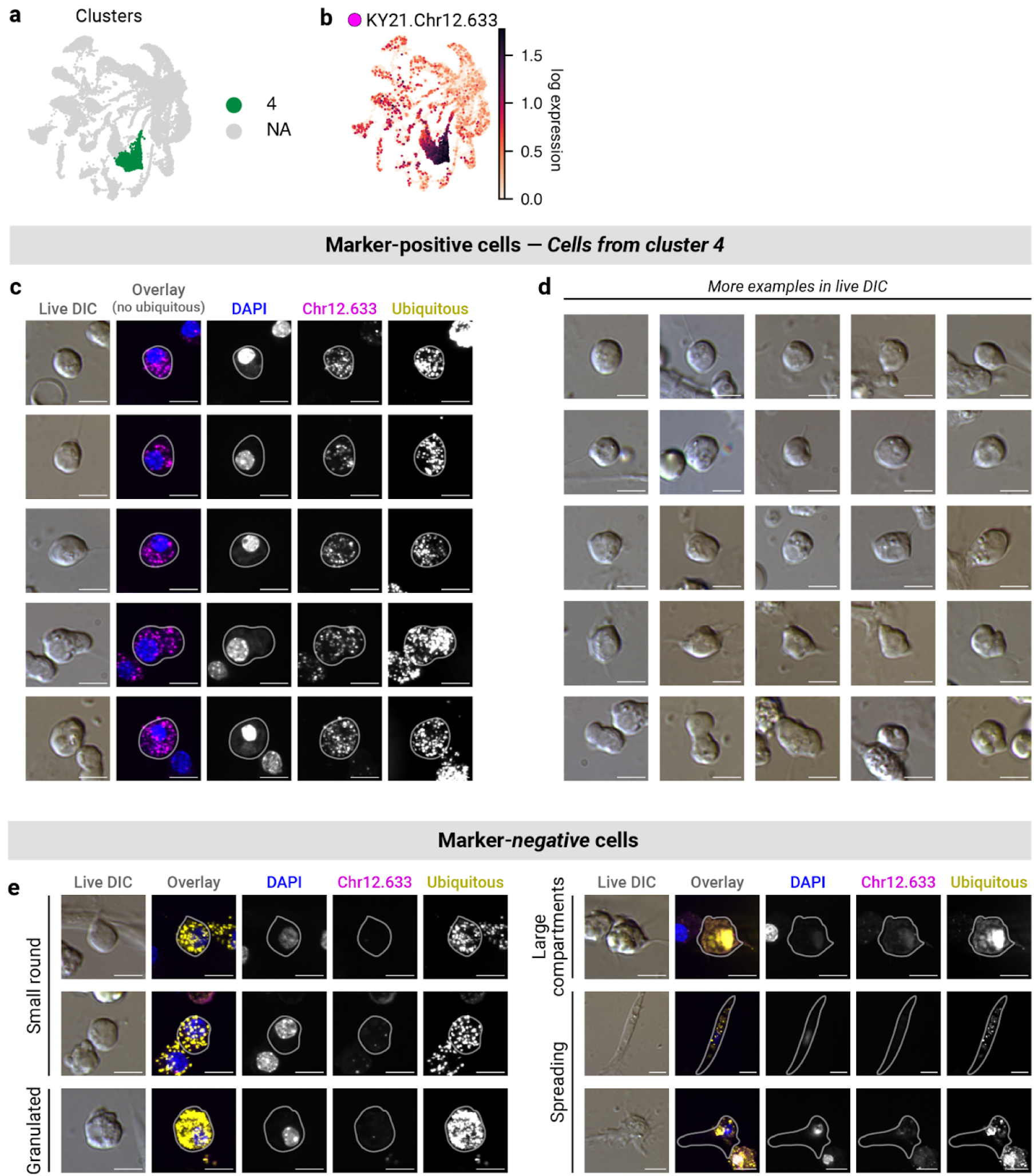
*3-4*: Cluster 4 (pre-RA) **(a, b)** UMAPs showing **(a)** cluster 4 and **(b)** marker gene expression (log_10_(CP10k + 1)). **(c, d)** Example marker-positive cell images, shown **(c)** with or **(d)** without fluorescence. **(e)** Example marker-negative cell images.

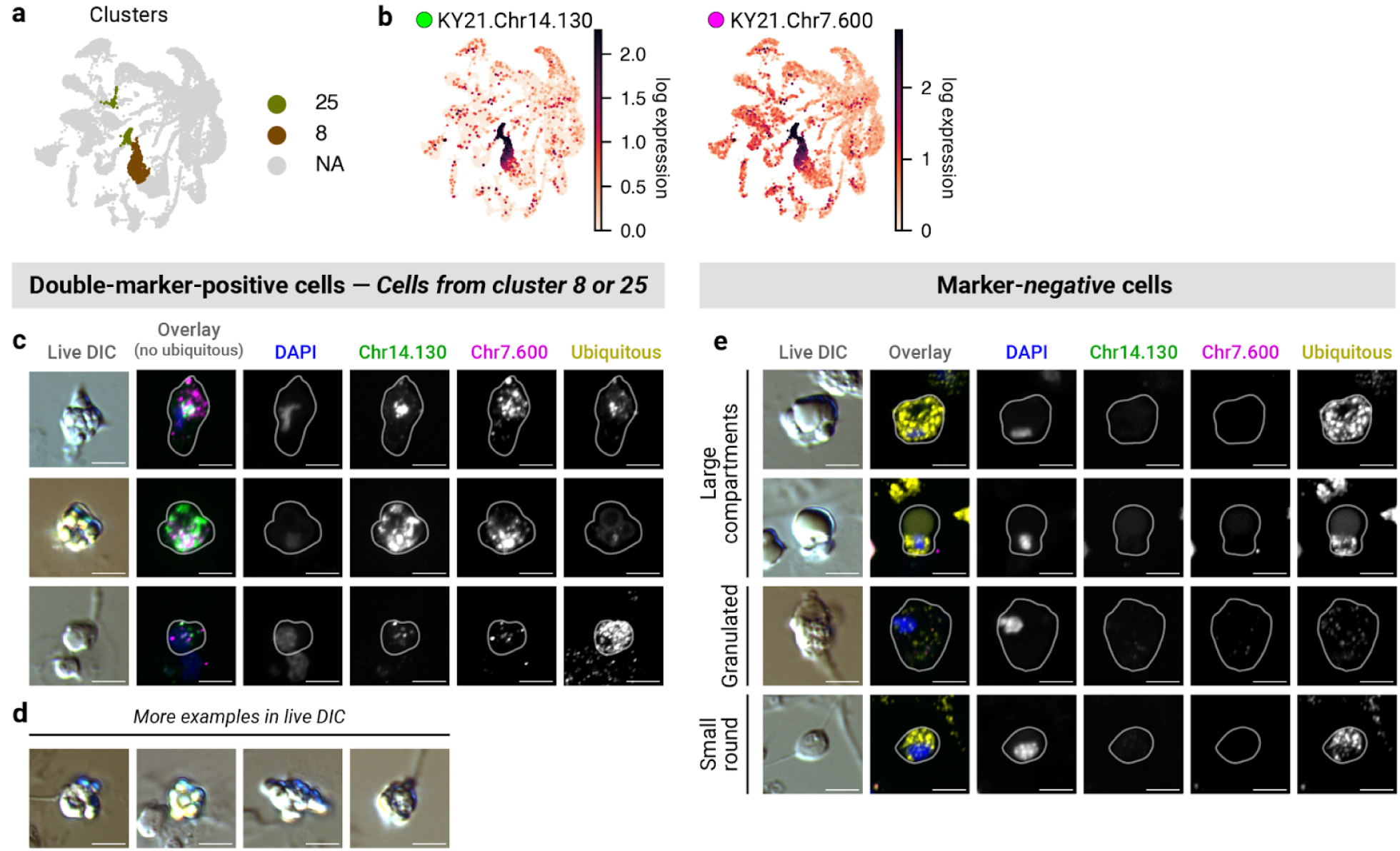
*3-5*: Cluster 8/25 (RA-1/2) **(a, b)** UMAPs showing **(a)** clusters 8 and 25 and **(b)** marker gene expression (log_10_(CP10k + 1)). **(c, d)** Example double-marker-positive cell images, shown **(c)** with or **(d)** without fluorescence. **(e)** Example marker-negative cell images.

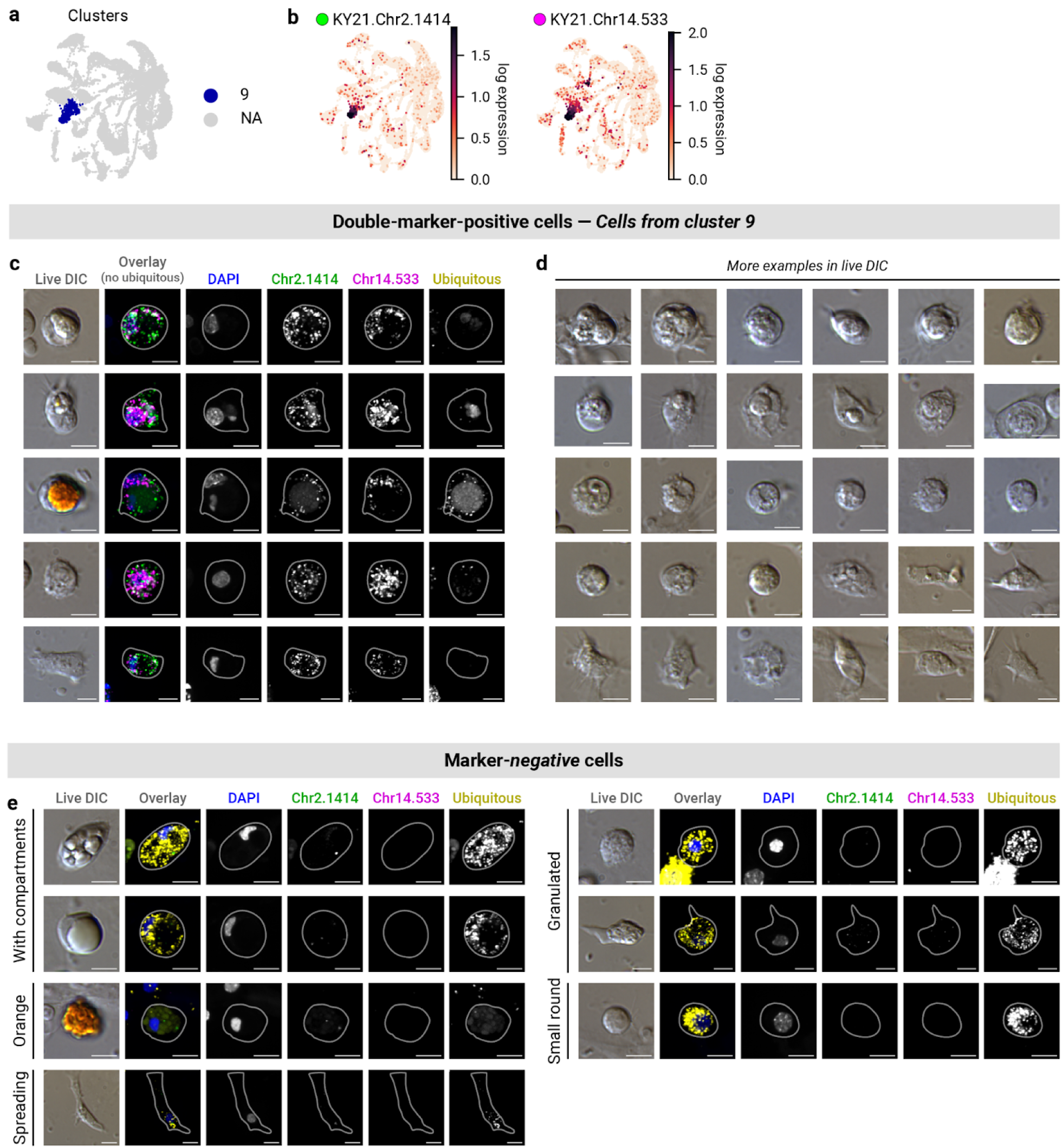
*3-6*: Cluster 9 (HA-2 (phag.)) **(a, b)** UMAPs showing **(a)** cluster 9 and **(b)** marker gene expression (log_10_(CP10k + 1)). **(c, d)** Example double-marker-positive cell images, shown **(c)** with or **(d)** without fluorescence. **(e)** Example marker-negative cell images.

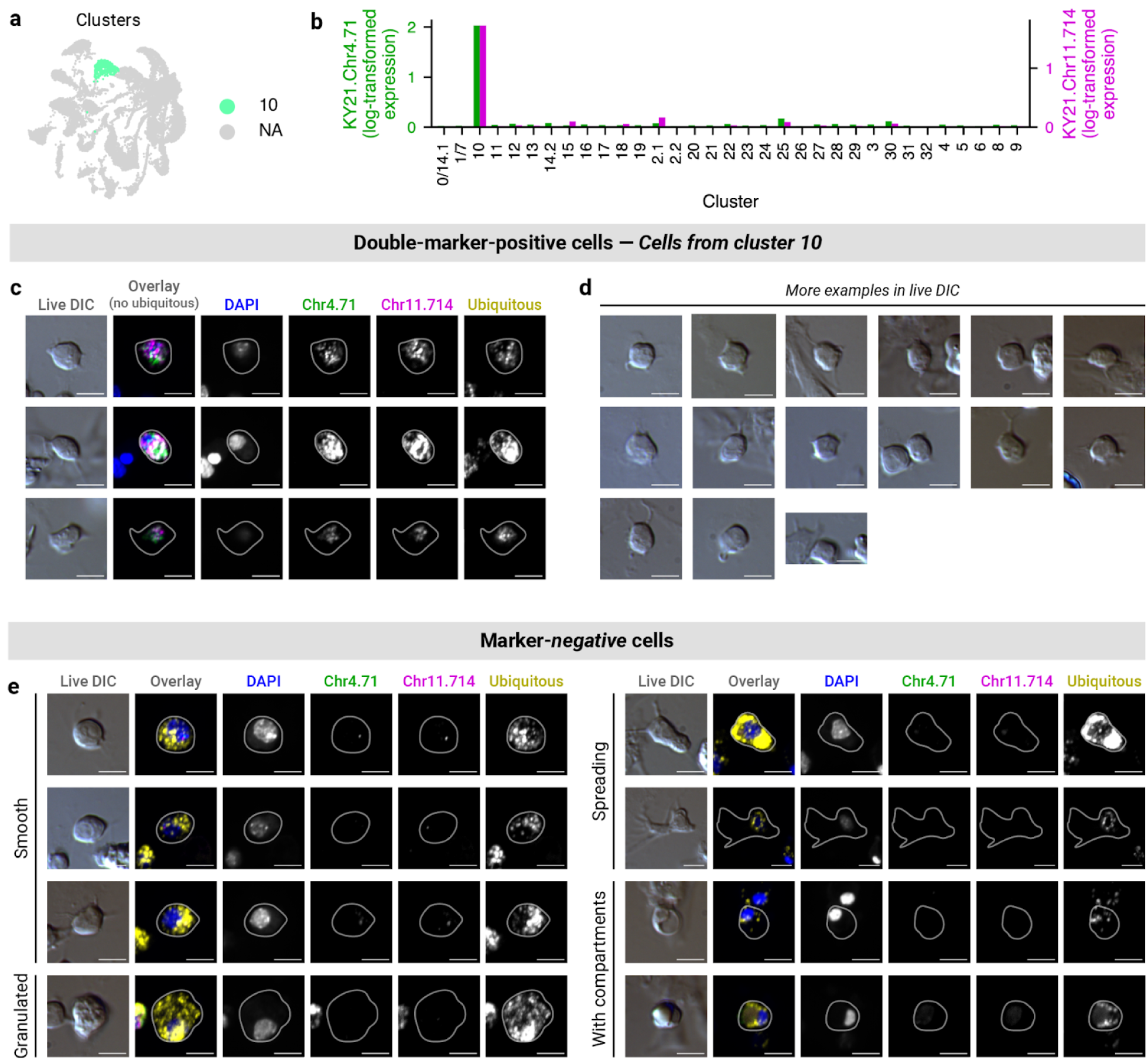
*3-7*: Cluster 10 (RSC) **(a)** UMAP showing cluster 10. **(b)** Bar plot showing marker gene expression (log_10_(CP10k + 1)). **(c, d)** Example double-marker-positive cell images, shown **(c)** with or **(d)** without fluorescence. **(e)** Example marker-negative cell images.

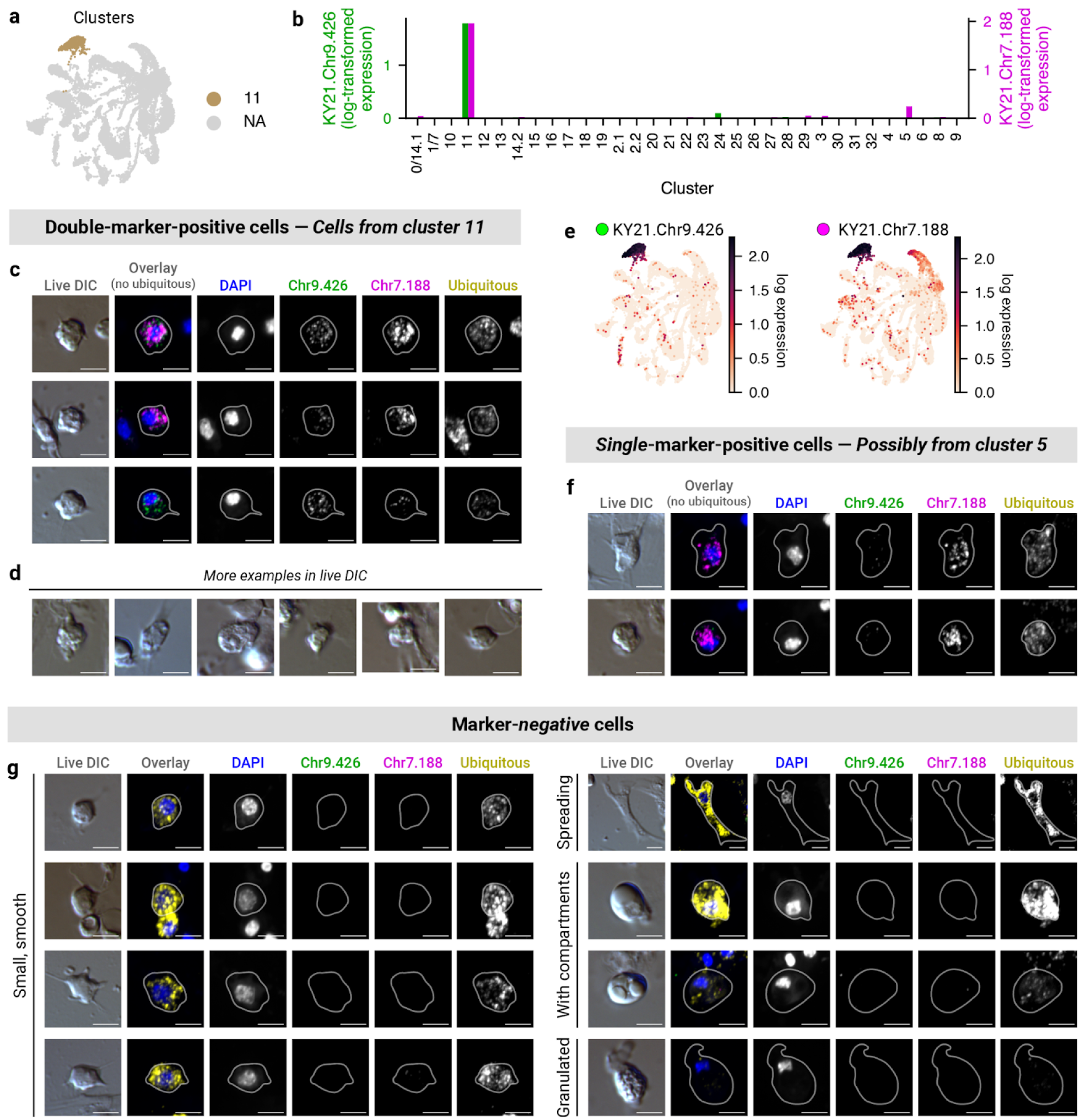
*3-8*: Cluster 11 (BLC) **(a)** UMAP showing cluster 11. **(b)** Bar plot showing marker gene expression (log_10_(CP10k + 1)). **(c, d)** Example double-marker-positive cell images, shown **(c)** with or **(d)** without fluorescence. **(e)** UMAPs showing marker gene expression (log_10_(CP10k + 1)) in part of cluster 5. **(f)** Example single-marker-positive cell images. **(g)** Example marker-negative cell images.

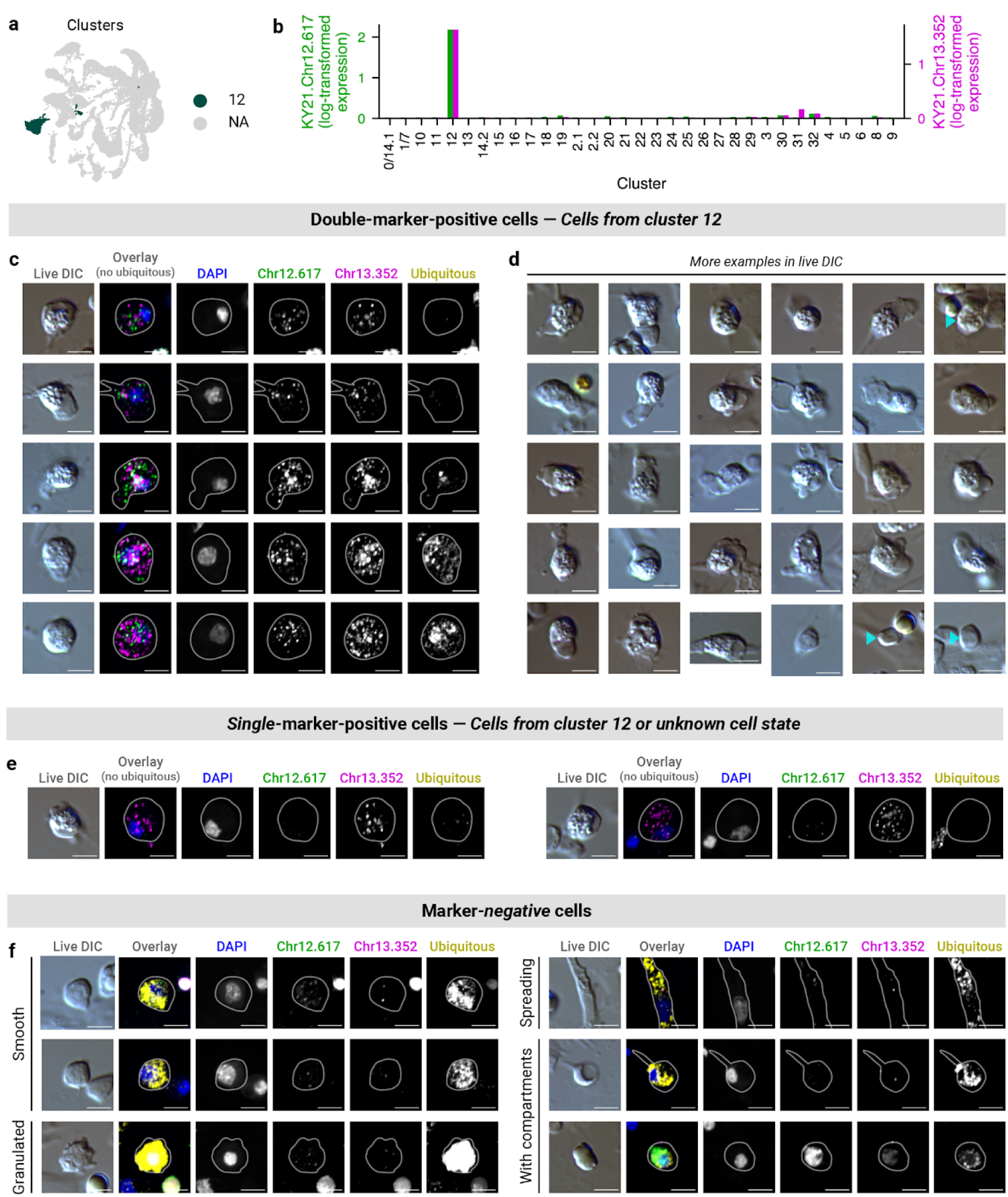
*3-9*: Cluster 12 (GA) **(a)** UMAP showing cluster 12. **(b)** Bar plot showing marker gene expression (log_10_(CP10k + 1)). **(c, d)** Example double-marker-positive cell images, shown **(c)** with or **(d)** without fluorescence. Teal arrows indicate double-marker-positive cells. **(e)** Example single-marker-positive cells. **(f)** Example marker-negative cell images.

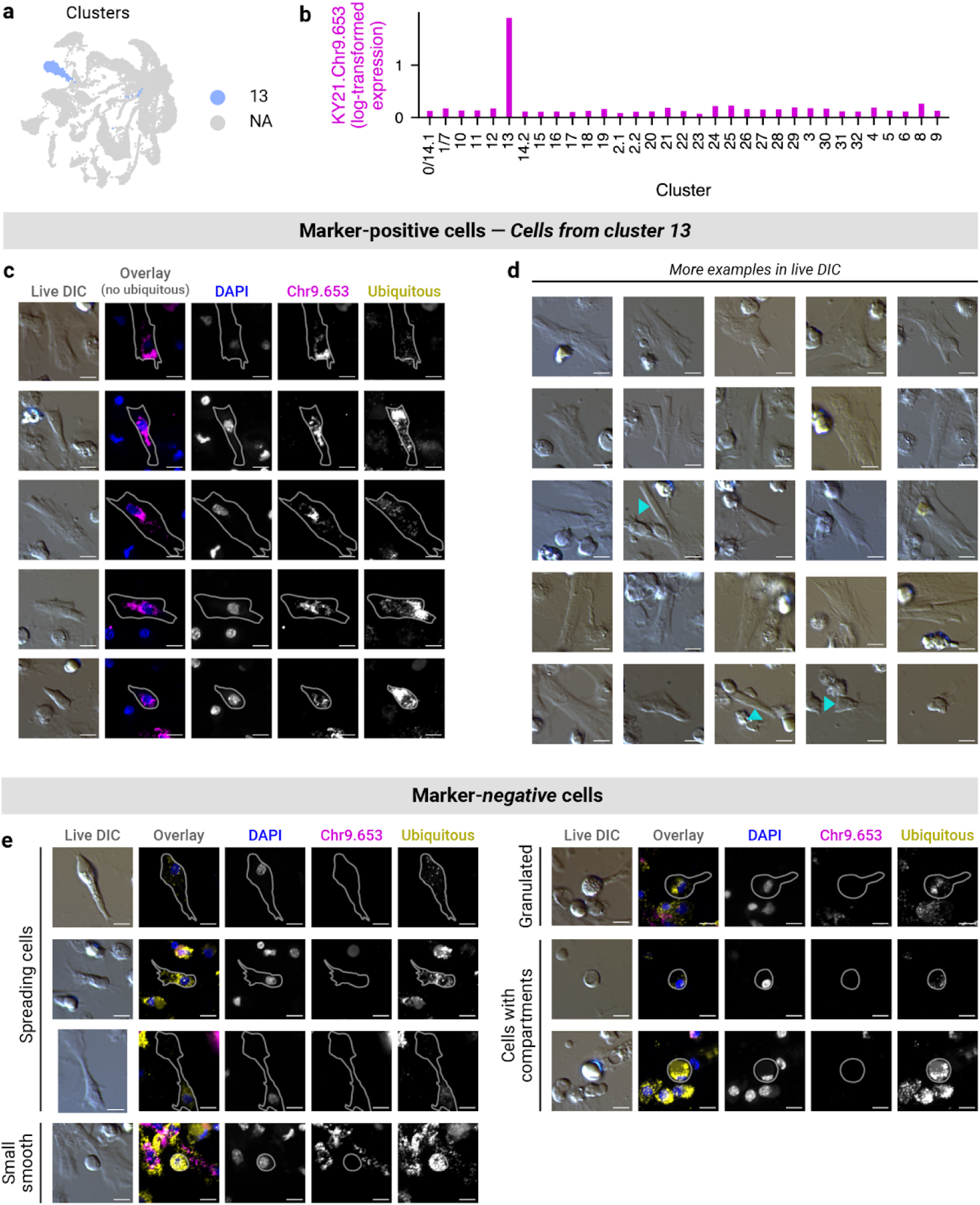
*3-10*: Cluster 13 (HA-3) **(a)** UMAP showing cluster 13. **(b)** Bar plot showing marker gene expression (log_10_(CP10k + 1)). **(c, d)** Example marker-positive cell images, shown **(c)** with or **(d)** without fluorescence. Teal arrows indicate marker-positive cells. **(e)** Example marker-negative cell images.

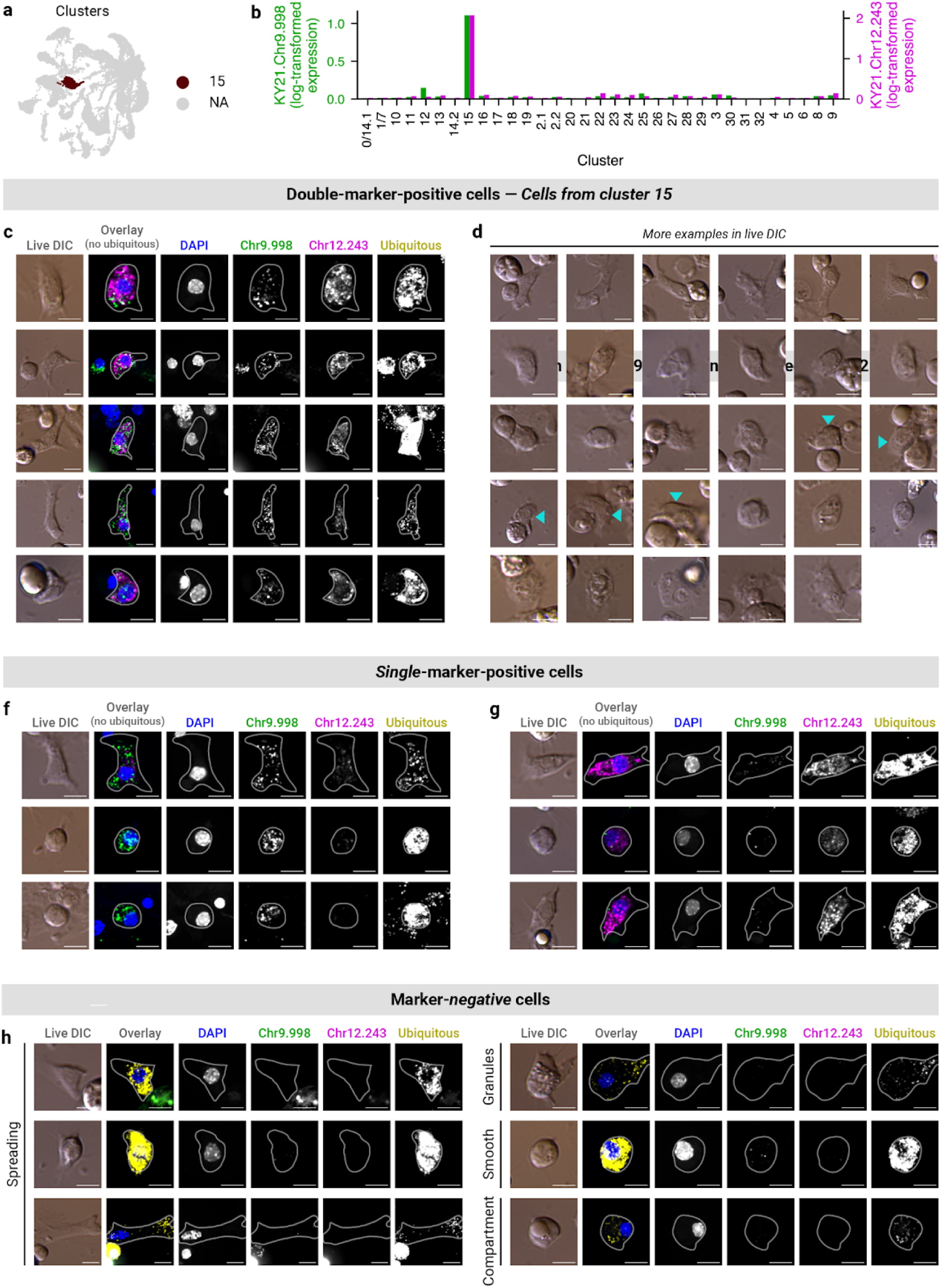
*3-11*: Cluster 13 (HA-3) **(a)** UMAP showing cluster 15. **(b)** Bar plot showing marker gene expression (log_10_(CP10k + 1)). **(c, d)** Example double-marker-positive cell images, shown **(c)** with or **(d)** without fluorescence. Teal arrows indicate double-marker-positive cells. **(e)** UMAPs showing marker gene expression (log_10_(CP10k + 1)). **(f, g)** Example single-marker-positive cell images. **(h)** Example marker-negative cell images.

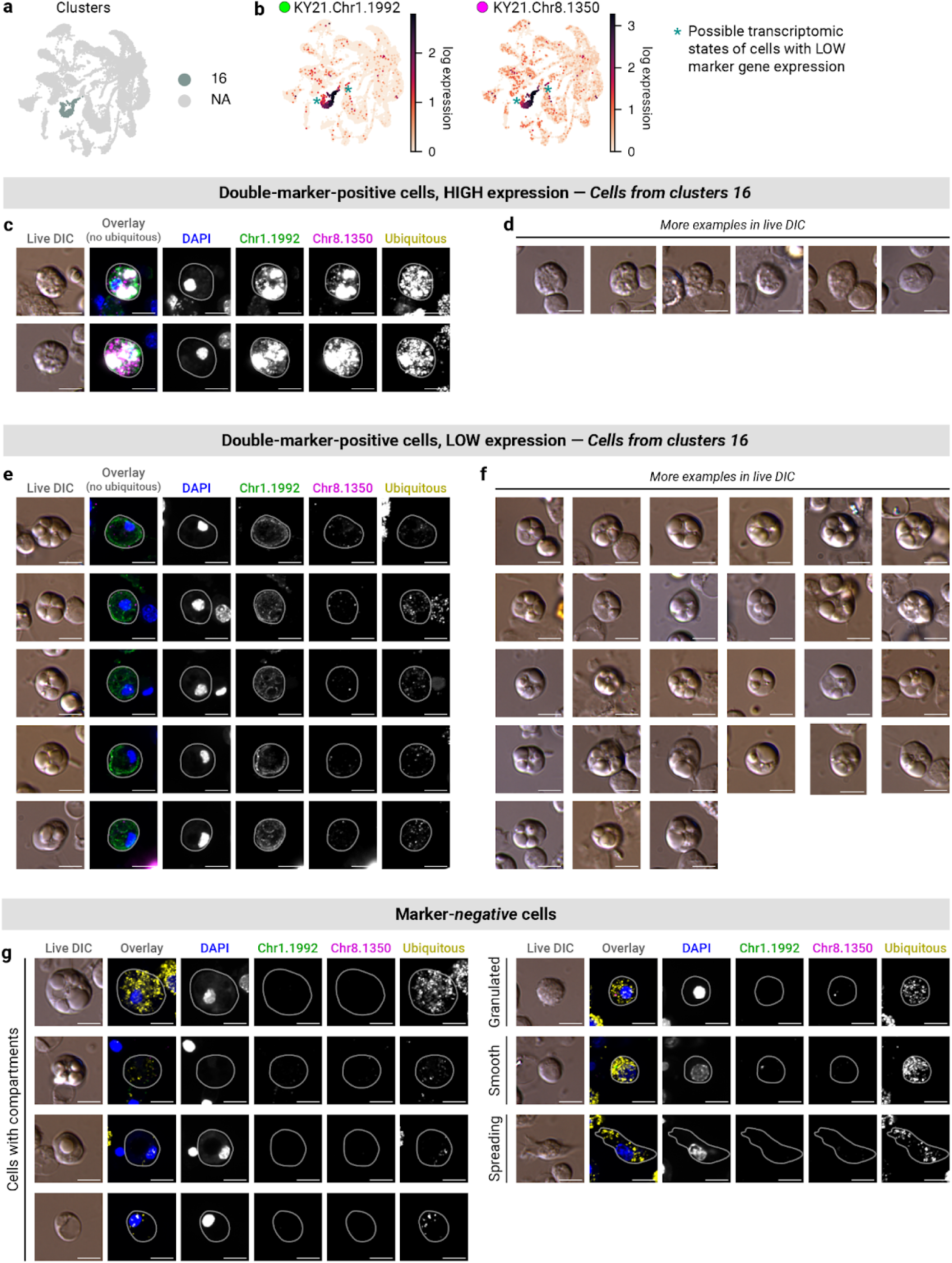
*3-12*: Cluster 13 (HA-3) **(a, b)** UMAPs showing **(a)** cluster 16 and **(b)** marker gene expression (log_10_(CP10k + 1)). **(c, d)** Example double-marker-positive cell images with high fluorescence, shown **(c)** with or **(d)** without fluorescence. **(e, f)** Example double-marker-positive cell images with low fluorescence, shown **(c)** with or **(d)** without fluorescence. **(g)** Example marker-negative cell images.

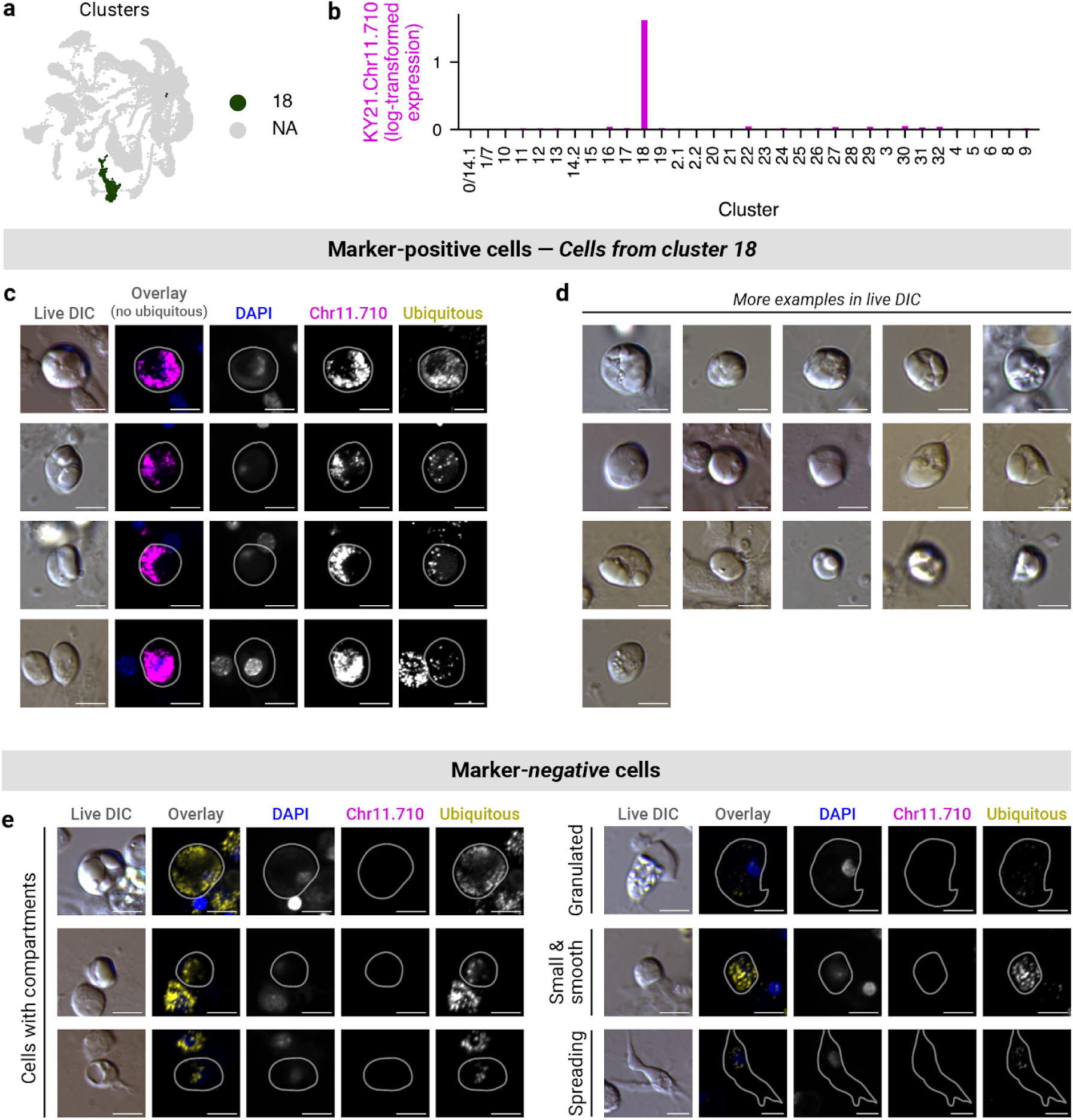
*3-13*: Cluster 18 (ICC) **(a)** UMAP showing cluster 18. **(b)** Bar plot showing marker gene expression (log_10_(CP10k + 1)). **(c, d)** Example marker-positive cell image, shown **(c)** with or **(d)** without fluorescence. **(e)** Example marker-negative cell images.

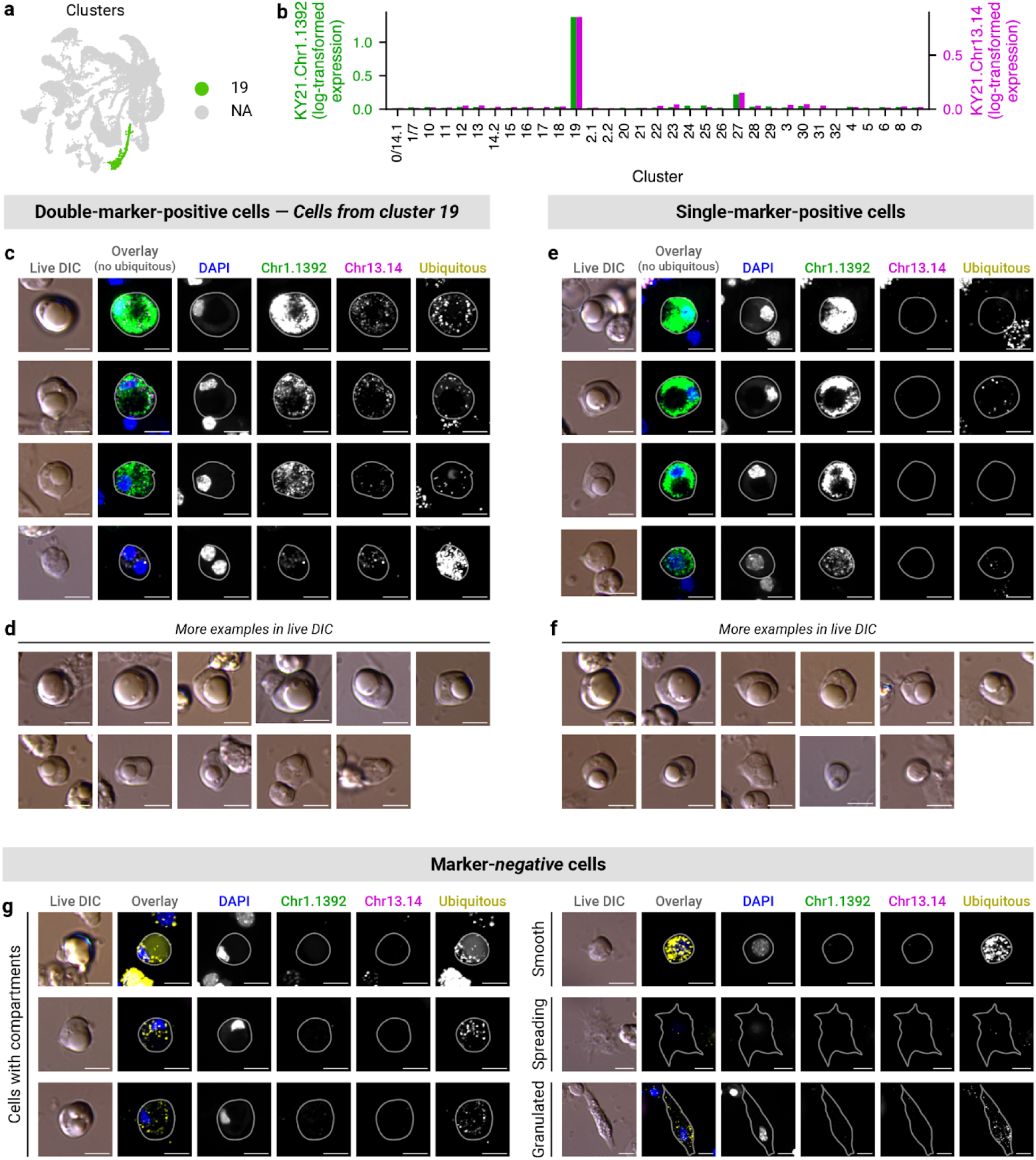
*3-14*: Cluster 19 (URG-1 (large)) **(a)** UMAP showing cluster 19. **(b)** Bar plot showing marker gene expression (log_10_(CP10k + 1)). **(c–f)** Example double-marker-positive cell images, shown **(c)** with or **(d)** without fluorescence. **(e, f)** Example single-marker-positive cell images, shown **(e)** with or **(f)** without fluorescence. **(g)** Example marker-negative cell images.

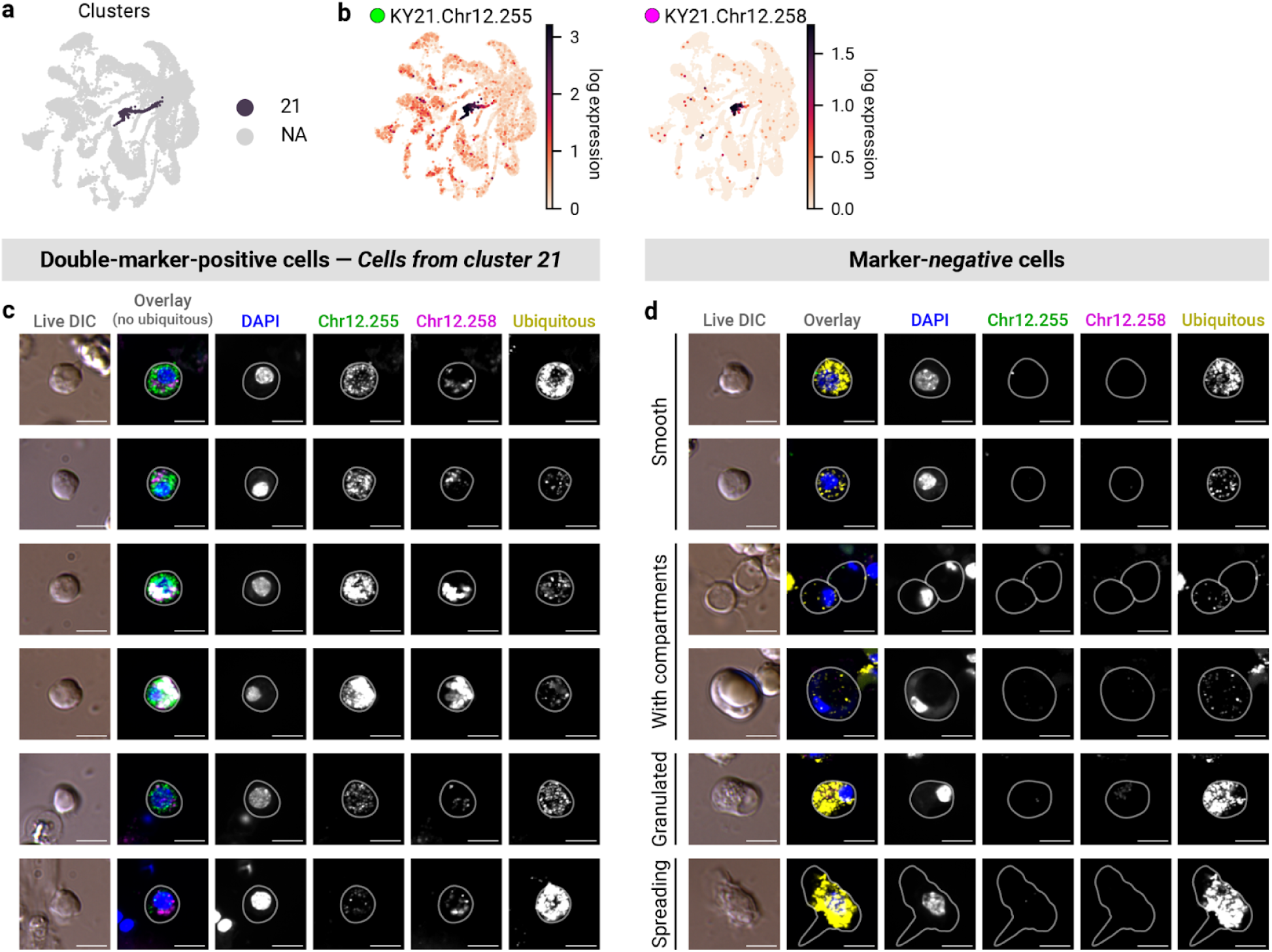
*3-15*: Cluster 21 (RC) **(a, b)** UMAPs showing **(a)** cluster 21 and **(b)** marker gene expression (log_10_(CP10k + 1)). **(c, d)** Example **(c)** marker-positive and **(d)** marker-negative cell images.

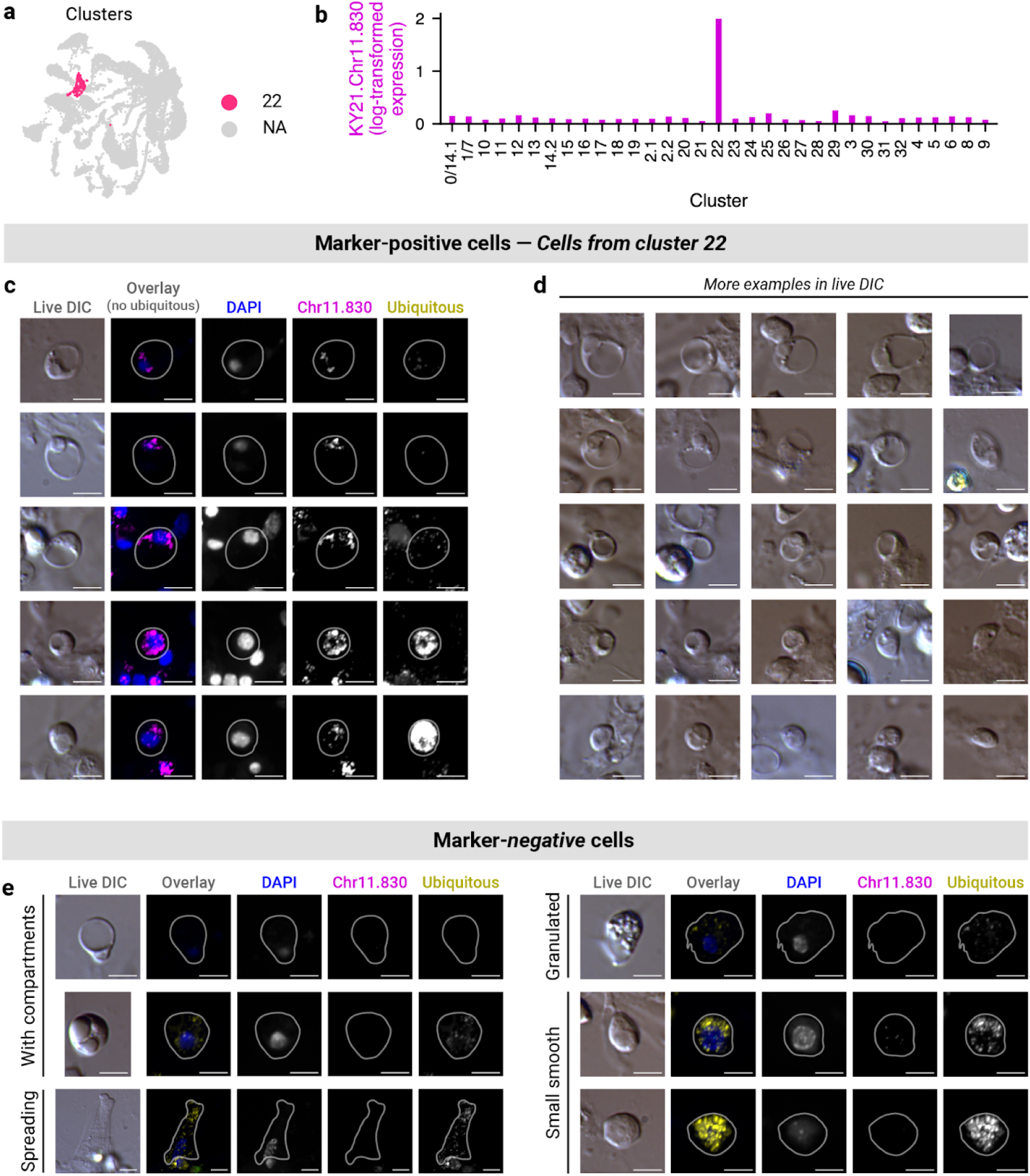
*3-16*: Cluster 22 (SRC) **(a)** UMAP showing cluster 22. **(b)** Bar plot showing marker gene expression (log_10_(CP10k + 1)). **(c, d)** Example marker-positive cell images, shown **(c)** with or **(d)** without fluorescence. **(e)** Example marker-negative cell images.

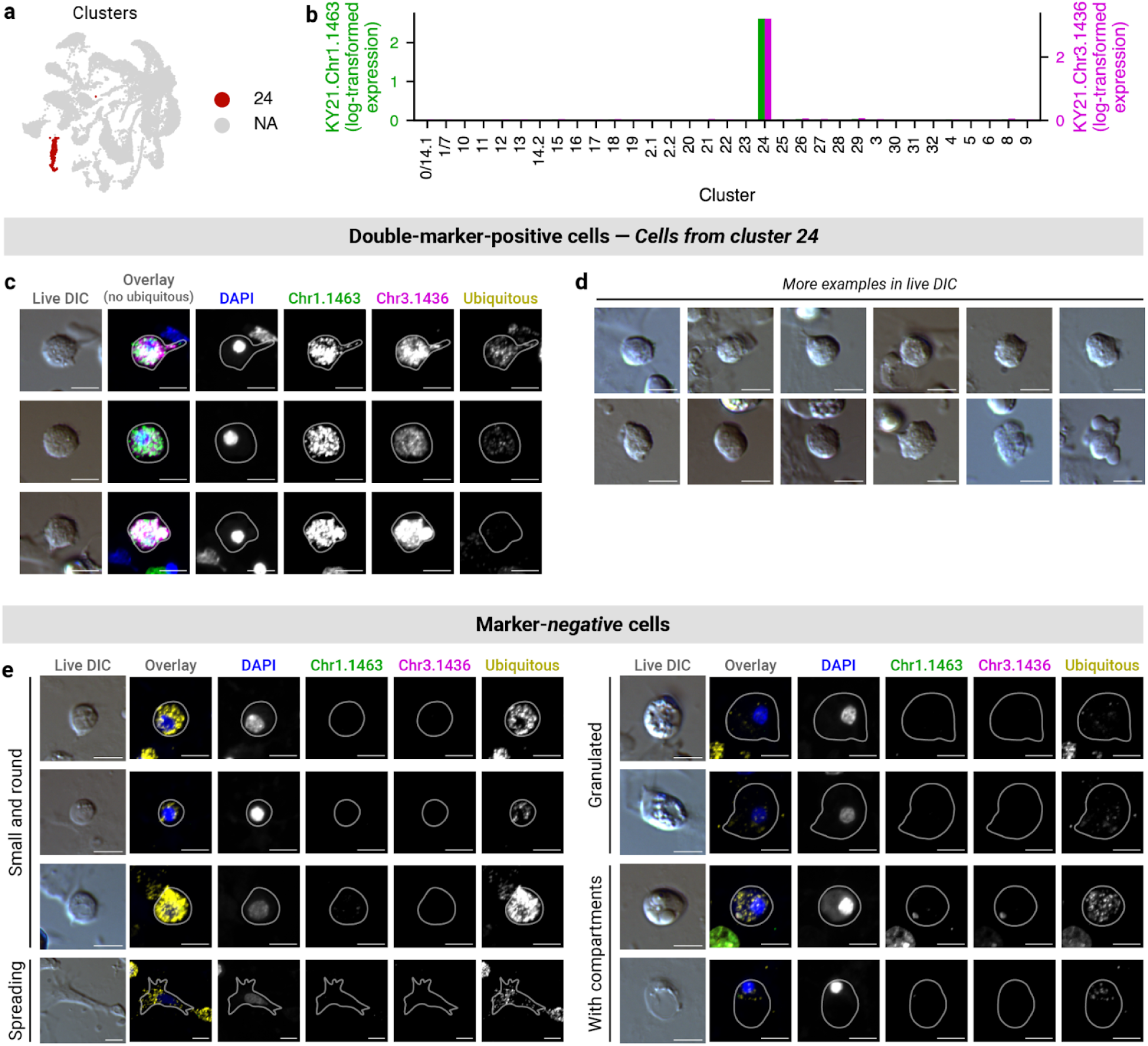
*3-17*: Cluster 24 (SGH-1) **(a)** UMAP showing cluster 24. **(b)** Bar plot showing marker gene expression (log_10_(CP10k + 1)). **(c, d)** Example double-marker-positive cell images, shown **(c)** with or **(d)** without fluorescence. **(e)** Example marker-negative cell images.

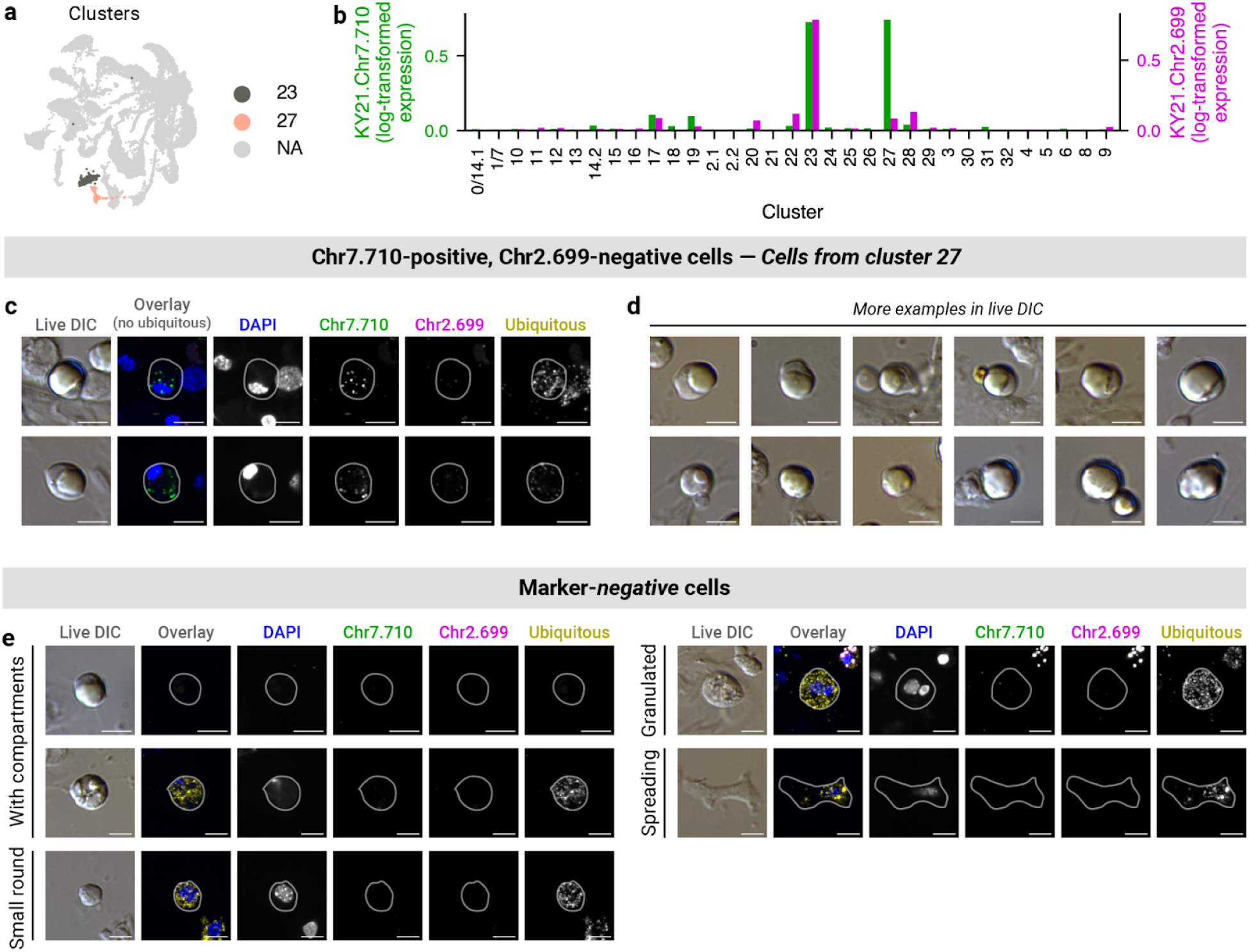
*3-18*: Cluster 27 (URG-2 (small)) **(a)** UMAP showing clusters 27 and 23. **(b)** Bar plot showing marker gene expression (log_10_(CP10k + 1)). Double-marker-positive cells (not observed) would represent cluster 23. Chr7.710-positive and Chr2.699-negative cells represent cluster 27. **(c, d)** Example Chr7.710-positive and Chr2.699-negative cell images, shown **(c)** with or **(d)** without fluorescence. **(e)** Example marker-negative cell images.

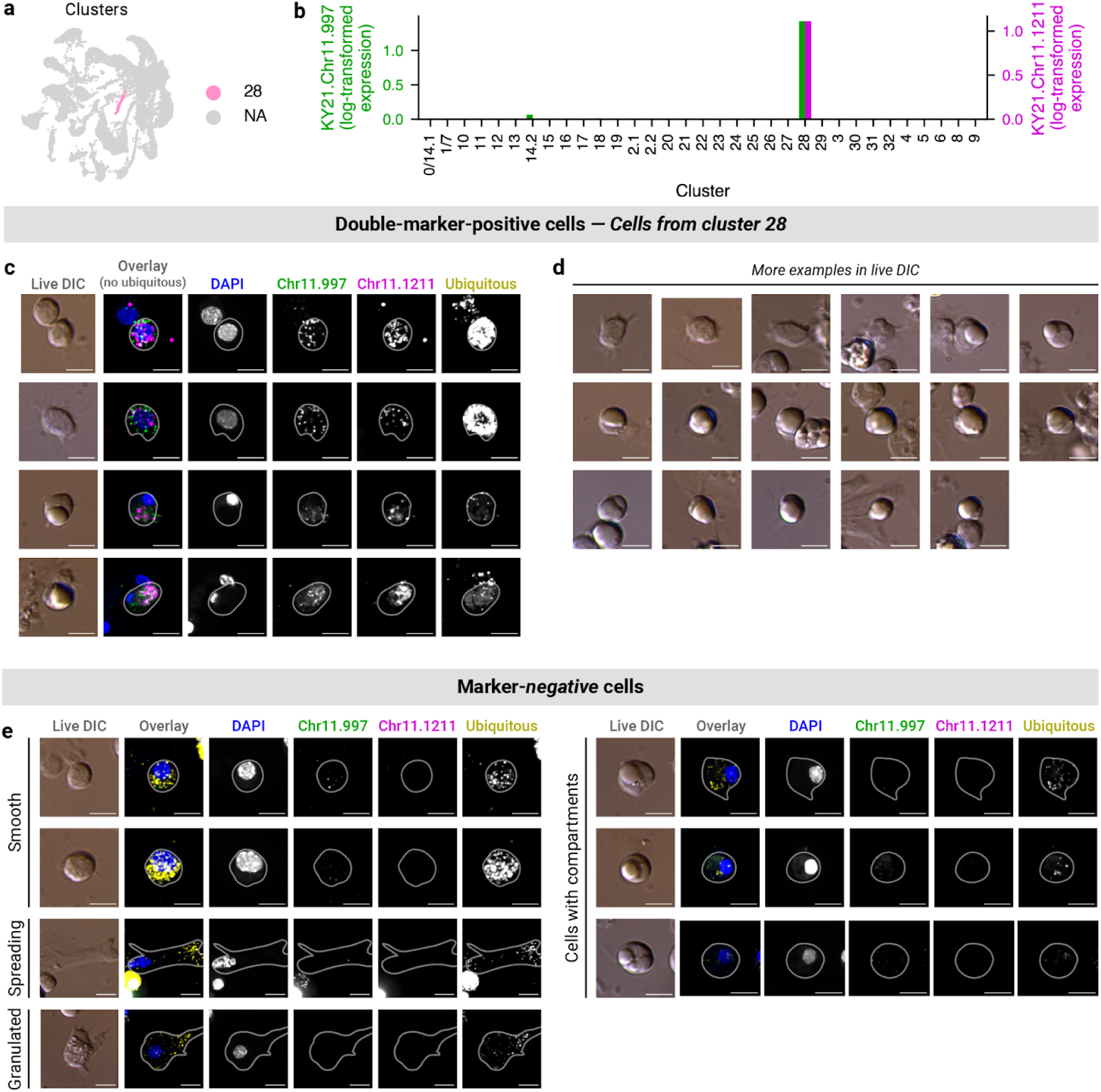
*3-19*: Cluster 28 (URG-3 (small)) **(a)** UMAP showing cluster 28 **(b)** Bar plot showing marker gene expression (log_10_(CP10k + 1)). **(c, d)** Example double-marker-positive cell images, shown **(c)** with or **(d)** without fluorescence. **(e)** Example marker-negative cell images.

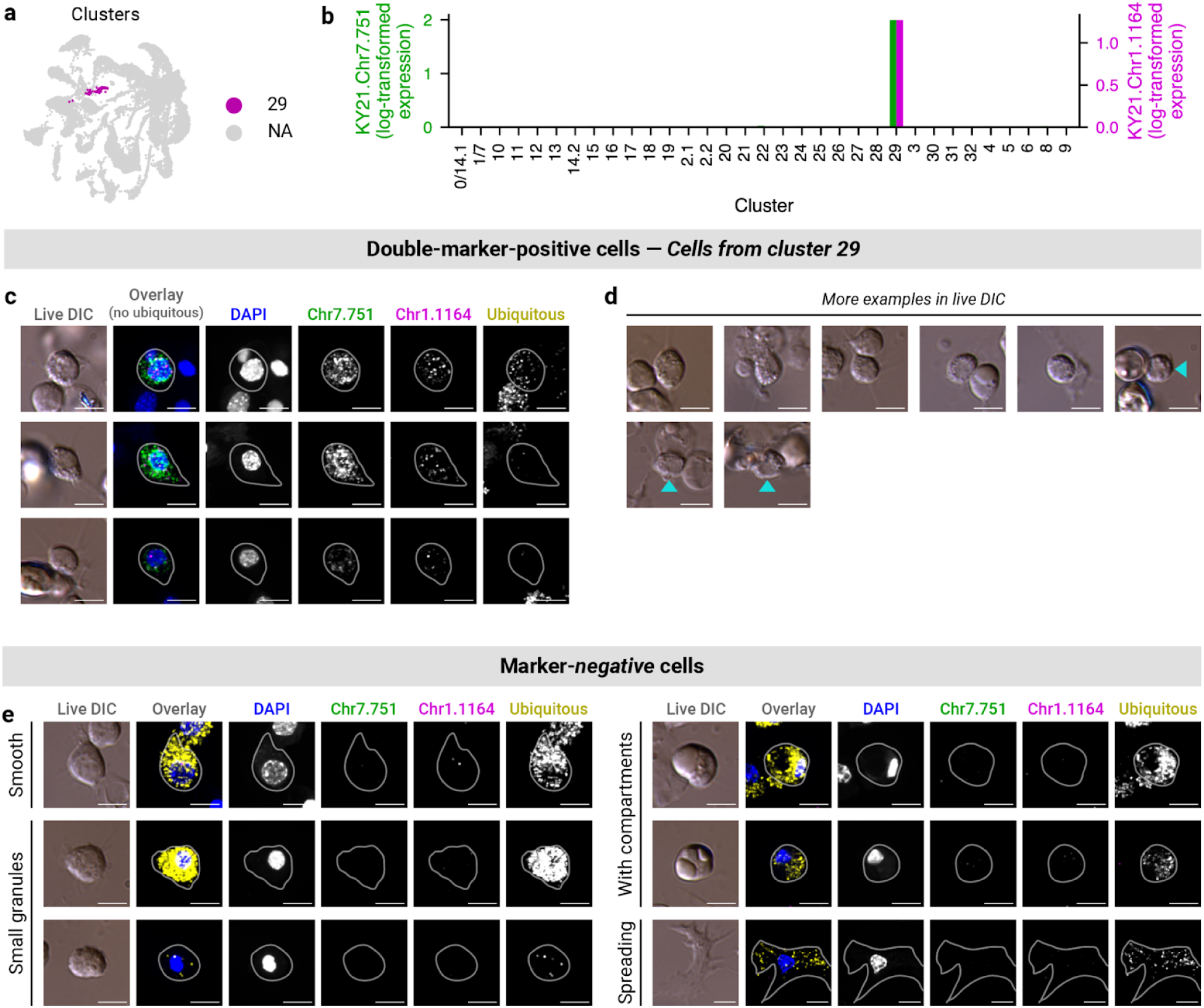
*3-20*: Cluster 29 (SGH-2) **(a)** UMAP showing cluster 29. **(b)** Bar plot showing marker gene expression (log_10_(CP10k + 1)). **(c, d)** Example double-marker-positive cell images, shown **(c)** with or **(d)** without fluorescence. Teal arrows indicate double-marker-positive cells. **(e)** Example marker-negative cell images.

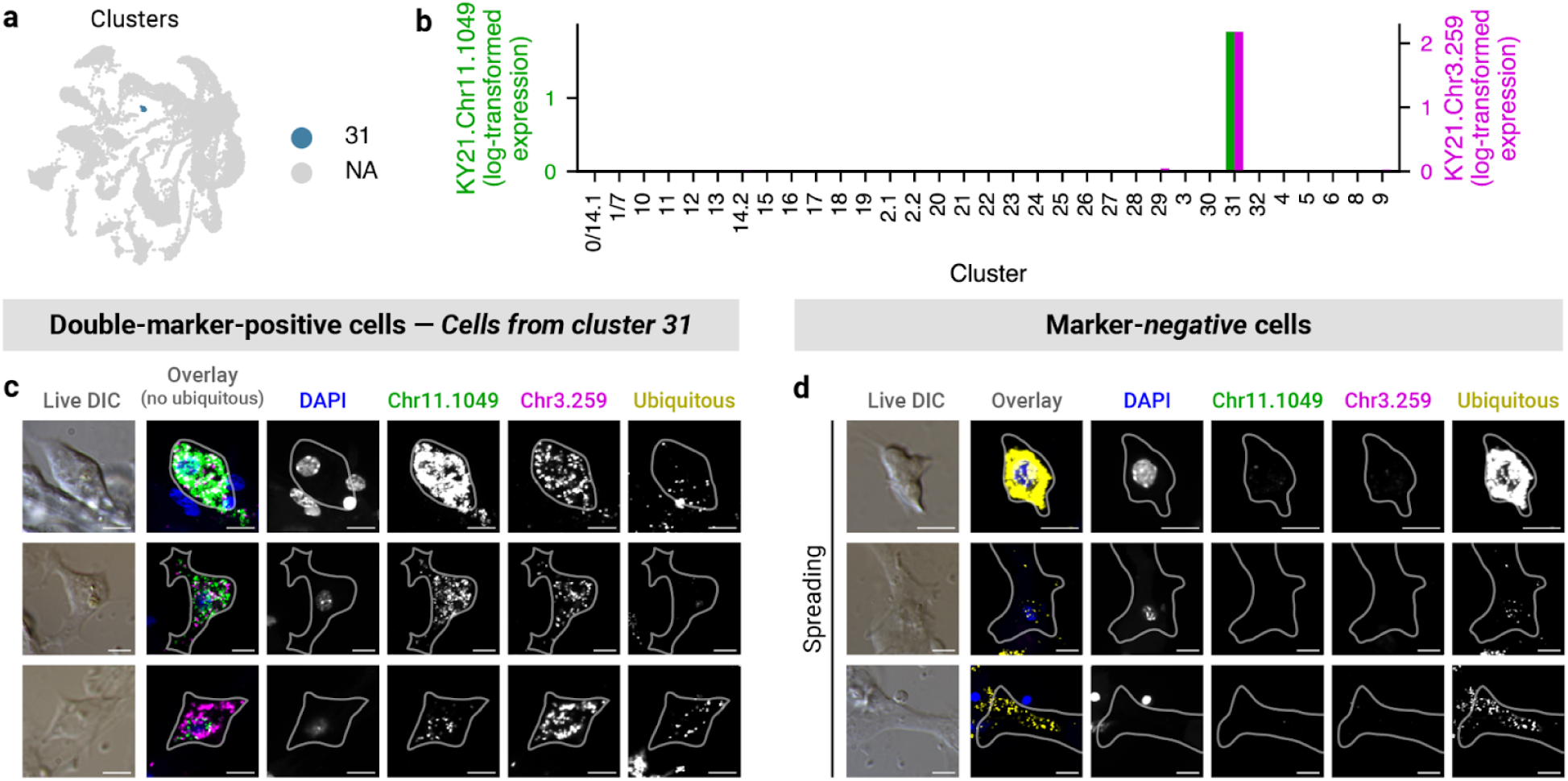
*3-21*: Cluster 31 (HA-5) **(a)** UMAP showing cluster 31. **(b)** Bar plot showing marker gene expression (log_10_(CP10k + 1)). **(c, d)** Example **(c)** marker-positive and **(d)** marker-negative cell images. All scale bars in 3-1 through 3-21 are 5 µm.

**Supplementary Fig. 4:**
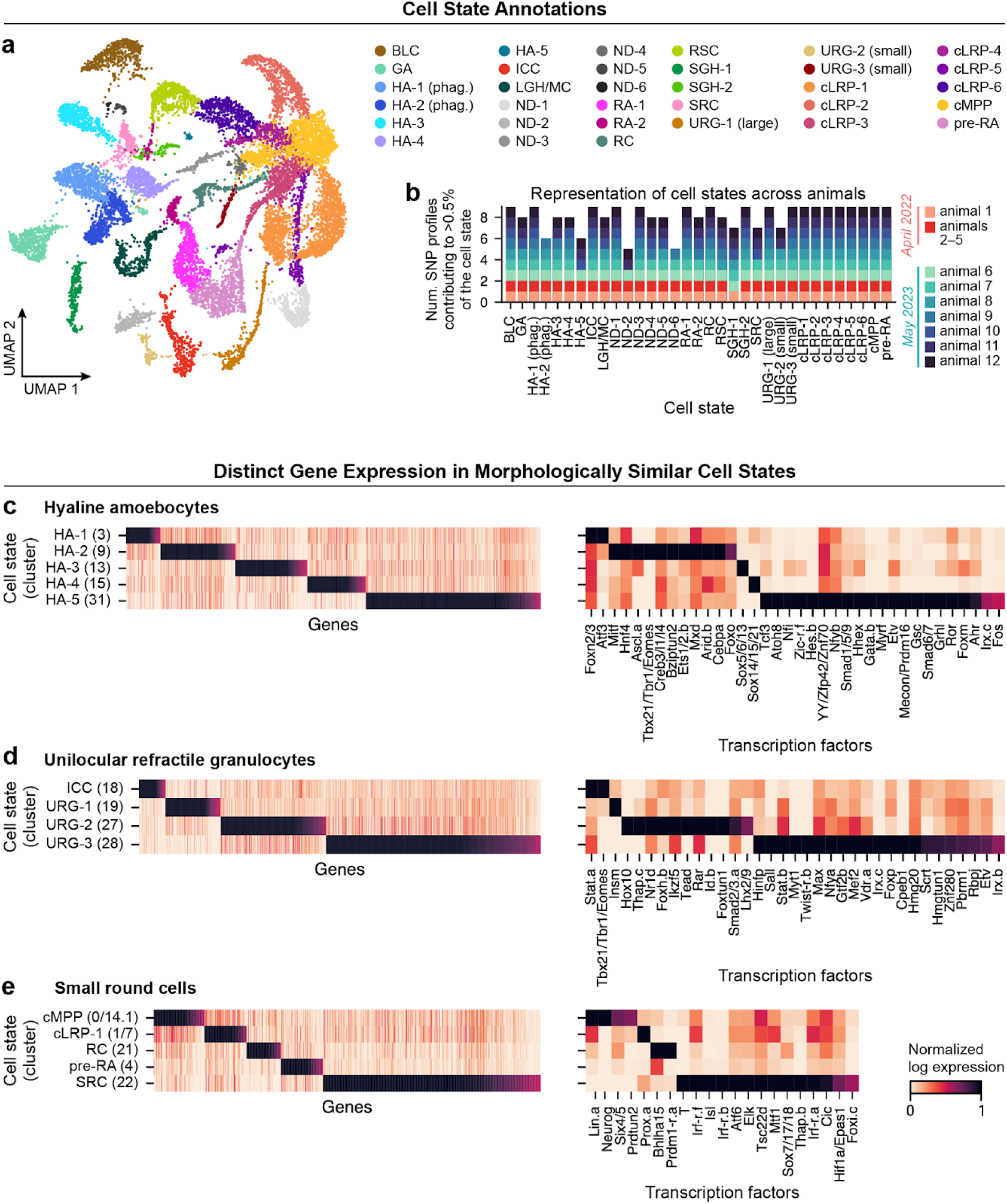
Cell state annotations based on HCR FISH, showing morphologically similar cells are transcriptionally distinct **(a)** UMAP colored by cell state annotation. **(b)** Number of SNP profiles contributing to each cell state. **(c–e)** Heatmaps showing genes which distinguish cell clusters that share the **(a)** hyaline amoebocyte, **(b)** unilocular refractile granulocyte, and **(c)** small round morphologies. Heatmaps on the left show all genes for which the mean log expression in the cluster of focus is more than two times higher than the level in all other clusters of the similar morphology. Heatmaps on the right show the subset of those genes which are transcription factors. Abbreviations: BLC, blebbing-like cell; GA, granular amoebocyte; HA, hyaline amoebocyte; ICC, irregular compartment cell; LGH/MC, large granules hemocyte/morula cell; RA, refractile amoebocyte; RC, round cell; RSC, round spreading cell; SGH, small granules hemocyte; SRC, signet ring cell; URG, unilocular refractile granulocyte; cLRP, candidate lineage-restricted progenitor; cMPP, candidate multipotent progenitor; ND, not determined.

**Supplementary Fig. 5:**
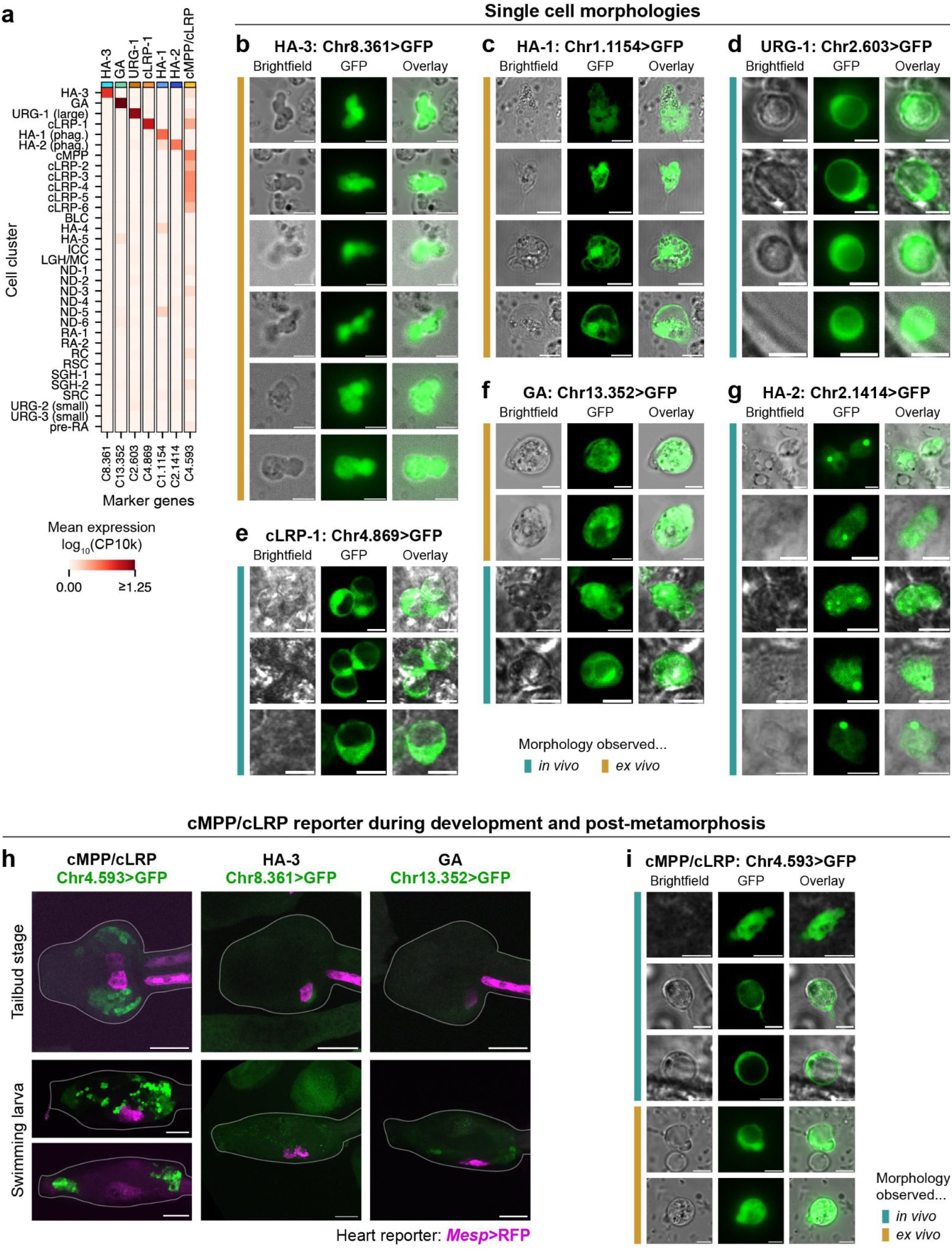
Live reporters **(a)** Heatmap of marker genes chosen for designing live reporters. **(b–g)** Example images showing single-cell morphologies for reporters predicted to label **(b)** HA-3, **(c)** HA-1, **(d)** URG-1, **(e)** cLRP-1, **(f)** GA, and **(g)** HA-2. Colored lines indicate which images were taken *in vivo* vs. *ex vivo*. Scale bars = 5 µm. **(h)** Images showing blood cell reporter (green) and heart reporter (magenta) fluorescence at the tailbud stage and the swimming larva stage. Only the progenitor reporter is fluorescent at these stages. The larva images for the cMPP/cLRP and GA reporters have been reflected to make the larva’s anterior to the left and the ventral side towards the bottom of each image. Scale bars = 50 µm. **(i)** Example images showing heterogeneous single-cell morphologies for the cMPP/cLRP reporter. Scale bars = 5 µm.

**Supplementary Fig. 6:**
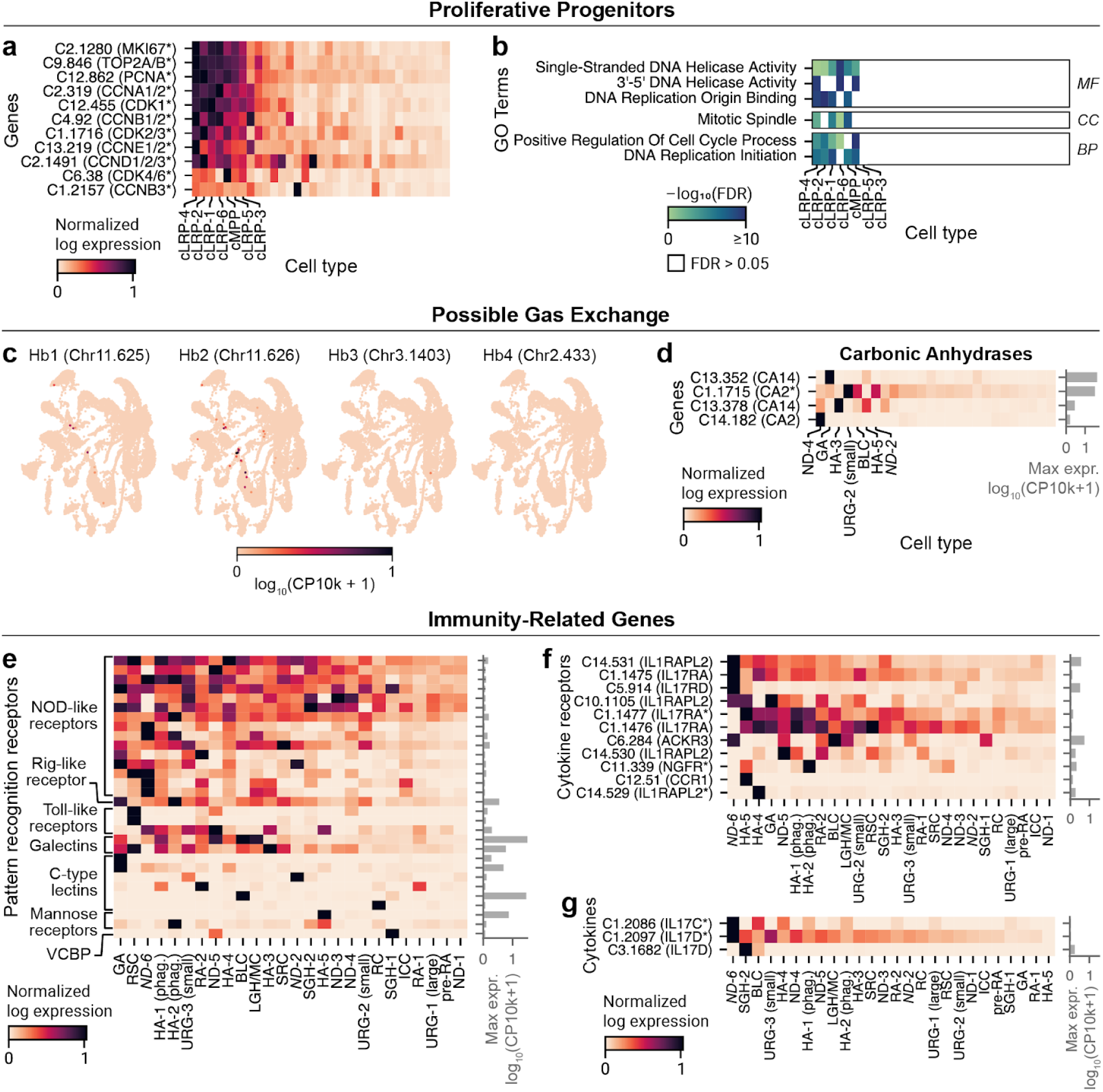
Gene expression across *C. robusta* cell states **(a, b)** Heatmaps showing **(a)** marker gene expression and **(b)** GO term enrichment related to proliferation. GO terms are separated by Molecular Function (MF), Cellular Component (CC) and Biological Process (BP). See **Supplementary Table 18** for accession numbers. **(c)** UMAP showing low or no expression of globin genes. **(d–g)** Heatmap showing expression of genes likely to be **(d)** carbonic anhydrases, **(e)** PRRs, **(f)** cytokine receptors, and **(g)** cytokines. Bar graphs to the right of each heatmap show each gene’s maximum mean expression across clusters. Genes in heatmaps are listed with an abbreviated *C. robusta* gene label (e.g. C2.319 instead of KY21.Chr2.319), followed by the closest human BLAST hit in parentheses. Asterisks indicate that the listed human gene(s) are OrthoFinder homologs. In this and all subsequent supplementary figures, cell states which both are present in fewer than 7/9 SNP profiles (see Supplementary Fig. 4b) and have not been verified by HCR FISH are labeled with italic text.

**Supplementary Fig. 7:**
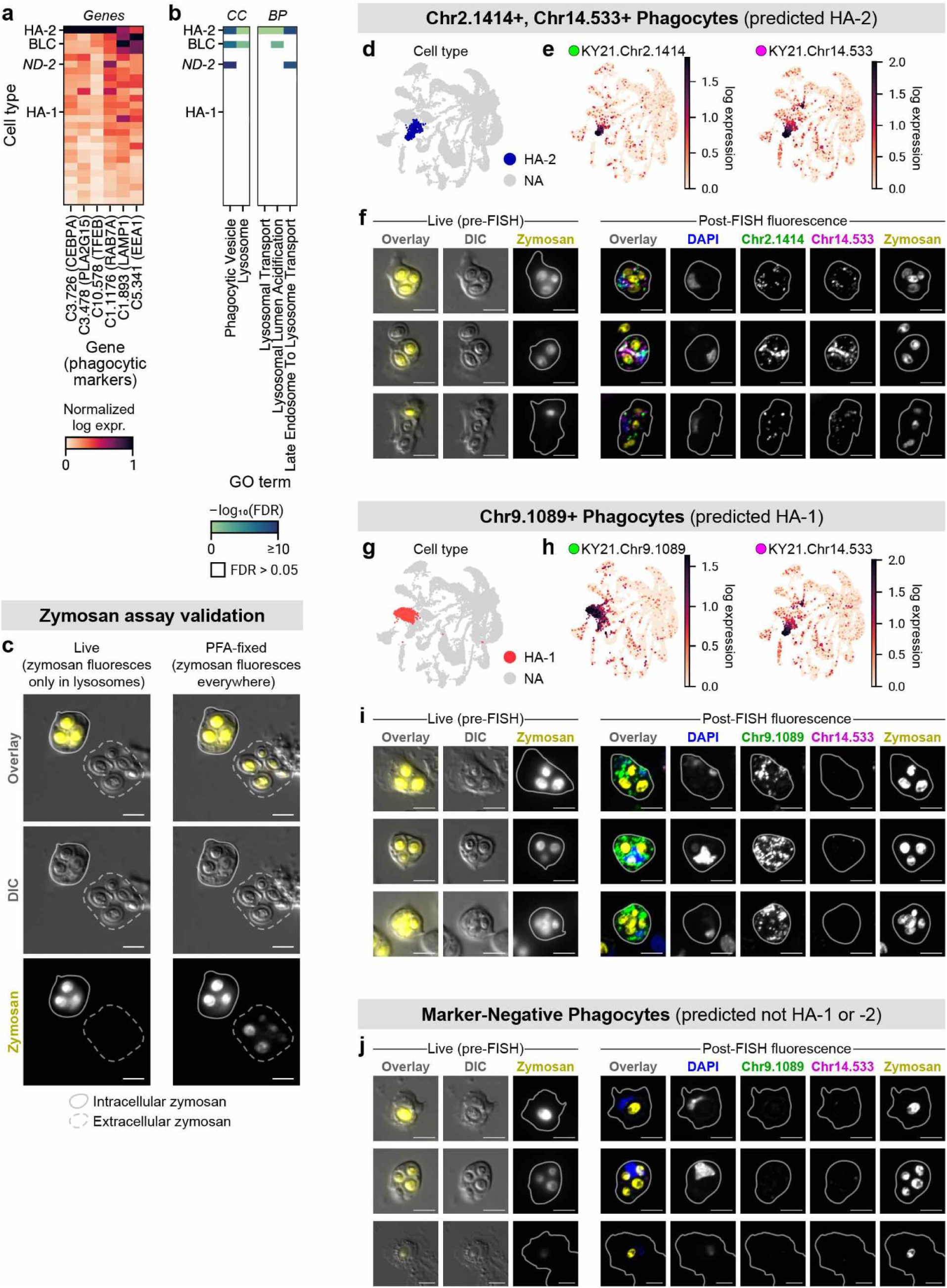
Phagocytosis in HA-2 and HA-1 **(a)** Expression of phagocytic markers in HA-2. **(b)** Phagocytosis-related Gene Ontology (GO) terms related to phagocytosis. Cellular Component (CC) and Biological Process (BP) terms are separated. See **Supplementary Table 18** for accession numbers. **(c)** Validation that only intracellular zymosan is fluorescent during live imaging. **(d, e, g, h)** UMAPs labeling the **(d)** HA-2 cluster, **(g)** HA-1 cluster, and **(e, h)** marker genes labeled with HCR FISH. **(f, i)** Example images showing cells with engulfed zymosan which are **(f)** positive for HA-2 markers (Chr14.533 and Chr2.1414) or **(i)** positive for the HA-1 marker (Chr9.1089) but negative for the HA-2 marker (Chr14.533). **(j)** Example images showing cells which are negative for both the HA-1 and HA-2 markers (Chr9.1089 and Chr14.533, respectively). Marker-negative cells accounted for 15% (42/282) of observed cells with engulfed zymosan.

**Supplementary Fig. 8:**
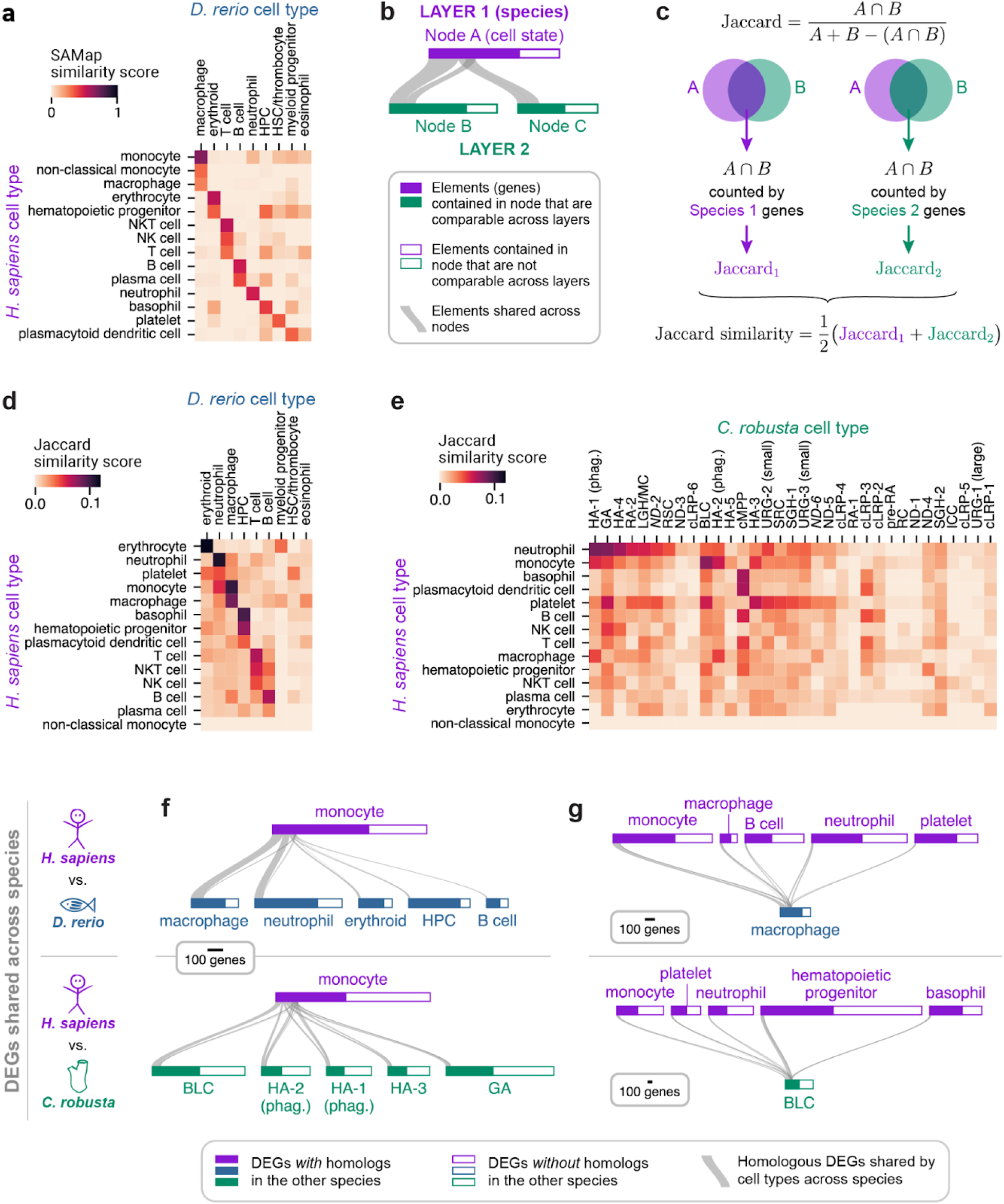
Comparison to vertebrate cell states **(a)** SAMap similarity matrix for the human to zebrafish comparison. **(b)** Cartoon of a generalized Sankey (GS) plot. See the Methods section (“Generalized Sankey (GS) Plots”) and the caption of Fig. 5c**,d** for a definition. **(c)** Diagram showing the calculation of a Jaccard similarity score between two cell states in two species, quantifying the overlap of DEGs while allowing for many-to-many gene homology. **(d, e)** Jaccard similarity score matrices for **(d)** human vs. zebrafish and **(e)** human vs. *C. robusta*. **(f, g)** GS plots comparing human monocytes to other species’ cell states. Panel **(f)** shows human monocytes vs. the most similar 5 zebrafish (top) or *C. robusta* (bottom) cell states. Panel **(d)** shows the most similar cell states in zebrafish (top) or *C. robusta* (bottom) to human monocytes vs. the most similar 5 human cell states.

**Supplementary Fig. 9:**
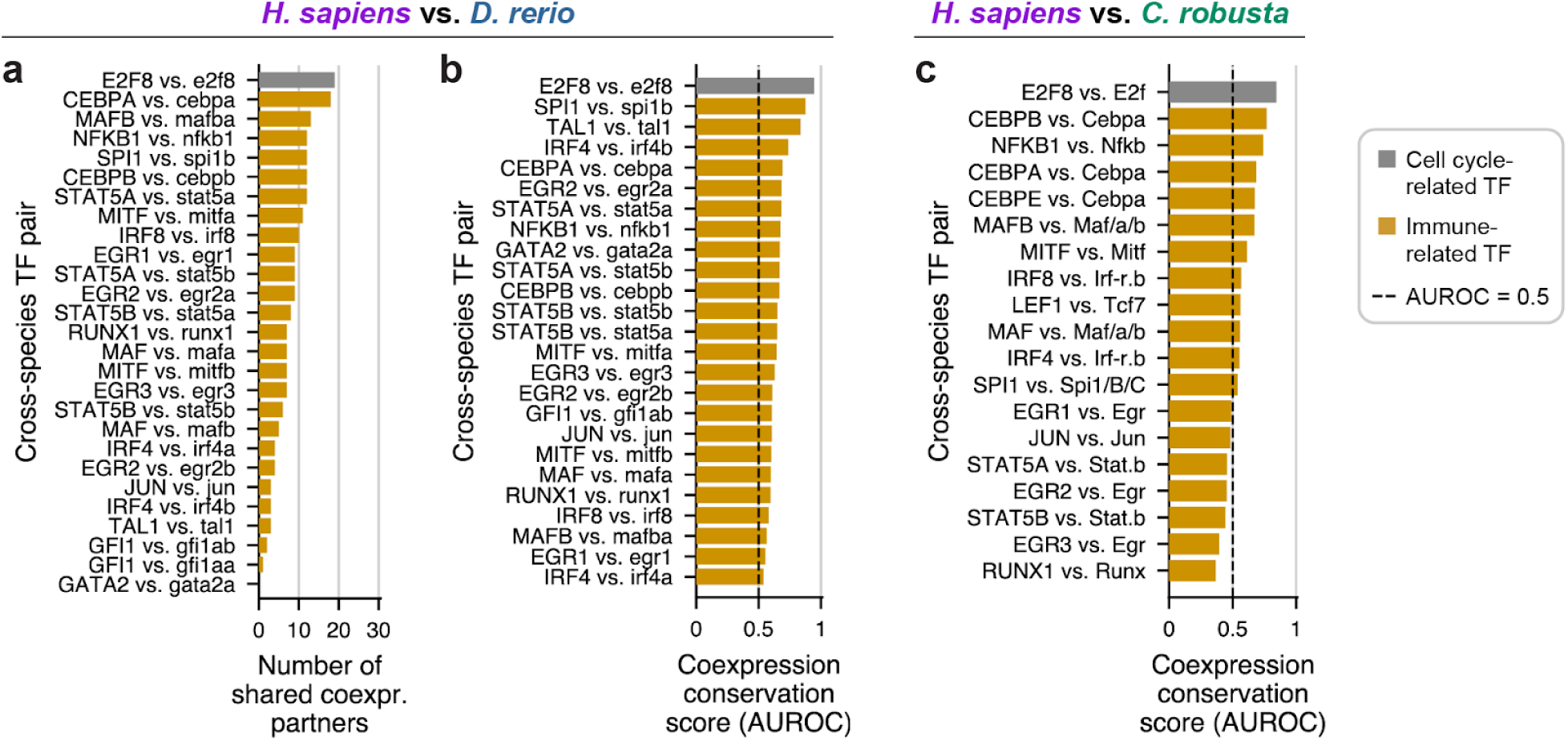
Comparison to vertebrate TFs & gene regulation Co-expression conservation scores for TF pairs in **(a,b)** zebrafish vs. human and **(c)** *C. robusta* vs. human. Scores in panel **(a)** quantify the number of co-expression partners shared across species. Scores in panels **(b)** and **(c)** calculate an AUROC-based score, for which scores at or below 0.5 indicate no significant co-expression conservation.

**Supplementary Fig. 10:**
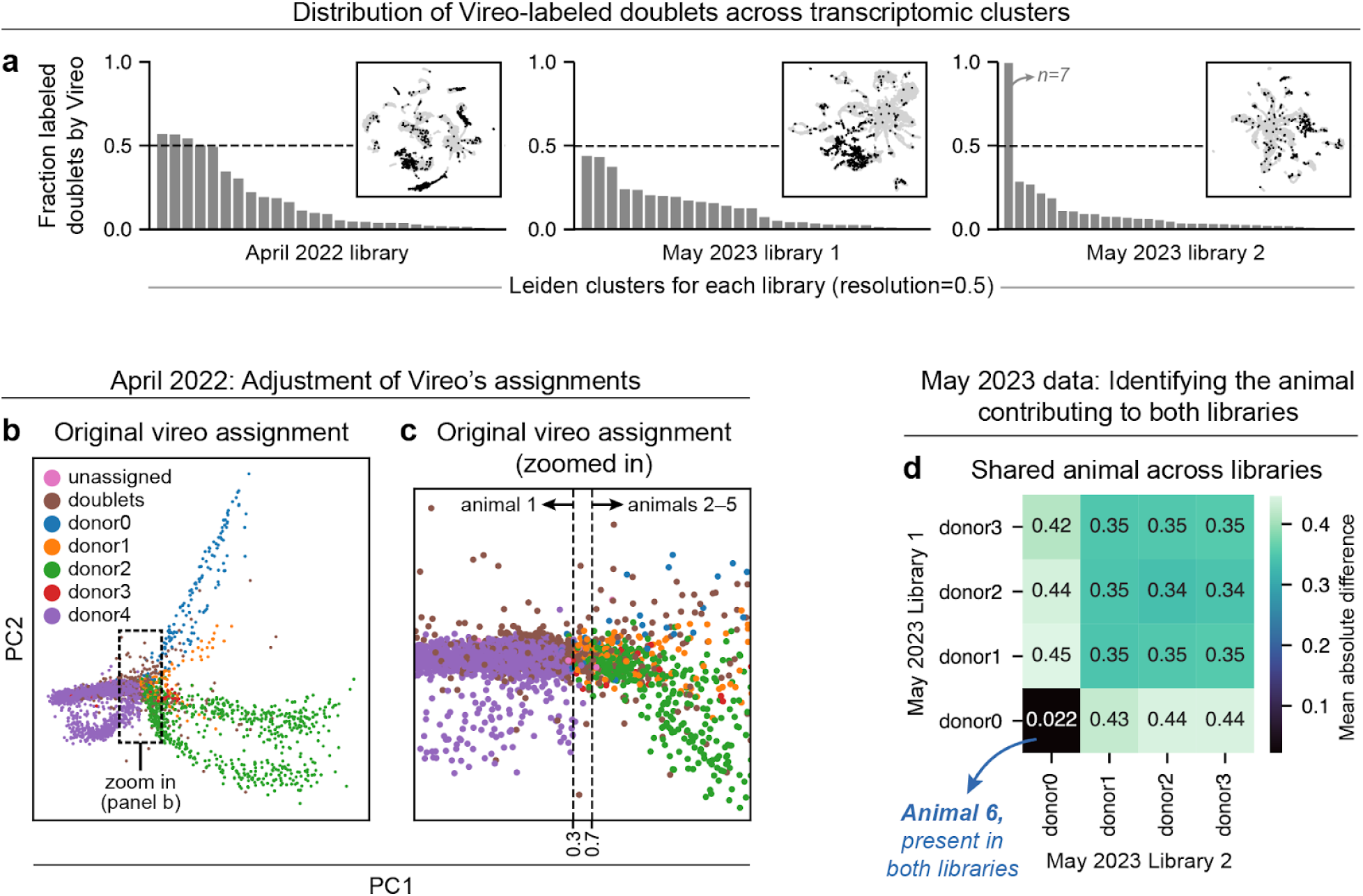
SNP demultiplexing **(a)** The fraction per cell cluster in each library that Vireo labeled as a doublet. The April 2022 library had four clusters in which over half of cells were labeled as doublets. One cluster in the May 2023 library 2 has 100% doublets, but since this cell cluster contains only 7 cells, it does not represent a large absolute number of called doublets. Inset: UMAP of each library individually, with cells colored as black if Vireo labeled them doublets. **(b, c)** Cell SNP profiles from the April 2022 dataset, plotted on the first two PCs defined by SNPs from a subset of transcriptional clusters. Cells are colored by the original Vireo animal assignments. The dashed box in **(a)** indicates the region plotted in **(b)**. The dotted lines in **(b)** represent the thresholds chosen for labeling cells as belonging to animal 1 or animals 2–5. **(d)** Mean absolute differences in genotype assignments between Vireo-assigned animals in the two May 2023 libraries. The small difference score for donor0 in library 1 and donor0 in library 2 indicates these are cells from the same animal.

**Supplementary Fig. 11:**
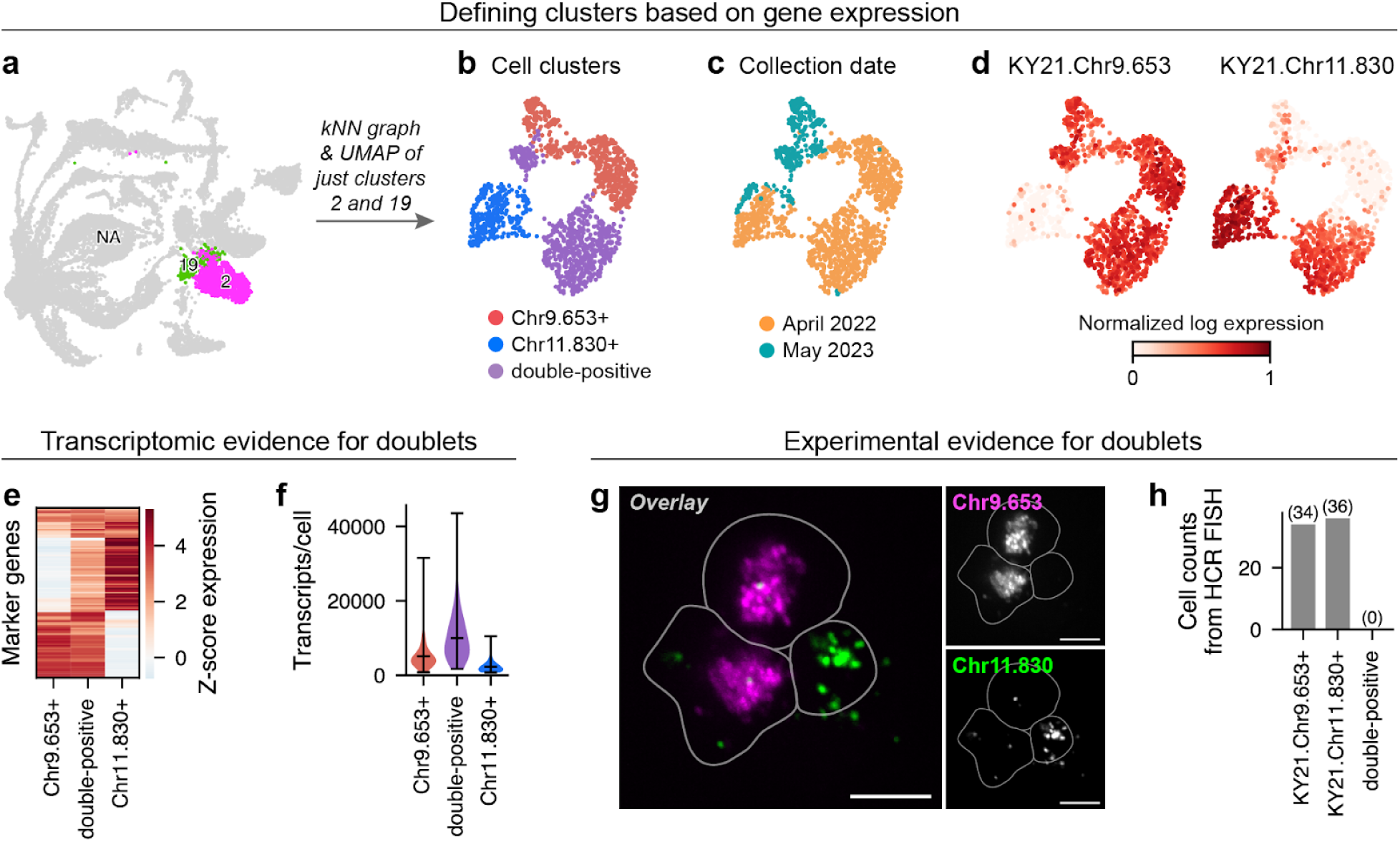
Doublet cluster identification and evidence **(a)** UMAP of the scRNA-seq dataset before removal of the doublet cluster. The two Leiden clusters, 2 and 19, contain the doublet cluster and the singlet clusters of cell states that comprise it. **(b–d)** For cells from clusters 2 and 19 in (a), a new *k-*nearest neighbor graph and UMAP embedding was generated. Cells are colored by **(b)** assignment to Chr9.653+, Chr11.830+ or double-positive clusters; **(c)** experimental day; **(d)** or expression of marker genes KY21.Chr9.653 (left) and KY21.Chr11.830 (right). **(e)** Genes which are enriched in at least one of Chr9.653+, Chr11.830+, or double-positive cells. Specifically, highly variable genes are included if they have a z-scored mean log-transformed expression above 2 in at least one of the three clusters. The genes are hierarchically clustered (seaborn.clustermap, metric=‘cosine’). **(f)** Violin plots showing the number of UMIs per cell barcode for each cell cluster. Black ticks indicate the mean and extrema. **(g)** Example image from HCR FISH experiments. We followed the HCR FISH protocol described in the Methods section (see “Cell preparation for HCR FISH with live imaging” and “HCR FISH for cluster annotation”), with the following change to reduce cell clumping: when blood cells were added to the 8-well coverslip they were diluted in CMF-ASW with 10 mM EDTA (ethylenediaminetetraacetic acid, Avantor EM-4050) instead of just CMF-ASW. **(h)** Cell counts from HCR FISH, showing zero detected double-positive cells. To assign a p-value, we define the null hypothesis as the observation of cell state abundances reflected in the scRNA-seq data: 43.6% of cells expressing either Chr9.653 or Chr11.830 would be double-marker-positive. Given this null, the probability of observing 0 double-marker-positive cells out of 70 observations is 3.8e-18. Even if cell state abundances in the scRNA-seq data set are inaccurate by ten fold, this probability would be 0.044. The p-value is therefore under 0.05.

**Supplementary Fig. 12:**
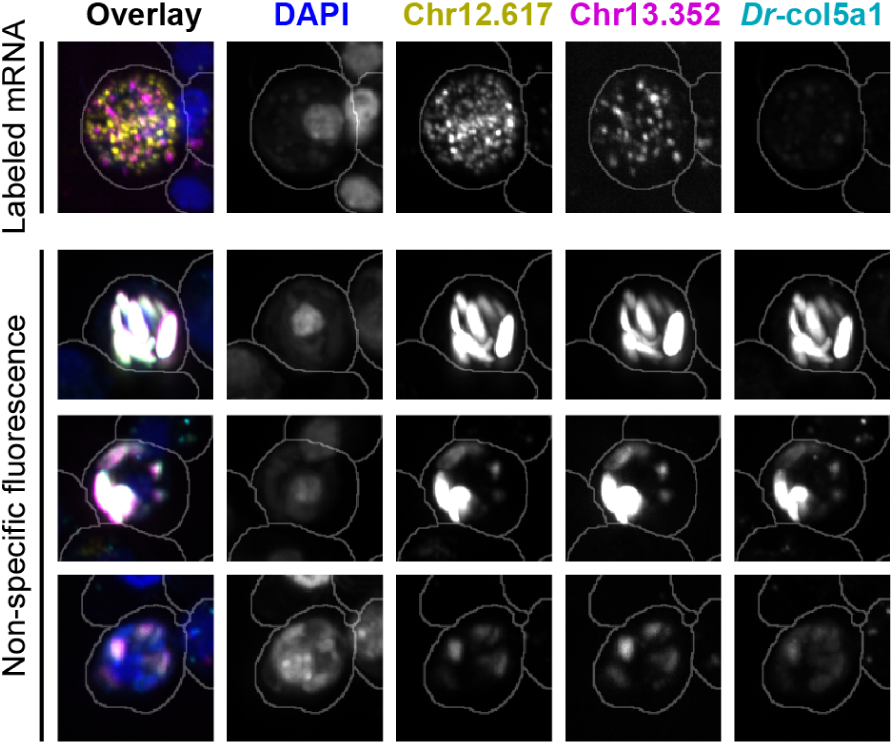
Non-specific fluorescence in HCR FISH experiments Demonstration of non-specific fluorescence in HCR FISH experiments via labeling zebrafish *col5a1*. In all observed cases, non-specific fluorescence was overlapping in all channels with HCR FISH labeling.

### Supplementary Videos

**Supp. Video 1:** HA-3 reporter in a whole juvenile HA-3 reporter-positive cells circulate in a juvenile. Green represents the HA-3 reporter, *Chr8.361>GFP*. Magenta represents a heart reporter, *Mesp>RFP*. *Separate AVI file*.

**Supp. Video 2:** HA-1 reporter in a whole juvenile HA-1 reporter-positive cells circulate in a juvenile. Green represents the HA-3 reporter, *Chr1.1154>GFP*. *Separate AVI file*.

**Supp. Video 3:** URG-1 reporter in a whole juvenile URG-1 reporter-positive cells circulate in a juvenile. Green represents the HA-3 reporter, *Chr2.603>GFP*. *Separate AVI file*.

**Supp. Video 4:** cLRP-1 reporter in a whole juvenile cLRP-1 reporter-positive cells circulate in a juvenile. Green represents the HA-3 reporter, *Chr4.869>GFP*. *Separate AVI file*.

**Supp. Video 5:** GA reporter-positive cell in a juvenile GA reporter-positive cell crawling in a juvenile. Green represents the HA-3 reporter, *Chr13.352>GFP*. *Separate AVI file*.

**Supp. Video 6:** HA-2 reporter-positive cell in a juvenile HA-2 reporter-positive cell crawling in a juvenile. Green represents the HA-2 reporter, *Chr2.1414>GFP*. *Separate AVI file*.

**Supp. Video 7:** HA-3 reporter-positive cell in a juvenile HA-3 reporter-positive cell crawling in a juvenile. Green represents the HA-2 reporter, *Chr8.361>GFP*. *Separate AVI file*.

**Supp. Video 8:** HA-1 reporter-positive cell in a juvenile HA-1 reporter-positive cell crawling in a juvenile. Green represents the HA-2 reporter, *Chr1.1154>GFP*. *Separate AVI file*.

**Supp. Video 9:** cMPP/cLRP reporter during metamorphosis cMPP/cLRP reporter-positive cells during metamorphosis. Green represents the cMPP/cLRP reporter, *Chr4.593>GFP*, and is shown as a maximum-intensity projection across z-slices. *Separate AVI file*.

**Supp. Video 10:** cMPP/cLRP reporter in a whole juvenile cMPP/cLRP reporter-positive cells are static in the tunic and circulating in a juvenile. Green represents the cMPP/cLRP reporter, *Chr4.593>GFP*. Magenta represents a heart reporter, *Mesp>RFP*. *Separate AVI file*.

### Supplementary Tables

**Supp. Table 1:** Previously identified marker genes of *C. robusta* blood morphotypes

*Separate CSV file*.

**Supp. Table 2:** Conversion from Leiden clusters to annotated cell states.

| Cluster | Cell state abbr. | Cell state full name (full) | Notes |
| --- | --- | --- | --- |
| 0/14.1 | cMPP | Candidate multi-potent progenitor | Labeled as progenitor, median expression of MKI67 > 0. |
| 1/7 | cLRP-1 | Candidate lineage-restricted progenitor 1 | Labeled as progenitor, median expression of MKI67 > 0. |
| 2.1 | cLRP-6 | Candidate lineage-restricted progenitor 6 | Labeled as progenitor, median expression of MKI67 > 0. |
| 2.2 | cLRP-4 | Candidate lineage-restricted progenitor 4 | Labeled as progenitor, median expression of MKI67 > 0. |
| 3 | HA-1 (phag.) | Hyaline amoebocyte 1, phagocytic |  |
| 4 | pre-RA | Precursor of refractile amoebocyte |  |
| 5 | cLRP-2 | Candidate lineage-restricted progenitor 2 | Labeled as progenitor, median expression of MKI67 > 0. |
| 6 | cLRP-3 | Candidate lineage-restricted progenitor 3 | Labeled as progenitor, median expression of MKI67 > 0. |
| 8 | RA-1 | Refractile amoebocyte 1 |  |
| 9 | HA-2 (phag.) | Hyaline amoebocyte 2, phagocytic |  |
| 10 | RSC | Round spreading cell |  |
| 11 | BLC | Blebbing-like cell |  |
| 12 | GA | Granular amoebocyte |  |
| 13 | HA-3 | Hyaline amoebocyte 3 |  |
| 14.2 | ND-4 | Not determined 4 |  |
| 15 | HA-4 | Hyaline amoebocyte 4 |  |
| 16 | LGH/MC | Large granules hemocyte/morula cell |  |
| 17 | ND-1 | Not determined 1 |  |
| 18 | ICC | Irregular compartment cell |  |
| 19 | URG-1 (large) | Unilocular refractile granulocyte 1, large |  |
| 20 | cLRP-5 | Candidate lineage-restricted progenitor 5 | Labeled as progenitor, median expression of MKI67 > 0. |
| 21 | RC | Round cell |  |
| 22 | SRC | Signet ring cell |  |
| 23 | ND-2 | Not determined 2 |  |
| 24 | SGH-1 | Small granules hemocyte 1 |  |
| 25 | RA-2 | Refractile amoebocyte 2 |  |
| 26 | ND-3 | Not determined 3 |  |
| 27 | URG-2 (small) | Unilocular refractile granulocyte 2, small |  |
| 28 | URG-3 (small) | Unilocular refractile granulocyte 3, small |  |
| 29 | SGH-2 | Small granules hemocyte 2 |  |
| 30 | ND-5 | Not determined 4 |  |
| 31 | HA-5 | Hyaline amoebocyte 5 |  |
| 32 | ND-6 | Not determined 5 |  |

**Supp. Table 3:** Likely homologs of *C6* in *C. robusta*.

| KY21 ID | BLAST percent identity with human C6 | BLAST e-value with human C6 |
| --- | --- | --- |
| KY21.Chr1.1558 | 27.739 | 6.71e-54 |
| KY21.Chr1.1559 | 26.882 | 4.40e-52 |
| KY21.Chr1.1560 | 29.326 | 1.28e-64 |
| KY21.Chr3.1186 | 31.471 | 3.57e-77 |
| KY21.Chr3.1187 | 30.000 | 1.93e-77 |
| KY21.Chr3.1189 | 31.296 | 1.32e-80 |
| KY21.Chr10.252 | 33.090 | 1.69e-90 |
| KY21.Chr11.710 | 32.904 | 4.18e-83 |
| KY21.Chr11.711 | 31.304 | 1.04e-68 |
| KY21.Chr11.712 | 31.250 | 1.73e-56 |
| KY21.Chr11.714 | 31.876 | 4.00e-79 |
| KY21.Chr14.171 | 33.210 | 4.04e-85 |
| KY21.Chr14.172 | 32.959 | 5.01e-85 |

**Supp. Table 4:** Homology of *C. robusta* genes detected by OrthoFinder and BLAST For every gene, this table lists whether genes are identified as having a human homology by OrthoFinder (column “OrthoFinder homolog(s)?”), or as having any detected BLAST similarity to a human gene with an e-value below 1e-6 (column “BLAST similarity?”). *Separate CSV file*.

**Supp. Table 5:** List of cell state–specific genes with no detected vertebrate homolog For every cell state, a column lists the genes enriched in that cell state (Wilcoxon rank-sum test, FDR<0.05, log fold change > 2) with no detected vertebrate homolog. A gene is considered to have no detected vertebrate homolog if both columns in **Supplementary Table 4** are no, i.e. there is no OrthoFinder homolog and no human gene with a BLAST e-value below 1e-6.

*Separate CSV file*.

**Supp. Table 6:** List of annotated *C. robusta* TFs with names and KY21 IDs

*Separate CSV file*.

**Supp. Table 7:**
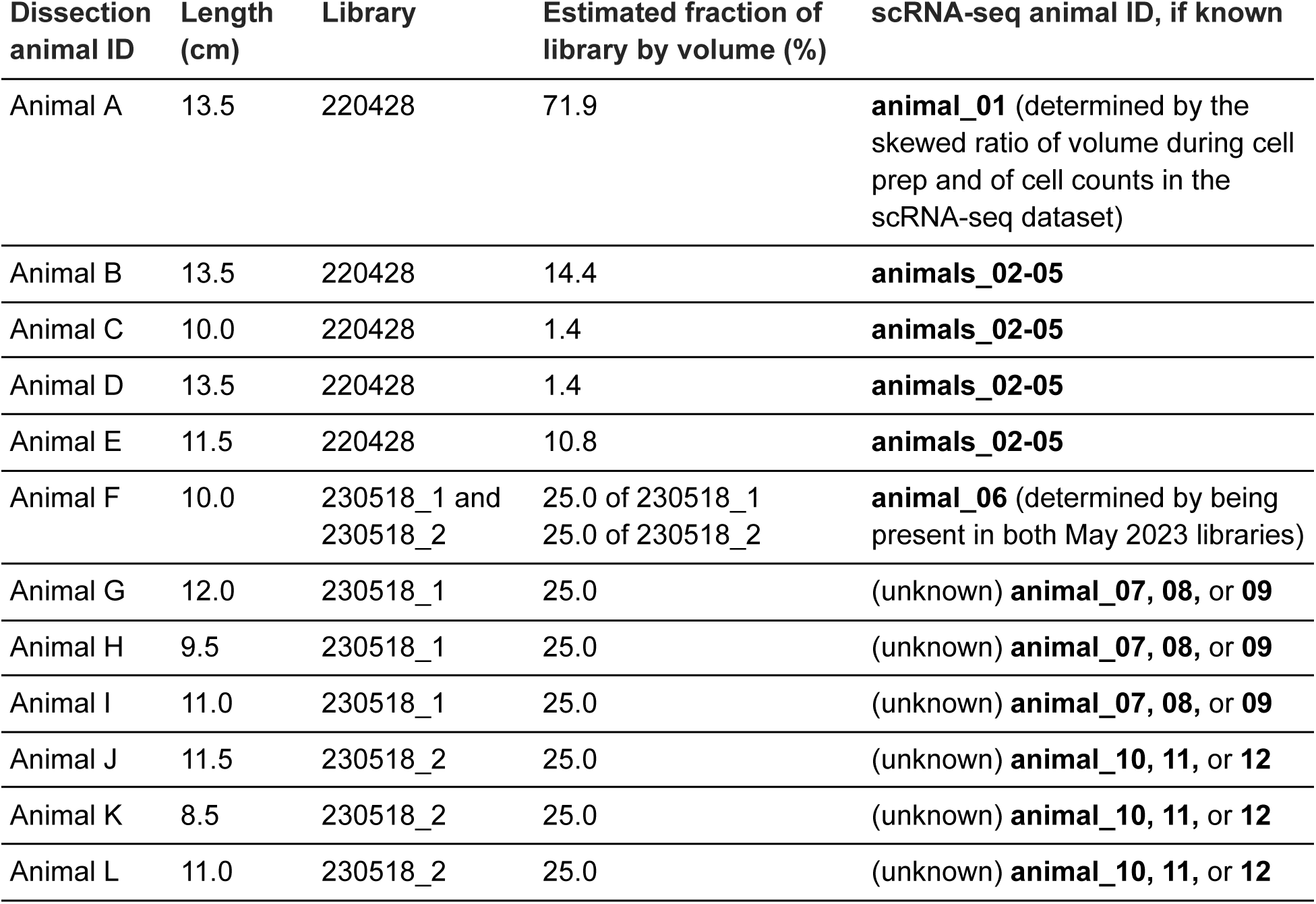
Animal metadata for the scRNA-seq experiment.

| Dissection animal ID | Length (cm) | Library | Estimated fraction of library by volume (%) | scRNA-seq animal ID, if known |
| --- | --- | --- | --- | --- |
| Animal A | 13.5 | 220428 | 71.9 | **animal_01** (determined by the skewed ratio of volume during cell prep and of cell counts in the scRNA-seq dataset) |
| Animal B | 13.5 | 220428 | 14.4 | **animals_02-05** |
| Animal C | 10.0 | 220428 | 1.4 | **animals_02-05** |
| Animal D | 13.5 | 220428 | 1.4 | **animals_02-05** |
| Animal E | 11.5 | 220428 | 10.8 | **animals_02-05** |
| Animal F | 10.0 | 230518_1 and 230518_2 | 25.0 of 230518_1<br>25.0 of 230518_2 | **animal_06** (determined by being present in both May 2023 libraries) |
| Animal G | 12.0 | 230518_1 | 25.0 | (unknown) **animal_07, 08, or 09** |
| Animal H | 9.5 | 230518_1 | 25.0 | (unknown) **animal_07, 08, or 09** |
| Animal I | 11.0 | 230518_1 | 25.0 | (unknown) **animal_07, 08, or 09** |
| Animal J | 11.5 | 230518_2 | 25.0 | (unknown) **animal_10, 11, or 12** |
| Animal K | 8.5 | 230518_2 | 25.0 | (unknown) **animal_10, 11, or 12** |
| Animal L | 11.0 | 230518_2 | 25.0 | (unknown) **animal_10, 11, or 12** |

**Supp. Table 8: Cell barcodes used to define the SNP PC space for the April 2022 library**

*Separate CSV file*.

**Supp. Table 9:** Cell barcodes labeled as likely doublets of SRC and HA-3

*Separate CSV file*.

**Supp. Table 10:** HCR FISH probe sets *Separate Excel file.* Sheet 1 summarizes probe pool names with HCR amplifier sequences. Sheet 2 lists the probe sequences for each probe pool.

**Supp. Table 11:** HCR FISH probes and fluorescent hairpin combinations used

*Separate CSV file*.

**Supp. Table 12:**
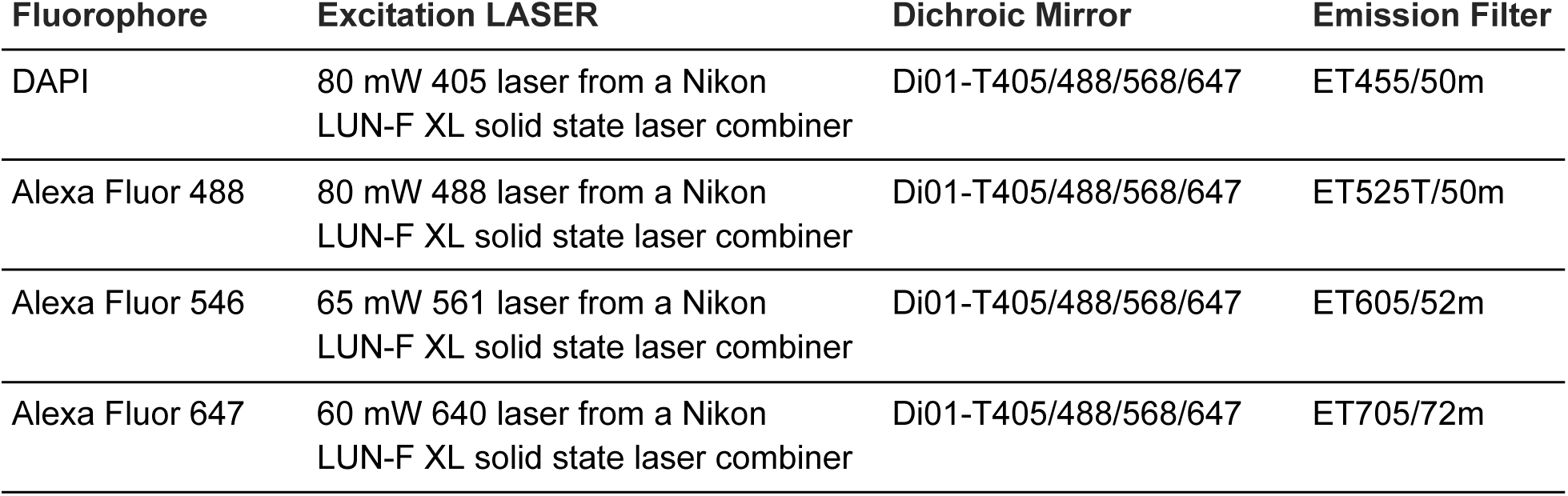
Confocal imaging information.

**Supp. Table 13: Live reporter constructs**

*Separate CSV file*.

**Supp. Table 14:**
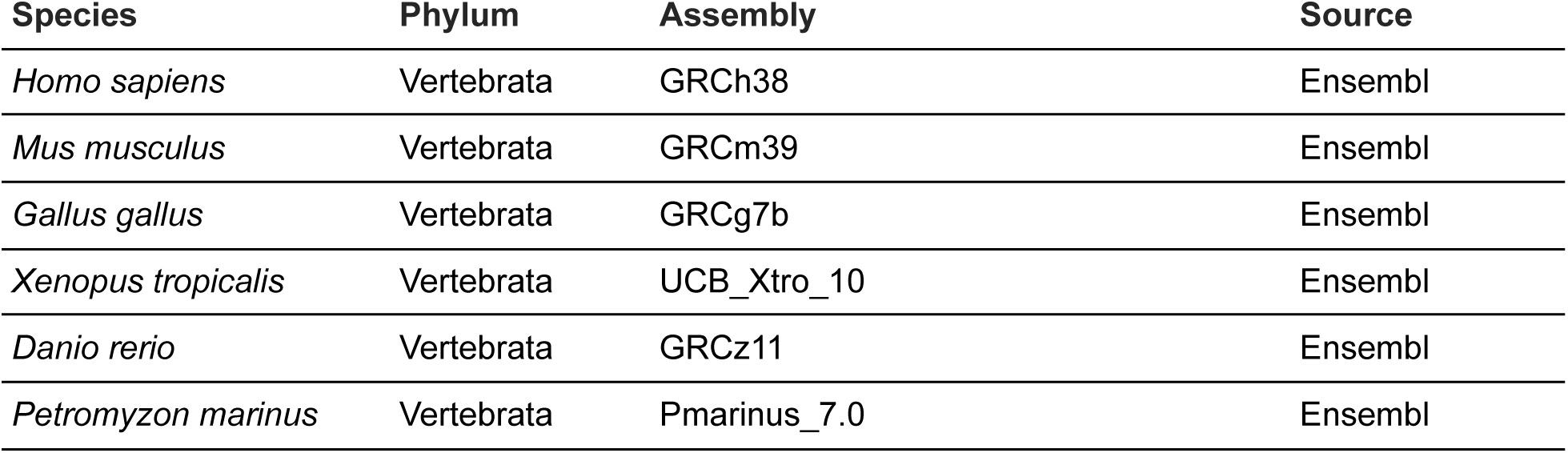
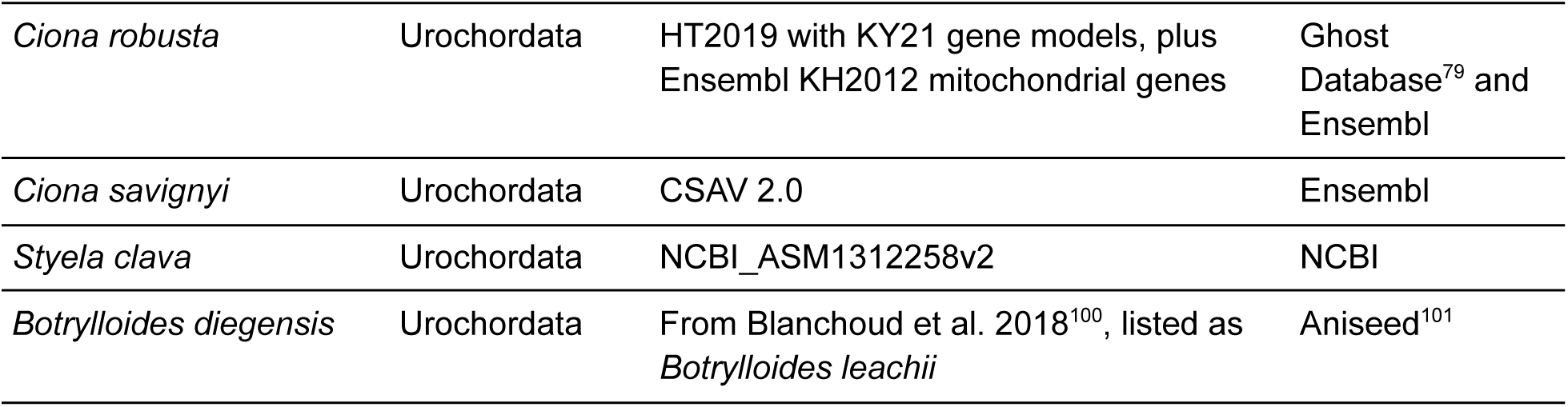
Proteomes used to run OrthoFinder and BLAST.

| Species | Phylum | Assembly | Source |
| --- | --- | --- | --- |
| *Homo sapiens* | Vertebrata | GRCh38 | Ensembl |
| *Mus musculus* | Vertebrata | GRCm39 | Ensembl |
| *Gallus gallus* | Vertebrata | GRCg7b | Ensembl |
| *Xenopus tropicalis* | Vertebrata | UCB_Xtro_10 | Ensembl |
| *Danio rerio* | Vertebrata | GRCz11 | Ensembl |
| *Petromyzon marinus* | Vertebrata | Pmarinus_7.0 | Ensembl |
| *Ciona robusta* | Urochordata | HT2019 with KY21 gene models, plus Ensembl KH2012 mitochondrial genes | Ghost Database<sup>79</sup> and Ensembl |
| *Ciona savignyi* | Urochordata | CSAV 2.0 | Ensembl |
| *Styela clava* | Urochordata | NCBI_ASM1312258v2 | NCBI |
| *Botrylloides diegensis* | Urochordata | From Blanchoud et al. 2018<sup>100</sup>, listed as *Botrylloides leachii* | Aniseed<sup>101</sup> |

**Supp. Table 15:** Sources for human immune gene lists.

| Figure panel | Gene set | Source(s) |
| --- | --- | --- |
| Fig. 4h | Complement activators | HUGO gene group<sup>102</sup>: Complement system activation components |
| Supplementary Fig. 6e | Pattern recognition receptors | PRRDB 2.0<sup>103</sup> |
| Supplementary Fig. 6f | Cytokines | Hanrahan et al.<sup>104</sup><br>HUGO gene groups<sup>102</sup>: Interleukins (IL), Interferons (IFN), Tumor necrosis factor superfamily (TNFSF), Transforming growth factor beta family (TGFB), Chemokine ligands (CCL) |
| Supplementary Fig. 6g | Cytokine receptors | Hanrahan et al.<sup>104</sup><br>HUGO gene groups<sup>102</sup>: Interleukin receptors, Interferon receptors, Tumor necrosis factor receptor superfamily, Chemokine receptors |

**Supp. Table 16: Renaming of cell state annotations in the Tabula Sapiens dataset**

*Separate CSV file*.

**Supp. Table 17:** Cell type annotations for each cell barcode in the zebrafish dataset

*Separate CSV file*.

**Supp. Table 18.**
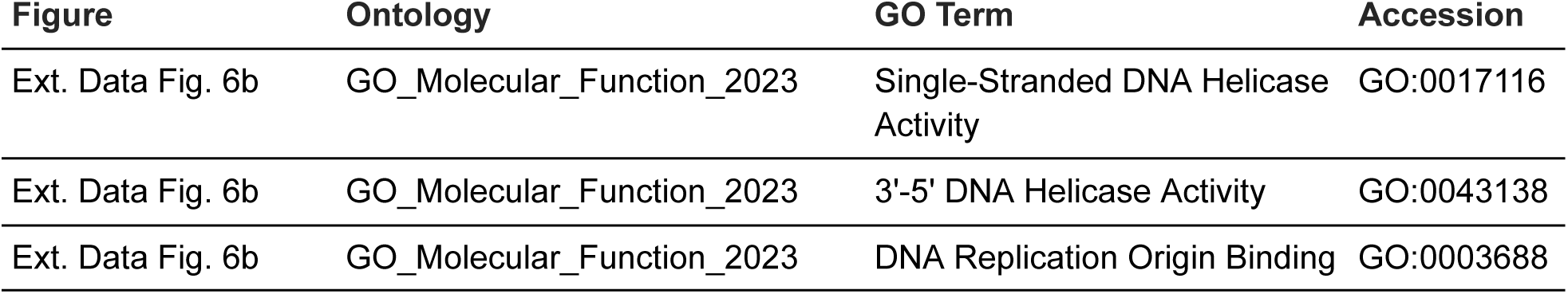
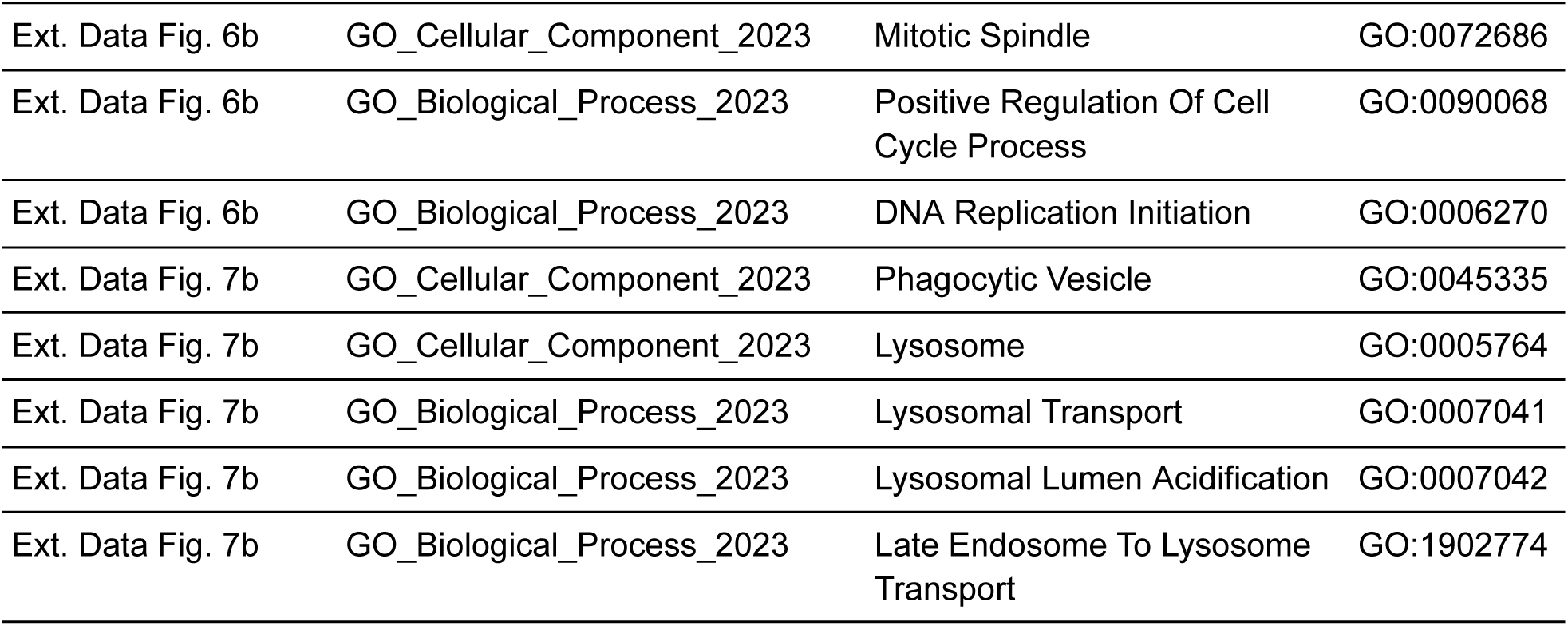
GO term accession numbers.

| Figure | Ontology | GO Term | Accession |
| --- | --- | --- | --- |
| Ext. Data Fig. 6b | GO_Molecular_Function_2023 | Single-Stranded DNA Helicase Activity | GO:0017116 |
| Ext. Data Fig. 6b | GO_Molecular_Function_2023 | 3'-5' DNA Helicase Activity | GO:0043138 |
| Ext. Data Fig. 6b | GO_Molecular_Function_2023 | DNA Replication Origin Binding | GO:0003688 |

| **Figure** | **Ontology** | **GO Term** | **Accession** |
| --- | --- | --- | --- |
| Ext. Data Fig. 6b | GO_Cellular_Component_2023 | Mitotic Spindle | GO:0072686 |
| Ext. Data Fig. 6b | GO_Biological_Process_2023 | Positive Regulation Of Cell Cycle Process | GO:0090068 |
| Ext. Data Fig. 6b | GO_Biological_Process_2023 | DNA Replication Initiation | GO:0006270 |
| Ext. Data Fig. 7b | GO_Cellular_Component_2023 | Phagocytic Vesicle | GO:0045335 |
| Ext. Data Fig. 7b | GO_Cellular_Component_2023 | Lysosome | GO:0005764 |
| Ext. Data Fig. 7b | GO_Biological_Process_2023 | Lysosomal Transport | GO:0007041 |
| Ext. Data Fig. 7b | GO_Biological_Process_2023 | Lysosomal Lumen Acidification | GO:0007042 |
| Ext. Data Fig. 7b | GO_Biological_Process_2023 | Late Endosome To Lysosome Transport | GO:1902774 |

## Bibliography

1. Arendt, D., Musser, J.M., Baker, C.V.H., Bergman, A., Cepko, C., Erwin, D.H., Pavlicev, M., Schlosser, G., Widder, S., Laubichler, M.D., et al. (2016). The origin and evolution of cell types. Nature Reviews Genetics 17, 744–757.

2. Babonis, L.S., Enjolras, C., Ryan, J.F., and Martindale, M.Q. (2022). A novel regulatory gene promotes novel cell fate by suppressing ancestral fate in the sea anemone Nematostella vectensis. Proc. Natl. Acad. Sci. U. S. A. 119, e2113701119.

3. Pomaville, M.B., Sattler, S.M., and Abitua, P.B. (2024). A new dawn for the study of cell type evolution. Development 151, dev200884.

4. Cooper, E.L. (1976). Evolution of blood cells. Ann. Inst. Pasteur Immunol. 127, 817–825.

5. David, C.N., Ozbek, S., Adamczyk, P., Meier, S., Pauly, B., Chapman, J., Hwang, J.S., Gojobori, T., and Holstein, T.W. (2008). Evolution of complex structures: minicollagens shape the cnidarian nematocyst. Trends Genet. 24, 431–438.

6. Pillai, A.S., Chandler, S.A., Liu, Y., Signore, A.V., Cortez-Romero, C.R., Benesch, J.L.P., Laganowsky, A., Storz, J.F., Hochberg, G.K.A., and Thornton, J.W. (2020). Origin of complexity in haemoglobin evolution. Nature 581, 480–485.

7. Brückner, A., Badroos, J.M., Learsch, R.W., Yousefelahiyeh, M., Kitchen, S.A., and Parker, J. (2021). Evolutionary assembly of cooperating cell types in an animal chemical defense system. Cell 184, 6138–6156.e28.

8. Müller, W., Blumbach, B., and Müller, I. (1999). Evolution of the innate and adaptive immune systems: relationships between potential immune molecules in the lowest metazoan phylum (Porifera) and those in vertebrates. Transplantation 68, 1215–1227.

9. Vandepas, L.E., Stefani, C., Domeier, P.P., Traylor-Knowles, N., Goetz, F.W., Browne, W.E., and Lacy-Hulbert, A. (2024). Extracellular DNA traps in a ctenophore demonstrate immune cell behaviors in a non-bilaterian. Nat. Commun. 15, 2990.

10. Rast, J.P., and Messier-Solek, C. (2008). Marine invertebrate genome sequences and our evolving understanding of animal immunity. Biol. Bull. 214, 274–283.

11. Bradley, J.E., and Jackson, J.A. (2008). Measuring immune system variation to help understand host-pathogen community dynamics. Parasitology 135, 807–823.

12. Guryanova, S.V., Balandin, S.V., Belogurova-Ovchinnikova, O.Y., and Ovchinnikova, T.V. (2023). Marine invertebrate antimicrobial peptides and their potential as novel peptide antibiotics. Mar. Drugs 21, 503.

13. Ebert, D., and Fields, P.D. (2020). Host-parasite co-evolution and its genomic signature. Nat. Rev. Genet. 21, 754–768.

14. Melillo, D., Marino, R., Italiani, P., and Boraschi, D. (2018). Innate immune memory in invertebrate metazoans: A critical appraisal. Front. Immunol. 9, 1915.

15. Leulier, F., and Lemaitre, B. (2008). Toll-like receptors--taking an evolutionary approach. Nat. Rev. Genet. 9, 165–178.

16. Little, T.J., Hultmark, D., and Read, A.F. (2005). Invertebrate immunity and the limits of mechanistic immunology. Nat. Immunol. 6, 651–654.

17. Buckley, K.M., and Rast, J.P. (2012). Dynamic evolution of toll-like receptor multigene families in echinoderms. Front. Immunol. 3, 136.

18. Prochazkova, P., Roubalova, R., Dvorak, J., Navarro Pacheco, N.I., and Bilej, M. (2020). Pattern recognition receptors in annelids. Dev. Comp. Immunol. 102, 103493.

19. Wiens, G.D., and Glenney, G.W. (2011). Origin and evolution of TNF and TNF receptor superfamilies. Dev. Comp. Immunol. 35, 1324–1335.

20. Huang, X.-D., Zhang, H., and He, M.-X. (2015). Comparative and evolutionary analysis of the interleukin 17 gene family in invertebrates. PLoS One 10, e0132802.

21. Dishaw, L.J., Smith, S.L., and Bigger, C.H. (2005). Characterization of a C3-like cDNA in a coral: phylogenetic implications. Immunogenetics 57, 535–548.

22. Wang, X.-W., and Wang, J.-X. (2013). Pattern recognition receptors acting in innate immune system of shrimp against pathogen infections. Fish Shellfish Immunol. 34, 981–989.

23. Peng, M., Li, Z., Cardoso, J.C.R., Niu, D., Liu, X., Dong, Z., Li, J., and Power, D.M. (2022). Domain-Dependent Evolution Explains Functional Homology of Protostome and Deuterostome Complement C3-Like Proteins. Front. Immunol. 13, 840861.

24. 24. Ratcliff, N.A., and Rowley, A.F. (1981). Invertebrate blood cells (Academic Press Inc.).

25. Kaufmann, S.H.E. (2008). Immunology’s foundation: the 100-year anniversary of the Nobel Prize to Paul Ehrlich and Elie Metchnikoff. Nat. Immunol. 9, 705–712.

26. Nagahata, Y., Masuda, K., Nishimura, Y., Ikawa, T., Kawaoka, S., Kitawaki, T., Nannya, Y., Ogawa, S., Suga, H., Satou, Y., et al. (2022). Tracing the evolutionary history of blood cells to the unicellular ancestor of animals. Blood 140, 2611–2625.

27. Carrau, T., Thümecke, S., Silva, L.M.R., Perez-Bravo, D., Gärtner, U., Taubert, A., Hermosilla, C., Vilcinskas, A., and Lee, K.-Z. (2021). The cellular innate immune response of the invasive pest insect Drosophila suzukii against Pseudomonas entomophila involves the release of extracellular traps. Cells 10, 3320.

28. Ballarin, L., Cima, F., and Sabbadin, A. (1998). Phenoloxidase and cytotoxicity in the compound ascidian Botryllus schlosseri. Dev. Comp. Immunol. 22, 479–492.

29. Evans, C.J., Hartenstein, V., and Banerjee, U. (2003). Thicker than blood: conserved mechanisms in Drosophila and vertebrate hematopoiesis. Dev. Cell 5, 673–690.

30. Coates, J.A., Brooks, E., Brittle, A.L., Armitage, E.L., Zeidler, M.P., and Evans, I.R. (2021). Identification of functionally distinct macrophage subpopulations in Drosophila. Elife 10. 10.7554/eLife.58686.

31. Yang, P., Chen, Y., Huang, Z., Xia, H., Cheng, L., Wu, H., Zhang, Y., and Wang, F. (2022). Single-cell RNA sequencing analysis of shrimp immune cells identifies macrophage-like phagocytes. Elife 11. 10.7554/eLife.80127.

32. 32. Casey, M.K.W. (2017). Janeway’s Immunobiology (Garland Science, Taylor & Francis Group, LLC).

33. Hartenstein, V. (2006). Blood Cells and Blood Cell Development in the Animal Kingdom. Annu. Rev. Cell Dev. Biol. 22, 677–712.

34. Longo, V., Parrinello, D., Longo, A., Parisi, M.G., Parrinello, N., Colombo, P., and Cammarata, M. (2021). The conservation and diversity of ascidian cells and molecules involved in the inflammatory reaction: The Ciona robusta model. Fish Shellfish Immunol. 119, 384–396.

35. Delsuc, F., Philippe, H., Tsagkogeorga, G., Simion, P., Tilak, M.-K., Turon, X., López-Legentil, S., Piette, J., Lemaire, P., and Douzery, E.J.P. (2018). A phylogenomic framework and timescale for comparative studies of tunicates. BMC Biol. 16, 39.

36. Millar, R.H. (1953). Ciona J. S. Colman, ed. (University Press of Liverpool).

37. Goodbody, I. (1974). The physiology of ascidians. Adv. Mar. Biol. 12, 1–149.

38. Cima, F., Franchi, N., and Ballarin, L. (2016). Origin and functions of tunicate hemocytes. In The Evolution of the Immune System: Conservation and Diversification (Elsevier Inc.), pp. 29–49.

39. 39. de Leo, G. (1992). Ascidian hemocytes and their involvement in defence reactions. Boll. Zool. 59, 195–214.

40. Ballarin, L., Cima, F., and Sabbadin, A. (1995). Morula Cells and Histocompatibility in the Colonial Ascidian Botryllus schlosseri. jzoo 12, 757–764.

41. Rosental, B., Kowarsky, M., Seita, J., Corey, D.M., Ishizuka, K.J., Palmeri, K.J., Chen, S.-Y., Sinha, R., Okamoto, J., Mantalas, G., et al. (2018). Complex mammalian-like haematopoietic system found in a colonial chordate. Nature 564, 425–429.

42. 42. What is your conceptual definition of “cell type” in the context of a mature organism? (2017). Cell Syst. 4, 255–259.

43. Dance, A. (2024). What is a cell type, really? The quest to categorize life’s myriad forms. Nature 633, 754–756.

44. Rowley, A.F. (1981). The blood cells of the sea squirt, Ciona intestinalis: Morphology, differential counts, and in vitro phagocytic activity. J. Invertebr. Pathol. 37, 91–100.

45. Parrinello, D., Parisi, M., Parrinello, N., and Cammarata, M. (2020). Ciona robusta hemocyte populational dynamics and PO-dependent cytotoxic activity. Dev. Comp. Immunol. 103, 103519.

46. 46. Zeng, F., Peronato, A., Ballarin, L., and Rothbächer, U. (2022). Sweet Tunicate Blood Cells: A Glycan Profiling of Haemocytes in Three Ascidian Species. In Advances in Aquatic Invertebrate Stem Cell Research: From Basic Research to Innovative Applications, L. Ballarin, B. Rinkevich, and B. Hobmayer, eds. (MDPI), pp. 351–379.

47. Scully, T., and Klein, A. (2023). A mannitol-based buffer improves single-cell RNA sequencing of high-salt marine cells. bioRxiv, 2023.04.26.538465. 10.1101/2023.04.26.538465.

48. Huang, X., and Huang, Y. (2021). Cellsnp-lite: an efficient tool for genotyping single cells. Bioinformatics 37, 4569–4571.

49. Huang, Y., McCarthy, D.J., and Stegle, O. (2019). Vireo: Bayesian demultiplexing of pooled single-cell RNA-seq data without genotype reference. Genome Biol. 20, 273.

50. Tusi, B.K., Wolock, S.L., Weinreb, C., Hwang, Y., Hidalgo, D., Zilionis, R., Waisman, A., Huh, J.R., Klein, A.M., and Socolovsky, M. (2018). Population snapshots predict early haematopoietic and erythroid hierarchies. Nature 555, 54–60.

51. Choi, H.M.T., Schwarzkopf, M., Fornace, M.E., Acharya, A., Artavanis, G., Stegmaier, J., Cunha, A., and Pierce, N.A. (2018). Third-generation in situ hybridization chain reaction: multiplexed, quantitative, sensitive, versatile, robust. Development 145. 10.1242/dev.165753.

52. Hotta, K., Dauga, D., and Manni, L. (2020). The ontology of the anatomy and development of the solitary ascidian Ciona: the swimming larva and its metamorphosis. Sci. Rep. 10, 17916.

53. La Manno, G., Soldatov, R., Zeisel, A., Braun, E., Hochgerner, H., Petukhov, V., Lidschreiber, K., Kastriti, M.E., Lönnerberg, P., Furlan, A., et al. (2018). RNA velocity of single cells. Nature 560, 494–498.

54. Wagner, D.E., Weinreb, C., Collins, Z.M., Briggs, J.A., Megason, S.G., and Klein, A.M. (2018). Single-cell mapping of gene expression landscapes and lineage in the zebrafish embryo. Science 360, 981–987.

55. Nishida, H., and Hirano, T. (1998). Developmental fates of larval tissues after metamorphosis in ascidian Halocynthia roretzi. I. Origin of mesodermal tissues of the juvenile Developmental Fates of Larval Tissues after Metamorphosis in Ascidian Halocynthia roretzi. Article in Developmental Biology 192, 199–210.

56. Tokuoka, M., Imai, K.S., Satou, Y., and Satoh, N. (2004). Three distinct lineages of mesenchymal cells in Ciona intestinalis embryos demonstrated by specific gene expression. Dev. Biol. 274, 211–224.

57. Totsuka, N.M., Kuwana, S., Sawai, S., Oka, K., Sasakura, Y., and Hotta, K. (2023). Distribution changes of non-self-test cells and self-tunic cells surrounding the outer body during Ciona metamorphosis. Dev. Dyn. 252, 1363–1374.

58. 58. Webb, B.Y.D.A. (1939). Observations on the blood of certain ascidians, with special reference to the biochemistry of vanadium. https://jeb.biologists.org/content/jexbio/16/4/499.full.pdf.

59. Michibata, H. (1996). The mechanism of accumulation of vanadium by ascidians: Some progress towards an understanding of this unusual phenomenon. Zoolog. Sci. 13, 489–502.

60. Trivedi, S., Ueki, T., Yamaguchi, N., and Michibata, H. (2003). Novel vanadium-binding proteins (vanabins) identified in cDNA libraries and the genome of the ascidian Ciona intestinalis. Biochim. Biophys. Acta 1630, 64–70.

61. Ebner, B., Burmester, T., and Hankeln, T. (2003). Globin Genes Are Present in Ciona intestinalis. Mol. Biol. Evol. 20, 1521–1525.

62. Geers, C., and Gros, G. (2000). Carbon dioxide transport and carbonic anhydrase in blood and muscle. Physiol. Rev. 80, 681–715.

63. Zhou, Z., Xu, M.-J., and Gao, B. (2016). Hepatocytes: a key cell type for innate immunity. Cell. Mol. Immunol. 13, 301–315.

64. Tabula Sapiens Consortium*, Jones, R.C., Karkanias, J., Krasnow, M.A., Pisco, A.O., Quake, S.R., Salzman, J., Yosef, N., Bulthaup, B., Brown, P., et al. (2022). The Tabula Sapiens: A multiple-organ, single-cell transcriptomic atlas of humans. Science 376, eabl4896.

65. Emms, D.M., and Kelly, S. (2019). OrthoFinder: phylogenetic orthology inference for comparative genomics. Genome Biol. 20, 238.

66. Tarashansky, A.J., Musser, J.M., Khariton, M., Li, P., Arendt, D., Quake, S.R., and Wang, B. (2021). Mapping single-cell atlases throughout Metazoa unravels cell type evolution. Elife 10. 10.7554/eLife.66747.

67. Singh, P.P., and Isambert, H. (2020). OHNOLOGS v2: A comprehensive resource for the genes retained from whole genome duplication in vertebrates. Nucleic Acids Res. 48, D724–D730.

68. Wang, H.-Y., Chen, J.-Y., Li, Y., Zhang, X., Liu, X., Lu, Y., He, H., Li, Y., Chen, H., Liu, Q., et al. (2024). Single-cell RNA sequencing illuminates the ontogeny, conservation and diversification of cartilaginous and bony fish lymphocytes. Nat. Commun. 15, 7627.

69. Krumsiek, J., Marr, C., Schroeder, T., and Theis, F.J. (2011). Hierarchical differentiation of myeloid progenitors is encoded in the transcription factor network. PLoS One 6, e22649.

70. Monticelli, S., and Natoli, G. (2017). Transcriptional determination and functional specificity of myeloid cells: making sense of diversity. Nat. Rev. Immunol. 17, 595–607.

71. Crow, M., Suresh, H., Lee, J., and Gillis, J. (2022). Coexpression reveals conserved gene programs that co-vary with cell type across kingdoms. Nucleic Acids Res. 50, 4302–4314.

72. Shi, Y., Strasser, A., Green, D.R., Latz, E., Mantovani, A., and Melino, G. (2024). Legacy of the discovery of the T-cell receptor: 40 years of shaping basic immunology and translational work to develop novel therapies. Cell. Mol. Immunol. 21, 790–797.

73. Austyn, J.M. (2016). Dendritic cells in the immune system-History, Lineages, tissues, tolerance, and Immunity. Microbiol. Spectr. 4. 10.1128/microbiolspec.MCHD-0046-2016.

74. Söderhäll, K. (2024). Invertebrate immunology - some thoughts about past and future research. Dev. Comp. Immunol. 161, 105256.

75. Sasaki, N., Ogasawara, M., Sekiguchi, T., Kusumoto, S., and Satake, H. (2009). Toll-like receptors of the ascidian Ciona intestinalis: prototypes with hybrid functionalities of vertebrate Toll-like receptors. J. Biol. Chem. 284, 27336–27343.

76. Brunetti, R., Gissi, C., Pennati, R., Caicci, F., Gasparini, F., and Manni, L. (2015). Morphological evidence that the molecularly determined Ciona intestinalis type A and type B are different species: Ciona robusta and Ciona intestinalis. J. Zoolog. Syst. Evol. Res. 53, 186–193.

77. Satou, Y., Tokuoka, M., Oda-Ishii, I., Tokuhiro, S., Ishida, T., Liu, B., and Iwamura, Y. (2022). A Manually Curated Gene Model Set for an Ascidian, Ciona robusta (Ciona intestinalis Type A). jzoo 39. 10.2108/zs210102.

78. Satou, Y., Nakamura, R., Yu, D., Yoshida, R., Hamada, M., Fujie, M., Hisata, K., Takeda, H., and Satoh, N. (2019). A Nearly Complete Genome of Ciona intestinalis Type A (C. robusta) Reveals the Contribution of Inversion to Chromosomal Evolution in the Genus Ciona. Genome Biol. Evol. 11, 3144–3157.

79. Satou, Y., Kawashima, T., Shoguchi, E., Nakayama, A., and Satoh, N. (2005). An integrated database of the ascidian, Ciona intestinalis: towards functional genomics. Zoolog. Sci. 22, 837–843.

80. Cunningham, F., Allen, J.E., Allen, J., Alvarez-Jarreta, J., Amode, M.R., Armean, I.M., Austine-Orimoloye, O., Azov, A.G., Barnes, I., Bennett, R., et al. (2022). Ensembl 2022. Nucleic Acids Res. 50, D988–D995.

81. Pertea, G., and Pertea, M. (2020). GFF Utilities: GffRead and GffCompare. F1000Res. 9. 10.12688/f1000research.23297.2.

82. Wolock, S.L., Lopez, R., and Klein, A.M. (2019). Scrublet: Computational Identification of Cell Doublets in Single-Cell Transcriptomic Data. Cell Syst 8, 281–291.e9.

83. Wolf, F.A., Angerer, P., and Theis, F.J. (2018). SCANPY: large-scale single-cell gene expression data analysis. Genome Biol. 19, 15.

84. Klein, A.M., Mazutis, L., Akartuna, I., Tallapragada, N., Veres, A., Li, V., Peshkin, L., Weitz, D.A., and Kirschner, M.W. (2015). Droplet barcoding for single-cell transcriptomics applied to embryonic stem cells. Cell 161, 1187–1201.

85. Polański, K., Young, M.D., Miao, Z., Meyer, K.B., Teichmann, S.A., and Park, J.-E. (2020). BBKNN: fast batch alignment of single cell transcriptomes. Bioinformatics 36, 964–965.

86. Cao, C., Lemaire, L.A., Wang, W., Yoon, P.H., Choi, Y.A., Parsons, L.R., Matese, J.C., Wang, W., Levine, M., and Chen, K. (2019). Comprehensive single-cell transcriptome lineages of a proto-vertebrate. Nature 571, 349–354.

87. Choi, H.M.T., Beck, V.A., and Pierce, N.A. (2014). Next-generation in situ hybridization chain reaction: higher gain, lower cost, greater durability. ACS Nano 8, 4284–4294.

88. Bruce, H.S., Jerz, G., Kelly, S., McCarthy, J., Pomerantz, A., Senevirathne, G., Sherrard, A., Sun, D.A., Wolff, C., and Patel, N.H. (2021). Hybridization Chain Reaction (HCR) In Situ Protocol.

89. Sofroniew, N., Lambert, T., Bokota, G., Nunez-Iglesias, J., Sobolewski, P., Sweet, A., Gaifas, L., Evans, K., Burt, A., Doncila Pop, D., et al. (2025). napari: a multi-dimensional image viewer for Python (Zenodo) 10.5281/ZENODO.15314358.

90. Davidson, B., Shi, W., and Levine, M. (2005). Uncoupling heart cell specification and migration in the simple chordate Ciona intestinalis. Development 132, 4811–4818.

91. Davidson, B., Shi, W., Beh, J., Christiaen, L., and Levine, M. (2006). FGF signaling delineates the cardiac progenitor field in the simple chordate, Ciona intestinalis. Genes Dev. 20, 2728–2738.

92. Zeller, R.W., Virata, M.J., and Cone, A.C. (2006). Predictable mosaic transgene expression in ascidian embryos produced with a simple electroporation device. Dev. Dyn. 235, 1921–1932.

93. Bergen, V., Lange, M., Peidli, S., Wolf, F.A., and Theis, F.J. (2020). Generalizing RNA velocity to transient cell states through dynamical modeling. Nat. Biotechnol. 38, 1408–1414.

94. Hagberg, A., Swart, P.J., and Schult, D.A. (2008). Exploring network structure, dynamics, and function using NetworkX (Los Alamos National Laboratory (LANL), Los Alamos, NM (United States)).

95. Gansner, E.R., and North, S.C. (2000). An open graph visualization system and its applications to software engineering. Softw. Pract. Exp. 30, 1203–1233.

96. Harvath, L., and Terle, D.A. (1999). Assay for phagocytosis. Methods Mol. Biol. 115, 281–290.

97. Camacho, C., Coulouris, G., Avagyan, V., Ma, N., Papadopoulos, J., Bealer, K., and Madden, T.L. (2009). BLAST+: architecture and applications. BMC Bioinformatics 10, 421.

98. Otto, E., Culakova, E., Meng, S., Zhang, Z., Xu, H., Mohile, S., and Flannery, M.A. (2022). Overview of Sankey flow diagrams: Focusing on symptom trajectories in older adults with advanced cancer. J. Geriatr. Oncol. 13, 742–746.

99. Lambert, S.A., Jolma, A., Campitelli, L.F., Das, P.K., Yin, Y., Albu, M., Chen, X., Taipale, J., Hughes, T.R., and Weirauch, M.T. (2018). The human transcription factors. Cell 172, 650–665.

100. Blanchoud, S., Rinkevich, B., and Wilson, M.J. (2018). Whole-Body Regeneration in the Colonial Tunicate Botrylloides leachii. Results Probl. Cell Differ. 65, 337–355.

101. Tassy, O., Dauga, D., Daian, F., Sobral, D., Robin, F., Khoueiry, P., Salgado, D., Fox, V., Caillol, D., Schiappa, R., et al. (2010). The ANISEED database: digital representation, formalization, and elucidation of a chordate developmental program. Genome Res. 20, 1459–1468.

102. Seal, R.L., Braschi, B., Gray, K., Jones, T.E.M., Tweedie, S., Haim-Vilmovsky, L., and Bruford, E.A. (2023). Genenames.Org: The HGNC resources in 2023. Nucleic Acids Res. 51, D1003–D1009.

103. Kaur, D., Patiyal, S., Sharma, N., Usmani, S.S., and Raghava, G.P.S. (2019). PRRDB 2.0: a comprehensive database of pattern-recognition receptors and their ligands. Database (Oxford) 2019, baz076.

104. Hanrahan, A.J., Iyer, G., and Solit, D.B. (2020). Intracellular Signaling. In Abeloff’s Clinical Oncology (Elsevier), pp. 24–46.e12.

